# Evolutionary signatures of self-incompatibility genes in grasses: high diversity, trans-generic polymorphism and co-evolution

**DOI:** 10.64898/2026.10.02.754995

**Authors:** Daniela Kupper, Marius Schmid, Steven Yates, Yutang Chen, Chloé Manzanares, Bruno Studer

## Abstract

In many flowering plants, self-incompatibility (SI) is a genetically controlled recognition system that prevents self-fertilization, thereby promoting outcrossing and maintaining genetic diversity within populations. The gametophytic SI system in grasses (Poaceae) is governed by two multiallelic loci, *S* and *Z*, each locus comprising of three genes governing the initial recognition between pollen and stigma. Identifying these genes establishes the molecular basis of SI in grasses, but how its unique two-locus architecture has shaped the evolution of the underlying loci and recognition genes remains largely unexplored. Here, we report targeted sequence capture of SI genes in 362 genotypes across nine forage grass species, revealing extensive allelic richness and sequence diversity. The SI genes exhibited elevated nucleotide diversity (π) (0.134 < π < 0.403) and great amino acid sequence divergence, ranging between 48.2-58.0% for the female determinants (sS and sZ) and 66.5-83.4% for the male determinants (SI-DUF247). The SI-DUF247 proteins also displayed trans-generic polymorphism, indicating that alleles predate speciation. Phylogenetic analysis of SI determinants showed congruence at both *S*- and *Z*-locus, consistent with co-evolution of recognition partners. Our results demonstrate that the unique two-locus SI system in grasses shares evolutionary characteristics with other SI systems, suggesting that their reproductive function is shaping their evolution. These findings also indicate that the grass SI loci are under long-term balancing selection to maintain recognition specificity while creating sufficient allelic diversity. Our study provides comprehensive sequence resources for SI genes and new insights into the evolution of the unique two-locus SI system of grasses.

## Introduction

Self-incompatibility (SI) is defined as a genetic mechanism that prevents self-fertilization via pollen-pistil recognition (de Nettancourt 1977; Takayama and Isogai 2005), occurring in approximately 40% of flowering plants and across at least 100 families (Igic et al. 2008). The genetic controls of SI as well as the physiological mechanisms are diverse (Zhang et al. 2024). While all systems mediate the rejection of self-pollen, the underlying genes are unique to each system and are non-homologous, supporting the hypothesis that SI has evolved independently multiple times (Steinbachs and Holsinger 2002; Franklin-Tong and Franklin 2003). Despite arising from independent origins, SI is governed by molecular interactions of the multiallelic genes expressed in the pistil and pollen. In most systems, these genes are encoded on a single non-recombining locus, the *S*-locus (de Nettancourt 1977; Fujii et al. 2016).

The grass SI system, however, deviates from this canonical single locus architecture. Already in the 1950s, classical genetic studies revealed that the SI system of grasses is controlled by two independent loci, *S* and *Z* (Lundquist 1956; Hayman 1956), which is unique among SI systems. It was hypothesized that the second SI locus arose from a duplication (Lundquist 1962) and therefore, the gene composition at both loci was expected to be similar (Yang et al. 2008). Nevertheless, despite decades of research on these loci, it was only recently that the genes governing self-recognition have been identified (Manzanares et al. 2016; Lian et al. 2021; Herridge et al. 2022; Rohner et al. 2023). In forage grasses, each locus is comprised of three tightly linked genes: two *DOMAIN OF UNKNOWN FUNCTION* (*DUF*) *247* genes expressed in the pollen and one gene encoding a secreted peptide expressed in the stigma, *STIGMA S* (*sS*) and *STIGMA Z* (*sZ*), at the *S*- or *Z*-locus, respectively (Herridge et al. 2022; Rohner et al. 2023).

Despite the differences between genetic control and molecular mechanism of all the identified SI systems, many evolutionary features are common. The first key feature shared among SI mechanisms is the high allelic richness and the extensive amino acid divergence between protein variants of a SI determinant. Allelic richness is important for maintaining sufficient compatible mating partners and genetic diversity within partners. In *Lolium perenne* L., high allelic diversity has been observed and to date, 24 *LpSDUF247-I* alleles have been identified (Rohner et al. 2023). In *Papaver rhoeas* L., the stigmatic *S-*proteins exhibit 40-50% sequence divergence (Wheeler et al. 2009). Rohner et al. (2023) also reported a high protein sequence divergence, ranging from 33% (sZ) to 79% (SDUF247-I) in grasses. In nature, rare SI alleles are favoured because they will more frequently find compatible partners (Wright 1939). The allelic balance at the SI genes in a species is strongly linked to these rare alleles advantage, named negative frequency-dependent selection (Castric and Vekemans 2004).

A second key feature of SI systems, directly linked to this form of balancing selection, is the conservation of some alleles over the evolutionary path of a species. When these allelic lineages persist through speciation events, the resulting polymorphism can be shared among species, or even genera, and is referred to as trans-specific or trans-generic polymorphism, respectively. Trans-generic alleles have, for examples, been reported for the *S*-RNase system in Solanaceae (Savage and Miller 2006). In *Arabidopsis halleri* (L.) O’Kane & Al-Shehbaz and *A. lyrata* L., 53 and 58 *S*-alleles, respectively, have been reported, of which 43 are shared between the two species and, therefore, predate their divergence (Durand et al. 2020).

A third key feature is the bipartite structure of pollen and pistil SI genes, which implies that recombination between pollen and pistil determinants at the *S*-locus in a single-locus system, or at the *S-* and *Z*-locus in grasses, may disrupt the linkage of the genes and, hence, lead to the loss of SI. The evolution of these genes is tightly linked in order to maintain a functional mechanism (Takuno et al. 2007). Co-evolution between male (pollen expressed) and female (pistil expressed) SI determinants were observed in the Brassica SI system (Sato et al. 2002; Takuno et al. 2007) and in the *S-RNase*-based SI system of *Petunia* (Kubo et al. 2015). This tight dependence between male and female determinants is also evident in grasses. Although the grass SI system is governed by two loci, it was shown that if one SI determinant at one locus is absent or disrupted, self-pollination is possible (Rohner et al. 2023).

Lastly, SI determinants often exhibit an elevated rate of evolution because mutations leading to new SI specificities are expected to be favored (Durand et al. 2020). Castric and Vekemans (2007) reported twelve and 13 codons under positive selection in the female determinant of *A. halleri* and *A. lyrata*, respectively. Also in *Phlox*, six codons under positive selection were identified in the female SI gene (Burgin et al. 2026).

Key to studying the mechanisms and evolutionary signatures of well-characterized SI systems is the availability of large collections of SI alleles (Paape et al. 2011; Fukushima et al. 2021; Maenosono et al. 2024). For the two-locus SI system of grasses, however, the limited availability of allele sequences has constrained such studies. Although high-quality genomes are available for major grass crops such as rice (*Oryza sativa* L.) and wheat (*Triticum aestivum* L.) (Zhu et al. 2021; Guo et al. 2025), these species are self-compatible (loss of SI through evolution and breeding) and, therefore, provide limited insights into the SI loci (Rohner et al. 2023). In contrast, reference-level genome assemblies of self-incompatible forage grasses are limited, because of their complex genomes (Chen et al. 2025). Furthermore, due to the high level of sequence divergence of SI alleles in grasses, the sequencing of the SI genes using PCR amplification and Sanger sequencing is often not successful (Maenosono et al. 2024). The main goal of this study was to investigate how the unique two-locus recognition system of grasses has shaped the long-term evolution of its molecular determinants. Using a large collection of SI gene sequences across multiple forage grasses, we aimed at evaluating allelic diversity at the *S-* and *Z*-locus, examining the persistence of SI alleles across species and genera and assessing whether the SI determinants within each locus exhibit shared evolutionary histories. We further aimed to determine whether the resulting evolutionary signatures are specific to the grass SI system or represent broader features shared with other well-characterized SI systems.

## Results

### Targeted sequence capture of the *S-* and *Z-*locus

A total of 362 plants from nine species of the Poaceae family were included in a targeted sequence capture (TSC), conducted in two successive rounds, hereafter referred to as TSC1 and TSC2. Across the four *SI-DUF247* genes, 386 unique protein sequences were identified (Table 1). We also obtained 39 unique protein sequences for the female determinants sS and sZ. For the two flanking genes *PYRIDOXAL PHOSPHATE HOMEOSTASIS PROTEIN* (*PLP*) and *GLYCEROL KINASE* (*GK*), we obtained more unique nucleotide sequences than possible for a single gene and 362 plants. With a BLAST search of five haplotype-resolved assemblies, we identified duplicated copies of *GK* in one *Z*-locus haplotype and duplicated copies of *PLP* in three *S*-locus haplotypes (data not shown). Because the retained copies were highly similar (mean similarity for GK of 98.09%, σ = 0.99% and for PLP 97.35%, σ = 0.99%) and indistinguishable after filtering, they were treated as a single gene in downstream analyses.

**Table 1.**
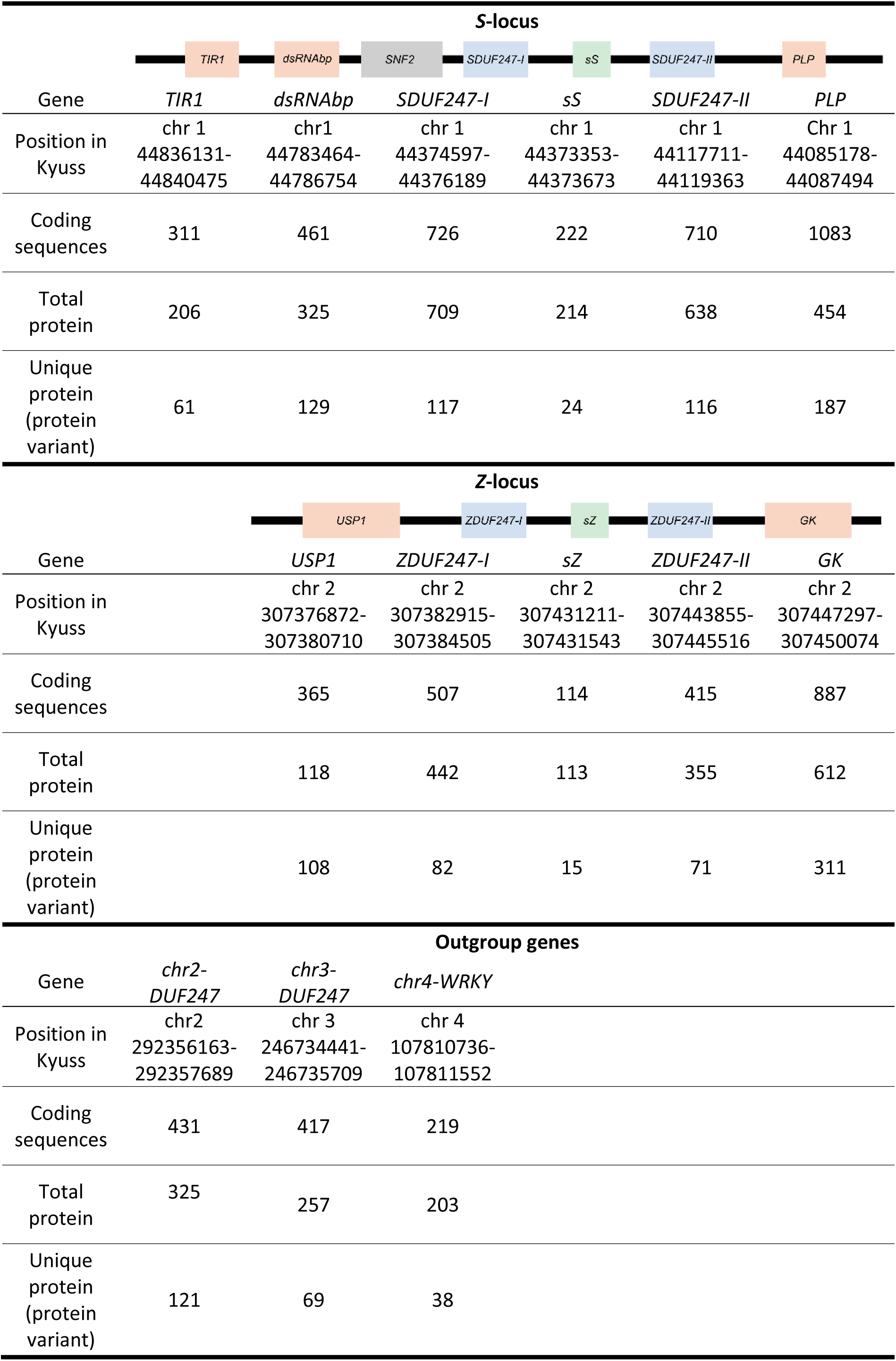
Description of the number of coding sequences and the corresponding total and unique protein sequences of the targeted genes. Flanking genes at the *S*-locus on chromosome (chr) 1 are *TRANSPORTER INHIBITOR RESPONSE 1-LIKE PROTEIN* (*TIR1*)*, DOUBLE-STRANDED RNA-BINDING PROTEIN 2* (*dsRNAbp*) and *PYRIDOXAL PHOSPHATE HOMEOSTASIS PROTEIN* (*PLP*). Flanking genes at the *Z*-locus on chr 2 are *UBIQUITIN CARBOXYL-TERMINAL HYDROLASE* (*USP1*) and *GLYCEROL KINASE* (*GK*). Outgroup genes are two genes containing a *DOMAIN OF UNKNOWN FUNCTION* (*DUF*) *247* on chr 2 and 3 (*chr2-DUF247, chr3-DUF247*) and *WRKY* on chr 4 (*chr4-WRKY*). Gene positions in the reference genome Kyuss v2.0 (Chen et al. 2024) are indicated. At the *S*-locus, *CHROMATIN-REMODELING COMPLEX SUBUNIT* (*SNF2*) was not included in the analysis.

As a quality control, the open reading frames (ORFs) of the SI genes from three *L. perenne* genomes and one *Lolium multiflorum* Lam. genome assemblies (Frei et al. 2021; Copetti et al. 2021; Rohner et al. 2023 and in-house data) were compared to the *de novo* assemblies of the same genotypes in the TSC2 dataset (Table S1). No mismatches were found within the ORFs, and only minor nucleotide differences (> 99% identity) occurred upstream or downstream of the ORF. These results confirmed the accuracy and reliability of the sequence capture and the automated assembly in reconstructing *SI-DUF247* contigs.

Another level of validation was the number of alleles found per gene and per genotype. Because the here presented sequences were derived from breeding material, we compared the pedigree with the alleles found in our data set. The population F13 was derived from a half-sib family containing seven parents in total (Thorogood et al. 2017). Therefore, it was expected to find a maximum of 14 alleles per gene. Indeed, we found only eleven alleles for *SDUF247-I*, ten alleles for *SDUF247-II* and two and three alleles for *ZDUF247-I* and *ZDUF247-II*, respectively. Moreover, genotype S23 Z, which is one of the parental plants of this family, shared one allele for SDUF247-I and SDUF247-II with genotypes from the F13 population.

The number of unique nucleotide sequences (alleles) and unique protein sequences (protein variants) varied between the different genes. Overall, flanking genes have more unique protein variants (n = 61-187) than the SI genes (n = 71-117 for the SI-DUF247) (Table 1). The position and number of nucleotide variation within a gene sequence is also important. In the case of the flanking genes, the many unique protein variants often resulted from variation in few nucleotides (see Table 1). The amino acid sequences of the flanking genes also showed high similarity, with mean identities ranging from 96.24% (σ = 5.07%) for dsRNAbp to 98.96% (σ = 1.82%) for TIR1. Due to the lower sequence diversity of the flanking genes, it was also easier to capture more of the alleles in the collection.

### Amino acid conservation, sequence diversity and interspecific relationships

When examining the conservation of amino acids (based on all-vs-all BLAST), the four DUF247 proteins clearly separated into four groups (Figure S1). Within each protein, the mean similarity ranged from 66.5% for ZDUF247-II (σ = 9.9%), 72.5% for ZDUF247-I (σ = 6.8%) to 83.4% for both SDUF247 (SDUF247-I: σ = 3.6%; SDUF247-II: σ = 3.9%). However, the mean similarity between groups was similar, ranging from 47.6% (σ = 1.7%) to 53.3% (σ = 1.8%) for SDUF247-II, ZDUF247-I and ZDUF247-II. The exception was SDUF247-I, which was less similar (44.7%, σ = 1.1% to 48.3%, σ = 1.5%) to all three other SI-DUF247 proteins. This indicates that whilst ZDUF247-I and ZDUF247-II are physically located at the same locus, they are roughly as similar to each other as they are to SDUF247-I and SDUF247-II, which are located on a different chromosome. The mean similarity within proteins for the female determinants was 58.0% (σ = 12.7%) for sS and 48.2% (σ = 18.9%) for sZ. The mean similarity between these two proteins was only 2.2% (σ = 9.6%) (Figure S1).

To assess sequence diversity and interspecific relationships, phylogenetic trees were constructed separately for each gene of the SI loci (the four *SI-DUF247* and the two female components *sS* and *sZ*) as well as the flanking genes and other genetically unrelated genes (see list in Table 1). A noticeable difference between all the phylogenetic trees was the phylogenetic scale and the branch length. The branch lengths of the *Z*-locus trees were approximately three times greater than those in the *S*-locus trees. Furthermore, the *SI-DUF247* genes and female determinants *sS* and *sZ* were characterized by long branches, whereas the dsRNAbp and GK formed compact trees with comparatively short branches (Figure 1 and Figure 2, Figure S2).

**Figure 1.**
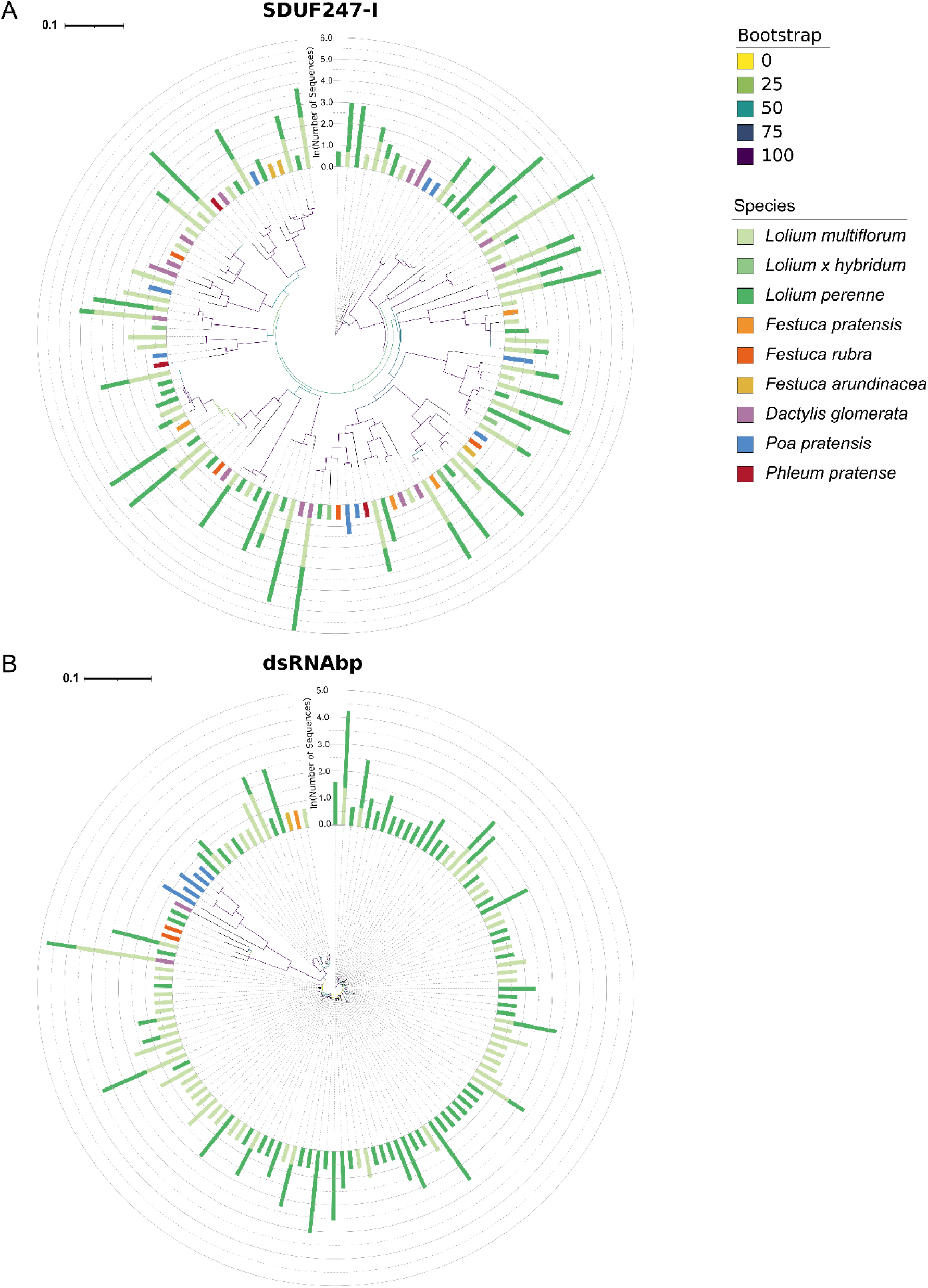
Maximum likelihood phylogenies of A) *SDUF247-I* and B) the *S*-locus flanking gene *dsRNAbp*. Branch lengths represent the number of amino acid substitutions per site, and branch colors indicate bootstrap support. Bar colors refer to the species in which the variant was found. The bar height is proportional to the natural logarithm of the number of sequences represented in this data set.

**Figure 2.**
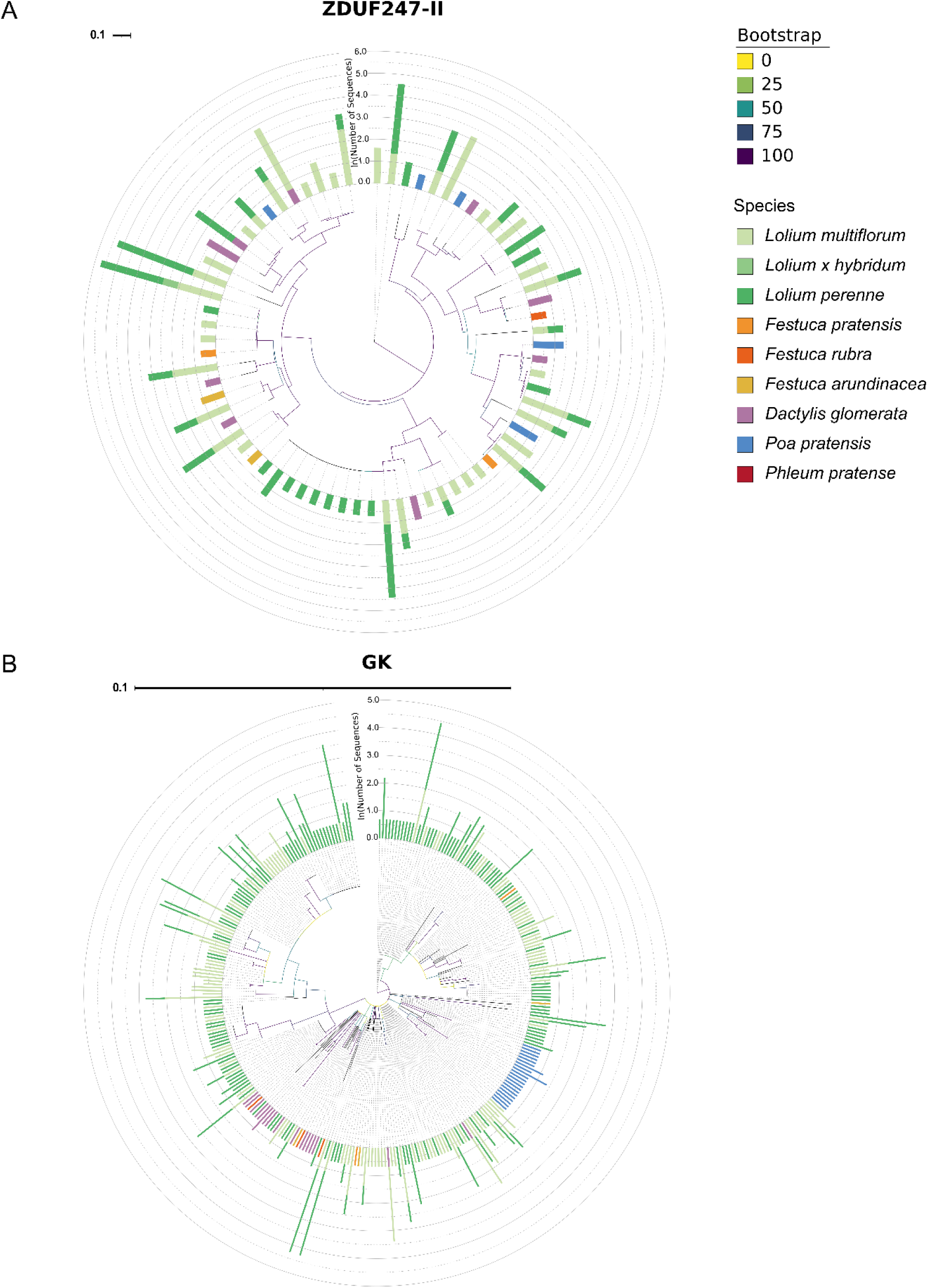
Maximum likelihood phylogenies of A) *ZDUF247-II* and B) the *Z*-locus flanking gene *GK.* Branch lengths represent the number of amino acid substitutions per site, and branch colors indicate bootstrap support. Bar colors refer to the species in which the variant was found. The bar height is proportional to the natural logarithm of the number of sequences represented in this data set.

Depending on the gene, the phylogenetic trees showed different grouping (Figure 1). For the SI-DUF247, the protein variants grouped independently of the species of origin. Indeed, in the case of SDUF247-I, a clade of the tree can group *Poa pratensis* L., *Phleum pratense* L., *Festuca pratensis* Huds., *L. multiflorum* and *L. perenne* (Figure 1A). A more concrete example is that for SDUF247-I, *Dactylis glomerata* L. shares the same unique protein sequence as *L. perenne* and *L. multiflorum,* while for SDUF247-II, the same protein variant is shared between *L. perenne*, *L. multiflorum* and *F. pratensis*. This grouping, independent of species, was similar for all the SI-DUF247, as well as for sS and sZ (Figure 1, Figure 2, Figure S2 and Figure S3). In contrast, protein variants of the flanking genes or even the outgroup DUF247 tended to group by the species as it can be seen for dsRNAbp and GK in Figure 1 and Figure 2.

### Topology of recognition partners

To study the co-evolution of the SI protein variants, we compared the topologies of the phylogenetic trees at the *S*- and *Z*-locus and assessed their congruence. For each SI locus, we grouped the variants based on 100% protein similarity, independently of the species. Only eleven to 13 protein variant groups were selected to simplify the readability of the tree (Figure 3). The tanglegrams of the *Z*-locus showed congruent branching pattern between the two *ZDUF247 g*enes, indicating co-evolution (Figure 3B). Less phylogenetic similarity was observed for the female determinant sZ, but the variant groups mostly retained corresponding phylogenetic groupings, supporting long-term evolutionary association between the male and female genes at *Z*. As for the *S*-locus, a similar pattern was observed, where the variant groups were staying associated with the same groups and clades independently of the genes.

**Figure 3.**
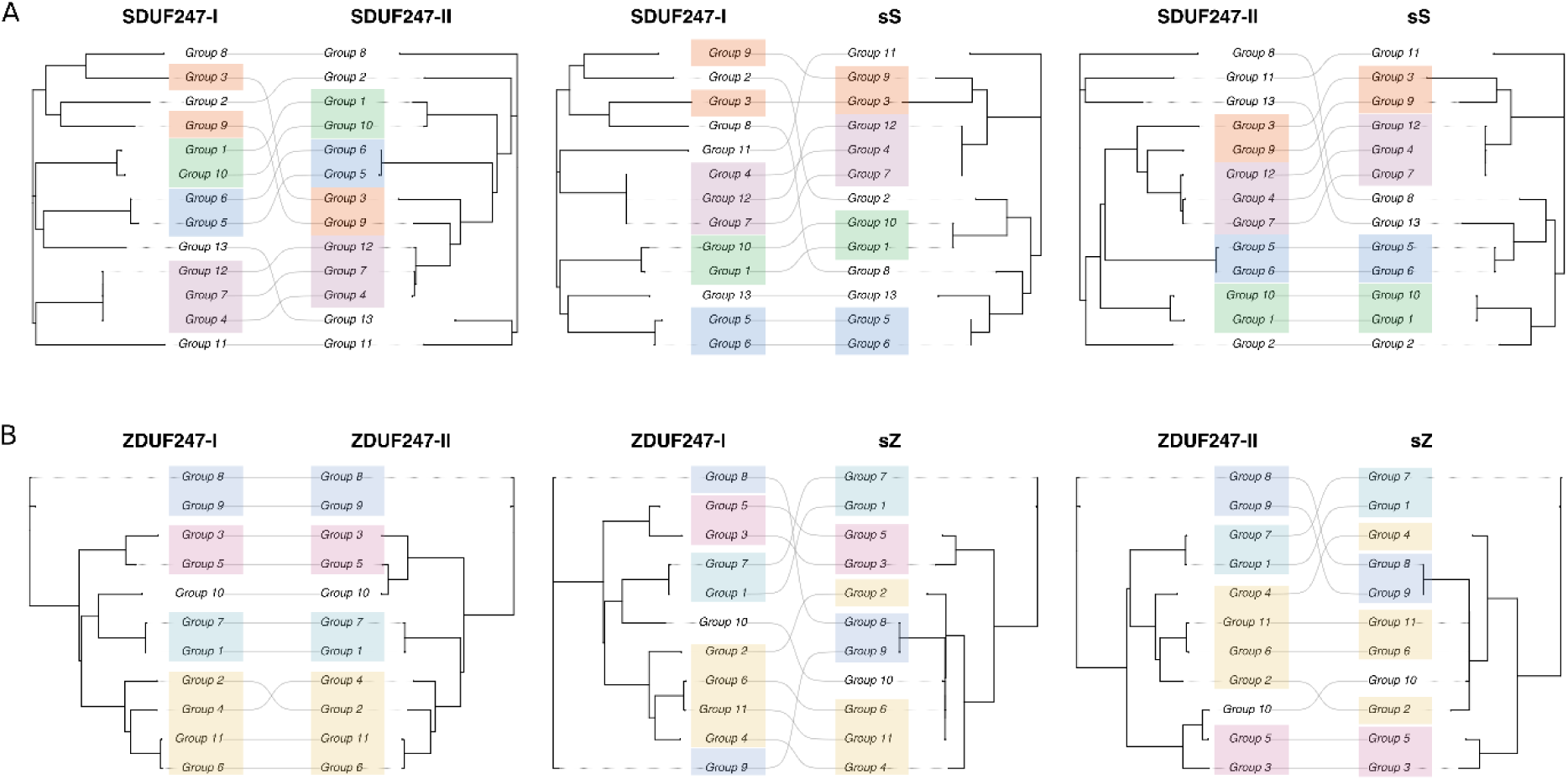
Tanglegram of A) *S*-locus genes *SDUF247-I, SDUF247-II* and *sS*; and B) of *Z*-locus genes *ZDUF247-I, ZDUF247-II* and *sZ*. At each locus, phylogenetic trees were pruned to protein variants representing multiple identical genotypes (= group). Groups that occur in proximity in at least two trees are highlighted with the same color.

### Positive selection at the SI loci

To test whether allelic variation in the SI proteins reflects positive selection, we analysed each SI gene, their flanking genes and the outgroup genes by calculating the nucleotide diversity (π) and applying codon-substitution models implemented in codeML. The nucleotide diversity was more than tenfold higher in the SI genes, ranging from π = 0.134 (*SDUF247-I*) to 0.403 (*sZ*), than in the flanking genes, where π ranged from 0.013 for *TIR1* to 0.046 for *GK* (Table 2). Interestingly, nucleotide diversity was twice as high for the *Z*-locus SI genes compared to the *S*-locus SI genes. When looking at the genes unlinked to SI, the nucleotide diversity in the *WRKY* gene (π = 0.037) was in a similar range as for the flanking genes (π ranging from 0.013 to 0.037), while nucleotide diversity in the outgroup *DUF247* was higher (π = 0.088). Furthermore, the intraspecific nucleotide diversity of the SI genes was also high across the different species, while the intraspecific nucleotide diversity of the other genes was generally lower (Table S3).

**Table 2.** Nucleotide diversity (π) and results of the codon substitution analysis of the self-incompatibility genes (*SDUF247-I/II, ZDUF247-I/II, sS* and *sZ*), flanking genes (*TIR1, dsRNAbp, PLP, USP1 and GK*) and outgroup genes (*chr2-DUF247, chr3-DUF247, chr4-WRKY*). The ratio of non-synonymous to synonymous substitution sites (ω) was calculated for each gene under model M8 and the number of sites with ω > 0.8 and Pr(ω > 1) ≥ 0.95 are shown.

|  |  |  | Codon substitution analysis |  |  |
| --- | --- | --- | --- | --- | --- |
|  |  |  | Length (codons) | Number of sites | Number of sites |
| | | | | $\omega > 0.8$ | $\text{Pr}(\omega > 1) \geq 0.95$ |
| S-locus | *TIR1* | 0.013 | 486 | 2 | 1 |
|  | *dsRNAbp* | 0.023 | 487 | 57 | 4 |
|  | *SDUF247-I* | 0.134 | 458 | 32 | 2 |
|  | *sS* | 0.291 | 65 | 5 | 0 |
|  | *SDUF247-II* | 0.149 | 407 | 21 | 0 |
|  | *PLP* | 0.029 | 222 | 5 | 1 |
| Z-locus | *USP1* | 0.026 | 459 | 11 | 7 |
|  | *ZDUF247-I* | 0.265 | 414 | 2 | 0 |
|  | *sZ* | 0.403 | 36 | 3 | 0 |
|  | *ZDUF247-II* | 0.281 | 380 | 5 | 0 |
|  | *GK* | 0.046 | 306 | 5 | 1 |
| Outgroup genes | *chr2-DUF247* | 0.088 | 348 | 17 | 1 |
|  | *chr3-DUF247* | 0.088 | 371 | 119 | 4 |
|  | *chr4-WRKY* | 0.037 | 60 | 12 | 0 |

**Table 3.** Summary of the plant material used for the targeted sequence capture assay. The species, the source of the plant material and the number of genotypes are displayed.

| Species | Source | Number of genotypes |
| --- | --- | --- |
| *Lolium perenne* | F13 population (Thorogood et al. 2017) | 54 |
| *Lolium perenne* | Diploid nursery population (Agroscope Reckenholz, Switzerland) | 43 |
| *Lolium perenne* | Tetraploid nursery population (Agroscope Reckenholz, Switzerland) | 13 |
| *Lolium perenne* | F2 self-compatible (SC) population (Cropano et al. 2021) | 9 |
| *Lolium perenne* | F2 biomass population (Anhalt et al. 2008) | 6 |
| *Lolium perenne* | Breeding program at DSV (Zaden, The Netherlands) | 2 |
| *Lolium perenne* | M6115 population (Agroscope Reckenholz, Switzerland) | 3 |
| *Lolium perenne* | Multiple varieties (Agroscope Reckenholz, Switzerland) | 5 |
| *Lolium perenne* | Genotype used in SI research or tissue culture (ETH Zurich, Switzerland) | 6 |
| *Lolium perenne* | Perennial ryegrass four-parents polycross (Agroscope Reckenholz, Switzerland) | 51 |
| *Lolium perenne* | Perennial ryegrass five-parents polycross (Agroscope Reckenholz, Switzerland) | 13 |
| *Lolium perenne* | Genotypes of a commercial perennial ryegrass varieties (Arara A and Arara B) (Agroscope Reckenholz, Switzerland) | 2 |
| *Lolium perenne* | Perennial ryegrass breeding material (Agroscope Reckenholz, Switzerland) | 3 |
| *Lolium perenne* | Genotypes used in self-incompatibility (SI) research (Kopecký et al. 2009; Manzanares et al. 2016; Rohner et al. 2023) | 6 |
| *Lolium perenne* | M34 and M35 population (Cropano et al. 2021) | 6 |
| *Lolium perenne* | F2 SC population (Agroscope Reckenholz, Switzerland) | 6 |
| *Lolium perenne* | Genotype involved in cytoplasmic male sterility (Vogt et al. 2020) | 3 |
| *Lolium perenne* and<br>*Lolium multiflorum* | Sequenced genotypes (Copetti et al. 2021; Frei et al. 2021; Rohner et al. 2023; Nagy et al. 2022, & DLF, Denmark) | 5 |
| *Lolium multiflorum* | Parents of the NAM population (Kiesbauer et al. 2026) | 49 |
| *Lolium multiflorum* | Italian ryegrass breeding material (Agroscope Reckenholz, Switzerland) | 41 |
| *Lolium multiflorum* | Genotypes with bacterial wilt resistance (Goettelmann et al. 2023) | 13 |
| *Dactylis glomerata* var.<br>*barexcel* | In-house grass reference material (ETH Zurich, Switzerland) | 1 |
| *Dactylis glomerata* | Orchard grass breeding material (Agroscope Reckenholz, Switzerland) | 5 |
| *Poa pratensis* var.<br>*tommy* | In-house grass reference material (ETH Zurich, Switzerland) | 2 |
| *Poa pratensis* | Common meadow-grass breeding material (Agroscope Reckenholz, Switzerland) | 5 |
| *Phleum pratense* var.<br>*tiller* | In-house grass reference material (ETH Zurich, Switzerland) | 1 |
| *Festuca pratensis* var.<br>*pradel* | In-house grass reference material (ETH Zurich, Switzerland) | 2 |
| *Festuca arundinacea*<br>var. *dulcia* | In-house grass reference material (ETH Zurich, Switzerland) | 2 |
| *Festuca rubra* var. *echo* | In-house grass reference material (ETH Zurich, Switzerland) | 2 |
| *Lolium multiflorum* var.<br>*westerwoldicum* | In-house grass reference material (ETH Zurich, Switzerland) | 1 |
| *Lolium x hybridum* var.<br>*ocadia* | In-house grass reference material (ETH Zurich, Switzerland) | 1 |
| *Lolium multiflorum* var.<br>*italicum* | In-house grass reference material (ETH Zurich, Switzerland) | 1 |

Significant positive selection was detected in *SDUF247-I* gene, with two codons showing strong posterior probability Bayes Empirical Bayes (BEB) support (Pr(ω > 1) ≥ 0.95) (Figure 3). When considering codons evolving under near-neutral or relaxed selection (ω > 0.8), 32 codons were identified in *SDUF247-I* and 21 in *SDUF247-II*, whereas only two such codons were detected in *ZDUF247-I* and five in *ZDUF247-II* (Figure S4). In the flanking genes, significant site-specific positive selection was found in *TIR1, USP1* and GK in which ω for most codons was generally low and only a few codons were estimated to be under positive selection (Figure S5). When looking at the outgroup *DUF247*, it becomes apparent that these genes are under positive selection. For examples, 14.57% of the sites for *DUF247* on chromosome 3 have an ω > 1 (Figure S4, Table S2).

To further investigate the spatial distribution of positively selected amino acid sites, we modelled a homo-hexamer of the SI-DUF247 as proposed by Lin et al. (2026) with AlphaFold and mapped the results of model M8 onto the predicted protein. All but one site under positive and relaxed selection were located in the non-cytoplasmic domain for all the SI-DUF247 proteins (Figure 4 and Figure S4). The exception is that for SDUF247-II, one site under relaxed selection was mapped on the trans-membrane domain (Figure S4). Interestingly, the two sites under positive selection in SDUF247-I were surface-exposed. These two residues appear in close spatial proximity to residues under near-neutral and relaxed selection. Furthermore, in the other SI-DUF247, residues under neutral and relaxed selection were often on the surface of the protein or in close proximity to neighboring subunit.

**Figure 4.**
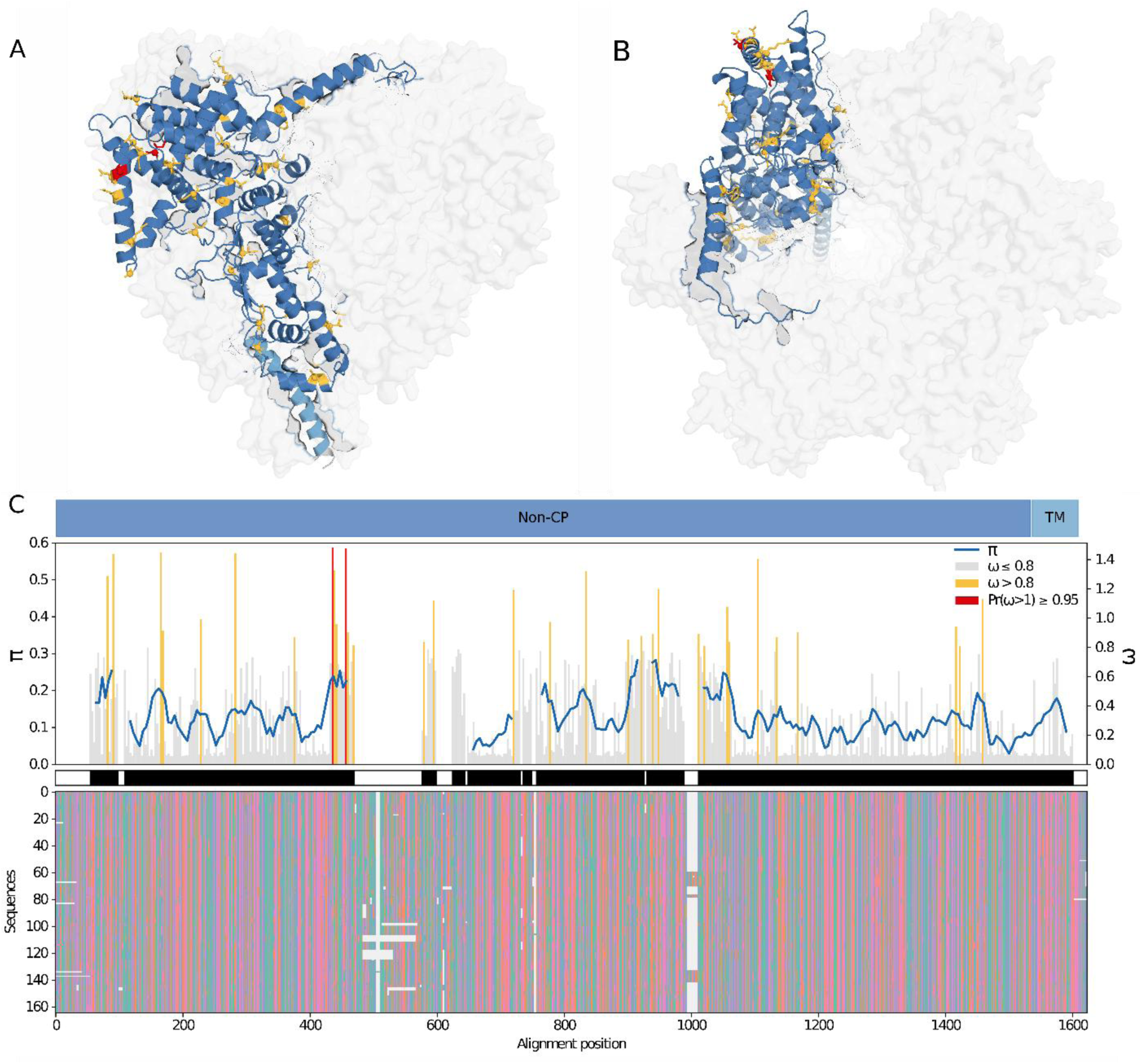
AlphaFold prediction of SDUF247-I, A) from the side, and B) the top. C) From bottom to top: Nucleotide sequence alignment of SDUF247-I, retained GBlocks are indicated in black. Nucleotide diversity (π), calculated with a sliding window approach across the well-aligned regions, is indicated in blue. The ratio of non-synonymous to synonymous substitution site (ω) across codons is indicated in; grey: purifying selection (mean ω < 0.8), yellow: neutral selection (mean ω > 0.8 but Pr(ω > 1) < 0.95), red: positive selection (Pr(ω > 1) ≥ 0.95). Non-cytoplasmic (Non-CP) and transmembrane (TM) domains are indicated in dark and light blue. The ratio of non-synonymous to synonymous substitution site (ω) across codons was also mapped on the predicted protein in A and B.

## Discussion

This study provides the largest allelic collection of grass SI determinants to date and the foundation to investigate the evolution of this unique SI system. Using this comprehensive sequence resource, we show for the first time that SI determinants do share the key features found in other SI systems. We have demonstrated the retention of ancient alleles through speciation and corroborate that the two-locus system results from a duplication of the *Z*-locus into the *S*-locus.

### High allelic diversity among the SI determinants despite originating from a duplication

The high sequence diversity of the SI determinants is known in the grass SI but also in other SI systems. The number of unique proteins captured for the female determinants sS and sZ (n = 39) is lower than the one for the SI-DUF247 (n = 386). This was expected, as we had less nucleotide sequences to design the targeted sequence capture assay, and these genes were only included in TSC2. The material sequenced in this study has been selected by breeders for various traits other than SI and, therefore, it is reasonable to assume that the allelic diversity of the SI genes in nature is greater than the one we describe here. However, the female components do show a large allelic diversity. The *SI-DUF247* genes prove to be as diverse between each other as within the same gene (allelic diversity). Based on the pairwise heatmap of all the SI-DUF247 proteins as well as the phylogenetic trees, we can differentiate all four SI-DUF247. In the phylogenetic tree from Sun et al. (2026), the *SI-DUF247* genes grouped according to their locus: *DUF247I-S* was clustered with *DUF247II-S* and *DUF247I-Z* clustered with *DUF247II-Z*. The branching in this tree suggests that the two-locus SI system in grass results from a duplication of one locus, the ancestral *Z*-locus, into the *S*-locus. This hypothesis that the two locus system arose from an ancestral duplication was already made by Lundquist (1962). From our individual phylogenetic trees, we can also notice the difference in the branch length between the *S-* and the *Z-* locus genes. This indicates that the *ZDUF247* genes have a higher divergence between their alleles, suggesting that they are older than the *SDUF247* genes. Our interpretation agrees with evolutionary analysis of Herridge et al. (2022), who proposed that the *Z*-locus represents the ancestral SI locus and that the *S*-locus arose through a subsequent duplication event. In *Leymus chinensis* (Trin.) Tzvel., Sun et al. (2026) also confirmed the duplication and dated *DUF247I/II-Z* at approximately 150 million years ago (MYA) and *DUF247I/II-S* approximately 50 MYA. When analysing the codon substitution, we only observed positive selection for SDUF247-I. However, the number of codons under relaxed selection in the SI-DUF247 proteins is also higher when we compare the SDUF247 to the ZDUF247 proteins. These patterns indicate that *SDUF247* genes experience substantially greater evolutionary rate heterogeneity and a relaxed constraint compared to the *ZDUF247* genes. This evolutionary observation is consistent with the duplication and an explanation for it could be that the *SDUF247* genes, which are younger, may have undergone accelerated diversification, creating new alleles to increase allelic diversity.

### Ancient SI alleles shared among different grass genera

Although our data do not allow us to date back the duplication of the *Z-*locus, we found with our large collection of alleles that the appearance of SI in grass is older than the speciation of many grasses. Several species like *D. glomerata*, *F. pratensis*, *L. perenne*, *L. multiflorum and L. hybridum* (hybrids between *L. multiflorum* and *L. perenne*) share common SI determinant alleles, with 100% amino acid similarity. The frequent clustering of *L. perenne*, *L. multiflorum* and *L. hybridum* sequences can be explained by the close evolutionary relationship among these species. *L. perenne* and *L. multiflorum* diverged only 2.58 MYA ago (Kumar et al. 2022), and have the ability to cross-fertilize. In contrast, it is estimated that the divergence between the *Lolium* species and *D. glomerata* date approximately 13.6 MYA and the divergence of *Lolium* and *Festuca* to approximately 3.12 MYA (Kumar et al. 2022). Furthermore, although the grasses have diverged, they have developed a species reproductive barrier, preventing most of them from cross-pollinating and therefore preventing the exchange of SI alleles. These trans-specific and trans-generic alleles that we are reporting, have been observed in other SI systems (Ioerger et al. 1990; Dwyer et al. 1991; Paape et al. 2011; Burgin et al. 2026) and are characteristics of negative-frequency-dependent selection, a form of balancing selection (Castric and Vekemans 2004). Additionally, we observed that the SI determinants exhibit a ten-times higher nucleotide diversity (π ranging from 0.134 for *SDUF247-I* up to 0.403 for *sZ*) compared to the flanking genes (0.013 < π < 0.046) and to the other unlinked genes studied (0.037 < π < 0.088). In *Arabidopsis*, a study measured the nucleotide diversity π of R genes to a maximum of 0.1 (π = 0.0047 on average for other genes) (Bakker et al. 2006). Such levels of diversity in the grass SI determinants are considerably higher than those reported for most plant genes and are comparable, to an extent, to the extreme diversity observed at plant loci subject to balancing selection (Castric and Vekemans 2004; Charlesworth 2006). Taken together, the observed trans-generic polymorphism and elevated nucleotide diversity support the hypothesis that balancing selection has shaped the evolutionary history of grass SI loci, maintaining ancient allelic lineages across both species and genus boundaries.

### Co-evolution of the male and female SI determinants

A common understanding of all SI systems is that there is a recognition between the male (pollen) and the female (stigma, pistil or even ovary) part of plant. This mechanism is often compared to a receptor-ligand interaction, like in disease resistance mechanisms, where the pathogen must be recognized by the host. Similarly, the interaction between recognition partners needs to be maintained to have a functional SI system. Evolutionary changes altering one recognition partner but not the other would indeed lead to a breakdown of SI (Fujii et al. 2016). In the Brassica SI system, highly similar topologies between the male (*S-LOCUS CYSTEINE-RICH PROTEIN SCR*) and the female SI determinants (*S-LOCUS RECEPTOR KINASE SRK*) were reported (Sato et al. 2002; Takuno et al. 2007), and it is suggested that they have co-evolved to preserve their specific interaction (Murase et al. 2020). Furthermore, in the *S-RNase*-based SI system of *Petunia*, the pollen-expressed *SLF* (*S-LOCUS F-BOX PROTEIN*) genes have co-evolved with *S*-RNase alleles from other haplotypes to maintain recognition of non-self *S*-RNases while avoiding recognition of the self *S*-RNase (Kubo et al. 2015; Bod’ová et al. 2018). In grass, we observed high congruence among the phylogenetic topologies of the SDUF247 as well as ZDUF247 proteins, demonstrating a co-evolution of the two male determinants at each locus. However, the congruence between the topology of male and the female determinants, independently of the SI locus, is reduced. This pattern may reflect an asymmetric diversification of SI genes, whereby the female determinant evolves more rapidly than its neighboring *SI-DUF247*, allowing for new allele formation. In *Arabidopsis*, such an asymmetric divergence model for SI receptor-ligands was proposed by Chantreau et al. (2019). Chantreau et al. (2019) reconstructed an ancestral SRK receptor from two distinct SRKs. The ancestral SRK recognized one SCR ligand but not the other, indicating that one ligand retained the ancestral specificity, whereas the second evolved a novel recognition specificity. Whether the emergence of new SI alleles in grasses also follows a similar asymmetric diversification process remains unknown. The emergence of new SI alleles can be different according to the SI system, and three main scenarios have been proposed by Charlesworth et al. (2005). To capture the evolutionary path of the grass SI genes, broader sampling of SI alleles in natural populations and across a broad set of species is required.

### Unique diversity among SI-DUF247

The DUF247 gene family is a family widely present in plants and can count up to 69 members in rice, spread over the genome (Zhang et al. 2024). In *L. perenne*, we have identified up to 63 genes containing a DUF247 domain (data not published). The high sequence diversity among the *SI-DUF247* genes is not seen in the other grass DUF247 family that we have sequenced. Zhang et al. (2024) assessed the evolution of all rice *DUF247* and concluded that for most of the genes in this family, strong purifying selection has happened. However, they did identify three tandem duplications that are under positive selection. The outgroup DUF247 genes we studied also showed many codons under positive selection, which could imply that they are involved in recognition mechanisms such as plant defense mechanisms. The DUF247 family has also been characterized as genes that have undergone duplication, whether in tandem or as blocks, like seen in Oak (*Quercus*) where the largest duplicated gene block is composed of members of the DUF247 family (Sork et al. 2022). Interestingly, we have observed on chromosome 7 of *L. perenne* Kyuss (Chen et al. 2024) two tandem repeats of a DUF247 gene (around 29 Mb for the first tandem and 186 Mb for the second). These observations were also made in rice by Zhang et al. (2024), where many tandem duplications on the same chromosome were observed. Another example of a DUF247 gene duplication is the *Y-SOFF* gene *(SUPPRESSOR OF FEMALE FUNCTION)*, mapped on the sex chromosome Y of Asparagus (*Asparagus officinalis* L.), which represses the development of female organ, therefore playing an important part in plant reproduction (Harkess et al. 2020). Bentz et al. (2026) have shown that this gene is resulting from a duplication of an ancient *DUF247* gene on chromosome 5 (not a sex chromosome) for ancestors of Asparagus, prior to dioecy in angiosperms. This new DUF247 gene then gained a new function (neofunctionalization) while being associated with the Y chromosome. The function of this DUF247 family is not well known but some DUF247 genes, like the *Y-SOFF* gene, have been characterized. In *Arabidopsis*, Kondou et al. (2013) showed that the over-expression of *DLE1 (DWARF AND LESION FORMATION 1)* (a member of the DUF247 family) activated some defense mechanism and concluded that it acts as a positive regulator of plant defense response. Another DUF247 gene in *Arabidopsis*, *DUF247-1*, was shown to be involved in the cell-wall polysaccharide composition, and to regulate other DUF247 gene expression (Wannitikul et al. 2023). In grasses, the SI-DUF247 are involved in the recognition of the female determinants and are predicted to be anchored in the cell membrane. Two mechanisms have been proposed by Rohner et al. (2023), depending on how the SI-DUF247 proteins interact between each other. It was also modeled by Lin et al. (2026) that SI-DUF247 proteins form a polymer, most likely a hexamer, that could resemble a channel. However, it is not known if the polymer is composed of only one SI-DUF247 protein (like proposed by Lin et al. (2026)), with both S-DUF247 proteins, or with all four SI-DUF247 (*S* and *Z* mixed). These various hypotheses remain to be experimentally proven to understand the grass SI mechanism at a physiological level.

We showed that the *SI-DUF247* genes are distinct from other *DUF247* genes in the grass genome in terms of allelic richness, amino acid divergence and trans-generic polymorphism. These findings further support their role as male SI determinants and therefore, we propose a new function-based nomenclature. *SDUF247-I* and *SDUF247-II* will be replaced by *SELF-INCOMPATIBILITY POLLEN S* 1 (*SIPS1*) and *SELF-INCOMPATIBILITY POLLEN S* 2 (*SIPS2*), respectively. Similarly, at the Z-locus, the male SI determinants will be named *SELF-INCOMPATIBILITY POLLEN Z 1* (*SIPZ1*) and *SELF-INCOMPATIBILITY POLLEN Z 2* (*SIPZ2*), replacing the current *ZDUF247-I* and *ZDUF247-II* names, respectively. To maintain consistency with nomenclature proposed for the male SI determinants, we further propose including the functional information also in the nomenclature of the female determinants. Accordingly, *sS* and *sZ* will be renamed *SELF-INCOMPATIBILITY STIGMA S (SISS)* and *SELF-INCOMPATIBILITY STIGMA Z (SISZ),* respectively. A consistent nomenclature in the community would facilitate their identification and comparison across different grasses. This will provide a framework for future evolutionary and functional studies with a larger collection of alleles, sequenced from broader grasses as well as ecotypes. By using prediction models to identify interacting amino acids or protein regions, it would enable us to better understand the possible interactions between the male and female components, as well as the *S-* and *Z*-locus. Combined with a transformation platform such as the one developed in *Arabidopsis* to study Papaver SI, the different models of interaction between the SI determinants could be studied and the evolution from one to two locus could be explored.

## Material and Methods

### Target sequence capture bait design and synthesis

The gene sequences of the four *SI-DUF247* (*SDUF247-I*, *SDUF247-II*, *ZDUF247-I* and *ZDUF247-II*), their flanking genes (*TIR1, dsRNAbp, PLP, USP1, GK*) and outgroup genes (*chr2-DUF247, chr3-DUF247*), were extracted from various genome assemblies (Mascher et al. 2013; Byrne et al. 2015; Velmurugan et al. 2016; Stein et al. 2018; Copetti et al. 2021; Nagy et al. 2022; Paril et al. 2022; Honig et al. 2017; Chen et al. 2025), an allele reconstruction approach (Manzanares et al. 2016), a BAC clone (Rohner et al. 2023) and sequence capture assays (Veeckman et al. 2019). In addition, the obtained sequences from TSC1 were used to design more baits for TSC2. More detailed information about the sequence acquisition can be found in Table S4.

The baits were designed with a length of 80 base pairs (bp) and with an overlap of 40 bp between adjacent sequences, resulting in an effective 2× coverage across the target regions. All baits were aligned to the *L. multiflorum* genome assembly (Copetti et al. 2021), and any baits matching regions annotated as repetitive elements were excluded. Redundant baits were further collapsed at 99% sequence identity. In TSC1, 18,483 unique baits designed and synthesized as a custom myBaits® kit (48 reactions, myBaits Custom Target Capture Kit with 1–20K unique probes; Arbor Biosciences, MI, USA). For TSC2, 33,452 unique baits were designed and synthesized (myBaits Custom Target Capture Kit with 20–40K unique probes; Arbor Biosciences). The resulting bait set is hereafter referred to as the “SI-baits”.

### Plant material sampling and plant DNA extraction

Leaf samples from 362 plants from the Poeae tribe (TSC1: n = 171, TSC2: n = 191) were collected (Table S4, Table S5). The plant material was flash-frozen in liquid nitrogen and stored in a freezer at -80 °C. Subsequently, the frozen samples were ground with a QIAGEN TissueLyser II (QIAGEN, Hilden, Germany). The genomic DNA was extracted from the pulverized samples using Mag-Bind® Plant DNA DS 96 Kit (Omega Bio-tek, GA, USA) combined with a KingFisher Flex Purification System (Thermo Fisher Scientific, Waltham, MA, USA) following the manufacturer’s protocol. The genomic DNA integrity was assessed with a 1% agarose gel, and the concentration and purity were quantified using a QIAxpert system (QIAGEN).

### Targeted sequence capture

A total of 250 ng for TSC1 and 500 ng for TSC2 of genomic DNA in 50 µl elution buffer (Omega Bio-tek, GA, USA) was fragmented with the Q800R2 Sonicator (Qsonica L.L.C, Newtown, CT, USA). The genomic DNA was fragmented for 150 seconds (TSC1) or 90 seconds (TSC2) with an amplitude of 20%, leading to a target fragment size of ∼500 bp (TSC1) and ∼420 bp (TSC2). The fragmentation was assessed for a subset of the 362 samples with a 2200 TapeStation using High-Sensitivity D1000 Screen Tape (Agilent Technologies, Santa Clara, CA, USA).

Dual-indexed libraries were prepared using the “NEBNext® Ultra™ II DNA Library Prep Kit for Illumina” combined with the “NEBNext® Multiplex Oligos for Illumina®” (Dual Index Primers Set 1 and Set 2, New England Biolabs, Ipswich, MA, USA). A size selection of 300 to 400 bp was performed during library preparation, and 12 PCR cycles were used for library enrichment. The concentration of the dual-indexed libraries was quantified using a Qubit fluorometer with the dsDNA quantification assay reagent (Thermo Fisher Scientific). Also, the library size distribution was quantified for a subset of libraries using a 2200 TapeStation using High-Sensitivity D1000 Screen Tape (Agilent Technologies). The observed average library sizes were approximately 620 bp and 540 bp for TSC1 and TSC2 libraries, respectively, which included for both the adapter sequences and the indexes.

Libraries were pooled according to the following criteria: similar concentration (±2.5 ng/µl), same species, and same ploidy level. For the TSC1 library, the dual-indexed libraries were divided into ten pools containing four libraries each, 29 pools containing four libraries each and two pools containing two libraries each, resulting in a total of 41 pools. For the TSC2 library, the dual-indexed libraries were distributed into 22 pools with five libraries each and seven pools containing six libraries each, resulting in a total of 39 pools.

Target enrichment of all pools was done using the SI-baits protocol “Hybridization Capture for Targeted NGS User Manual Version 5” (Arbor Biosciences). Hybridization and washing were carried out at 60 °C, and 13 cycles were performed to recover the enriched libraries.

The size distribution of the enriched libraries was quantified using a 2200 TapeStation with High-Sensitivity D1000 Screen Tape (Agilent Technologies). The concentration of the enriched libraries was quantified using a Qubit Fluorometer with dsDNA quantification assay reagent (Thermo Fisher Scientific). The molarity was determined based on the average size and concentration of each enriched library pool.

The 41 enriched libraries of TSC1 were normalized to 15 nM and pooled. To remove any residual adapter dimers and fragments below the size of 300 bp, a size selection step was performed with 0.7 × AMPure XP Beads (Beckman Coulter, Brea, CA, USA) following the manufacturer’s protocol. The size-selected pool had an average fragment size of 577 bp and was normalized to 4 nM. The fragments were sequenced using the Illumina MiSeq v3 (2 × 300 bp, 20-25 M reads, 13.2-15 Gb throughput) and the Illumina NovaSeq 6000 (2 × 150 bp, 1 SP Flowcell, 400 M reads).

The 39 enriched library pools of TSC2 were normalized to 10 nM and pooled. Subsequently, the fragments were sequenced twice using the Illumina NovaSeq 6000 (2 × 150 bp, 1 SP Flowcell, 400 M reads).

### *De novo* assembly of gene contigs

Raw reads were mapped to the target gene nucleotide sequences using the BBsplit function of the aligner tool BBmap (v. 39.1, Bushnell 2014). The gene and genotype-specific reads were then *de novo* assembled using SPAdes v3.15.4 with “careful” mode (--careful) (Bankevich et al. 2012), which minimizes mismatches and includes an additional error correction step using the MismatchCorrector tool.

Assembled contigs with a coverage below 10× were removed. For each gene, a minimal gene length was defined and used for filtering (e.g., below 200 bp for *sS* and *sZ* and below 1,000 bp for the *SI-DU247s*). Detailed filtering parameters for each gene can be found in Table S6. Next, the coding sequence (CDS) was predicted with Miniprot (v0.18-r281, Li 2023) using multiple reference protein sequences as queries. The best-supported gene model was then selected and translated to a protein sequence. Additional size filtering was then applied to the CDS and the protein sequence using SeqKit (’seq -m’, v2.13.0, Shen et al. 2024). For each gene, the filtering length was defined based on the length of the query sequence and optimized to reach a sufficient quality of the subsequent alignment (Table S6).

### Alignment and curation of coding sequences and protein sequences

The CDS and protein sequences were renamed according to the species and the genotype. Next, the CDS and the protein sequences were combined into a single fasta file and deduplicated (based on sequence) using SeqKit (’rmdup’, v2.13.0, Shen et al. 2024), resulting in nonredundant CDS and protein sequence datasets. The information on identical groups (duplicates) was stored in a separate file (duplicate list) to keep track of the species and the genotype represented by the sequence. The CDS alignment with MAFFT (Katoh & Standley, 2013), followed by extraction of conserved regions using Gblocks (Castresana, 2000), was conducted with the codon-alignment tool TranslatorX (v1.1, Abascal et al. 2010). Protein sequences were also aligned with MAFFT (v7.526, Katoh & Standley, 2013) and trimmed with Gblocks (v.0.91b, Castresana, 2000).

### Phylogeny and topology

Phylogenetic trees were constructed using iqtree3 (v3.1.2, Wong et al., 2025) and visualized with iTOL (v7.6, Letunic and Bork 2024). To investigate the topology of the SI determinants, 13 groups for the *S*-locus and 11 groups for the *Z*-locus were manually defined (Table S7). In short, for each tip in *sS* and *sZ* tree, the list of genotypes represented by that tip (duplicate list) was compared with the list of the *SI-DUF247* genes of the corresponding locus. When multiple genotypes overlapped in all three lists, the tree tip was retained in all three trees. The unrooted trees were pruned in R (v4.5.0, R Core Team 2025) using ape (v5.8.1, Paradis and Schliep 2019). Tanglegrams were drawn with the R package phytools (v2.5.2, Revell 2024).

### Pairwise comparison

All unique SI-DUF247 proteins were concatenated into a single fasta file and subjected to all-vs-all BLAST (v2.17.0, Altschul et al., 1990) in tabular format (-outfmt 6). These data were loaded in R and the percent similarity was plotted as a heatmap with dendrogram using the library pheatmap (v1.0.13, Kolde, 2025). Mean similarity within flaking genes was also calculated based on an all-vs-all BLAST (v2.17.0, Altschul et al., 1990).

### Nucleotide diversity

To evaluate the genetic diversity of the genes included in this study, we calculated the nucleotide diversity (π) as the mean proportion of pairwise nucleotide differences across aligned sequences (Nei and Li 1979). For each gene alignment, the overall nucleotide diversity was estimated across all aligned sequences. In addition, intraspecific nucleotide diversity was calculated separately for each species. Local variation in nucleotide diversity across the gene was further investigated using a sliding-window approach, with a window size of 20, and a step size of 5. Calculations were performed using a customized Python script, available at https://github.com/dakupper/Evo_SI-genes.

### Selection pressure

First, we constructed a phylogenetic tree based on the nucleotide sequence using RAxML (v8.2.12, Stamatakis 2014). Analyses were performed under the GTRGAMMA model, and branch support was assessed using 1,000 rapid bootstrap replicates followed by a search for the best-scoring maximum likelihood tree.

To assess patterns of molecular evolution across codons, we then applied codon-substitution models implemented using PAML (v4.10.10, Yang 2007). For each gene, we fitted M0, M1a, M2a, M7, and M8 models. M1a and M7 represent nearly neutral models, with ω variation among codons modeled using discrete site classes and a beta distribution, respectively (ω ≤ 1). M2a and M8 account for positive selection by adding an additional site class with ω > 1 to M1a and M7, respectively. We tested for positive selection using (LRT) between the nested model pairs M1a and M2a, and M7 and M8. LRT statistics (2ΔlnL) with two degrees of freedom were compared against the χ²-distribution (Yang 2007). M8 (β + ω > 1) model allows codons to evolve under a mixture of purifying selection (0 < ω < 1), neutral evolution (ω = 1), and positive selection (ω > 1). Posterior probabilities for each codon site were estimated using the BEB method of Yang et al. (2005). Codons were considered to be under statistically significant positive selection if Pr(ω > 1) ≥ 0.95, the conventional threshold for BEB support. Codons with elevated posterior mean ω values but lacking strong BEB support were interpreted as evolving under relaxed purifying selection rather than positive selection.

For visualization and interpretation of model M8, codons were classified into three evolutionary categories:

1. Purifying selection - mean ω < 0.8
2. Neutral or relaxed selection -mean ω > 0.8 but Pr(ω > 1) < 0.95
3. Positive selection - Pr(ω > 1) ≥ 0.95

This approach distinguishes sites with strong statistical evidence for adaptive evolution from those exhibiting non-significant but elevated nonsynonymous rates.

### AlphaFold visualization

According to Lin et al. (2026) a hexamer complex for each SI-DUF247 was modeled with AlphaFold3 on the AlphaFold Server (Abramson et al. 2024). In brief, homo-hexamer complexes were modeled with 12 Ca^2+^ ions. The best model was then visualized with PyMOL (v3.1.6.1, Schrödinger, LLC 2015). Predicted domains of the protein were extracted with InterPro (https://www.ebi.ac.uk/interpro/search/sequence/, accessed 18.08.2026). Coordinates of the positive and relaxed selection sites were also extracted and mapped on the protein.

### Code development and validation

Custom scripts were developed for the codon substitution analysis and calculations of nucleotide diversity as well as for data visualizations. The code was partially developed with the help of Microsoft Copilot (GPT 5.6), which was used to generate preliminary code suggestions and assist for debugging. All generated code was reviewed, tested and validated by the authors. The authors take responsibility for the final implementation and results.

## Acknowledgements

We thank Dr Christoph Grieder and Dr Michelle Nay (Agroscope), Dr Daniel Thorogood (IBERS, Aberystwyth University), Dr Thomas Lübberstedt (Iowa State University) as well as DSV Zaden Netherlands B.V. for providing us with plant material used in this study. We are grateful to Dr Miguel Lorea Sanchez (Molecular Plant Breeding, ETH Zurich) for helping to design the sequence capture experiment.

Sequencing with Illumina MiSeq was performed at the Genetic Diversity Centre (GDC), ETH Zurich, while the Illumina NovaSeq 6000 was conducted at the Functional Genomic Center Zurich (FGCZ). We thank Silvia Kobel, Dr Aria Minder Pfyl (GDC, ETH Zurich) and Ingrid Stoffel-Studer (Molecular Plant Breeding, ETH Zurich) for their assistance in the lab.

We further thank Dr Niklaus Zemp (GDC, ETH Zurich) for generating the *de novo* gene assemblies used in this study and for his assistance in planning the sequence capture experiment. We also want to thank Prof Dr Kentaro Shimizu and Dr Kazuki Fukushima (Evolutionary and Ecological Genomics, University of Zurich) for valuable discussions and sharing their expertise on the evolutionary and diversity related aspects of SI genes. We want to acknowledge Verena Knorst-Rashid (Molecular Plant Breeding, ETH Zurich) for her support in maintaining plant material.

This study was funded by the Swiss National Science Foundation (SNSF, grant number 197708).

## Authors contribution

M.S., C.M., S.Y. and B.S. conceived the study. M.S. prepared the samples. D.K., SY., Y.C., analyzed the data. D.K. prepared the visualizations. D.K. and C.M. wrote the manuscript with assistance from B.S. and S.Y. All authors read and approved the final manuscript.

## Data availability

The sequencing data and the *de novo* assemblies of the alleles generated in this study are available on NCBI: PRJNA1535929. Code used for the analysis of the *de novo* assembled genes is available at https://github.com/dakupper/Evo_SI-genes.

## References

Abascal, Federico, Rafael Zardoya, and Maximilian J. Telford. 2010. ‘TranslatorX: Multiple Alignment of Nucleotide Sequences Guided by Amino Acid Translations.’ Nucleic Acids Research (England) 38 (Web Server issue): W7–13. 10.1093/nar/gkq291.

Abramson, Josh, Jonas Adler, Jack Dunger, et al. 2024. ‘Accurate Structure Prediction of Biomolecular Interactions with AlphaFold 3’. Nature 630 (8016): 493–500. 10.1038/s41586-024-07487-w.

Altschul, Stephen F., Warren Gish, Webb Miller, Eugene W. Myers, and David J. Lipman. 1990. ‘Basic Local Alignment Search Tool’. Journal of Molecular Biology 215 (3): 403–10. 10.1016/S0022-2836(05)80360-2.

Anhalt, Ulrike C. M., Pat J. S. Heslop-Harrison, Stephen L. Byrne, Anthony Guillard, and Susanne Barth. 2008. ‘Segregation Distortion in Lolium: Evidence for Genetic Effects’. Theoretical and Applied Genetics 117 (2): 297–306. 10.1007/s00122-008-0774-7.

Bakker, Erica G., Christopher Toomajian, Martin Kreitman, and Joy Bergelson. 2006. ‘A Genome-Wide Survey of R Gene Polymorphisms in Arabidopsis.’ The Plant Cell (England) 18 (8): 1803–18. 10.1105/tpc.106.042614.

Bankevich, Anton, Sergey Nurk, Dmitry Antipov, et al. 2012. ‘SPAdes: A New Genome Assembly Algorithm and Its Applications to Single-Cell Sequencing’. Journal of Computational Biology 19 (5): 455–77. 10.1089/cmb.2012.0021.

Bentz, Philip C., Sarah B. Carey, Francesco Mercati, et al. 2026. ‘Two Independent Origins of XY Sex Chromosomes in Asparagus’. Molecular Biology and Evolution 43 (2): msag003. 10.1093/molbev/msag003.

Bod’ová, Katarína, Tadeas Priklopil, David L. Field, Nicholas H. Barton, and Melinda Pickup. 2018. ‘Evolutionary Pathways for the Generation of New Self-Incompatibility Haplotypes in a Nonself-Recognition System.’ Genetics (United States) 209 (3): 861–83. 10.1534/genetics.118.300748.

Burgin, Grace A., Nia Faith Lewis, and Robin Hopkins. 2026. ‘A Lectin Receptor-like Kinase Controls Self-Pollen Recognition in *Phlox*’. Proceedings of the National Academy of Sciences 123 (8): e2525299123. 10.1073/pnas.2525299123.

Bushnell, Brian. 2014. ‘BBMap: A Fast, Accurate, Splice-Aware Aligner’. In Conference: 9th Annual Genomics of Energy & Environment Meeting, Walnut Creek, CA.

Byrne, Stephen L., Istvan Nagy, Matthias Pfeifer, et al. 2015. ‘A Synteny-based Draft Genome Sequence of the Forage Grass *Lolium Perenne*’. The Plant Journal 84 (4): 816–26. 10.1111/tpj.13037.

Castresana, Jose. 2000. ‘Selection of Conserved Blocks from Multiple Alignments for Their Use in Phylogenetic Analysis’. Molecular Biology and Evolution 17 (4): 540–52. 10.1093/oxfordjournals.molbev.a026334.

Castric, Vincent, and Xavier Vekemans. 2004. ‘INVITED REVIEW: Plant Self-incompatibility in Natural Populations: A Critical Assessment of Recent Theoretical and Empirical Advances’. Molecular Ecology 13 (10): 2873–89. 10.1111/j.1365-294X.2004.02267.x.

Castric, Vincent, and Xavier Vekemans. 2007. ‘Evolution under Strong Balancing Selection: How Many Codons Determine Specificity at the Female Self-Incompatibility Gene SRK in Brassicaceae?’ BMC Evolutionary Biology 7 (1): 132. 10.1186/1471-2148-7-132.

Chantreau, Maxime, Céline Poux, Marc F. Lensink, Guillaume Brysbaert, Xavier Vekemans, and Vincent Castric. 2019. ‘Asymmetrical Diversification of the Receptor-Ligand Interaction Controlling Self-Incompatibility in Arabidopsis’. eLife 8 (November): e50253. 10.7554/eLife.50253.

Charlesworth, Deborah. 2006. ‘Balancing Selection and Its Effects on Sequences in Nearby Genome Regions’. PLOS Genetics 2 (4): e64. 10.1371/journal.pgen.0020064.

Charlesworth, Deborah, Xavier Vekemans, Vincent Castric, and Sylvain Glémin. 2005. ‘Plant Self-Incompatibility Systems: A Molecular Evolutionary Perspective’. New Phytologist 168 (1): 61–69. 10.1111/j.1469-8137.2005.01443.x.

Chen, Yutang, Dario Copetti, Roland Kölliker, et al. 2025. ‘Chromosome-Level Haplotype-Resolved Assembly of Highly Heterozygous Grass Genomes with PhaseGrass’. Nature Communications 17 (1): 12. 10.1038/s41467-025-66377-5.

Chen, Yutang, Roland Kölliker, Martin Mascher, et al. 2024. ‘An Improved Chromosome-Level Genome Assembly of Perennial Ryegrass (Lolium Perenne L.)’. Gigabyte 2024 (March): 1–11. 10.46471/gigabyte.112.

Copetti, Dario, Steven A. Yates, Maximilian M. Vogt, et al. 2021. *Evidence for High Intergenic Sequence Variation in Heterozygous Italian Ryegrass (* Lolium Multiflorum *Lam*.) Genome Revealed by a High-Quality Draft Diploid Genome Assembly. Preprint. Genomics. 10.1101/2021.05.05.442707.

Cropano, Claudio, Iain Place, Chloé Manzanares, et al. 2021. ‘Characterization and Practical Use of Self-Compatibility in Outcrossing Grass Species’. Annals of Botany 127: 841–52. 10.1093/aob/mcab043.

Durand, Eléonore, Maxime Chantreau, Audrey Le Veve, et al. 2020. ‘Evolution of Self-incompatibility in the Brassicaceae: Lessons from a Textbook Example of Natural Selection’. Evolutionary Applications 13 (6): 1279–97. 10.1111/eva.12933.

Dwyer, Kathy G., Michele A. Balent, June B. Nasrallah, and Mikhail E. Nasrallah. 1991. ‘DNA Sequences of Self-Incompatibility Genes from Brassica Campestris and B. Oleracea: Polymorphism Predating Speciation’. Plant Molecular Biology 16 (3): 481–86. 10.1007/BF00024000.

Franklin-Tong, Vernonica E., and F. Chris H. Franklin. 2003. ‘Gametophytic Self-Incompatibility Inhibits Pollen Tube Growth Using Different Mechanisms’. Trends in Plant Science 8 (12): 598–605. 10.1016/j.tplants.2003.10.008.

Frei, Daniel, Elisabeth Veekman, Daniel Grogg, et al. 2021. ‘Ultralong Oxford Nanopore Reads Enable the Development of a Reference-Grade Perennial Ryegrass Genome Assembly’. Genome Biology and Evolution 13 (8). 10.1093/GBE/EVAB159.

Fujii, Sota, Ken Ichi Kubo, and Seiji Takayama. 2016. ‘Non-Self- and Self-Recognition Models in Plant Self-Incompatibility’. Nature Plants 2 (9). 10.1038/nplants.2016.130.

Fukushima, Kazuki, Toko Kanomata, Aoi Kon, et al. 2021. ‘Spatiogenetic Characterization of *S* Receptor Kinase (*SRK*) Alleles in Naturalized Populations of *Raphanus Sativus* L. Var. *Raphanistroides* on Yakushima Island’. Genes & Genetic Systems 96 (3): 129–39. 10.1266/ggs.20-00066.

Goettelmann, Florian, Ralf Koebnik, Veronica Roman-Reyna, Bruno Studer, and Roland Kölliker. 2023. ‘High Genomic Plasticity and Unique Features of Xanthomonas Translucens Pv. Graminis Revealed through Comparative Analysis of Complete Genome Sequences.’ BMC Genomics (England) 24 (1): 741. 10.1186/s12864-023-09855-8.

Guo, Dongling, Yan Li, Hengyun Lu, et al. 2025. ‘A Pangenome Reference of Wild and Cultivated Rice’. Nature 642 (8068): 662–71. 10.1038/s41586-025-08883-6.

Harkess, Alex, Kun Huang, Ron van der Hulst, et al. 2020. ‘Sex Determination by Two Y-Linked Genes in Garden Asparagus.’ The Plant Cell (England) 32 (6): 1790–96. 10.1105/tpc.19.00859.

Hayman, David Lindsay. 1956. ‘The Genetical Control of Incompatibility in *Phalaris Coerulescens* Desf’. Australian Journal of Biological Sciences 9: 321–31. 10.1071/BI9560321.

Herridge, Rowan, Tyler McCourt, Jeanne M. E. Jacobs, Peter Mace, Lynette Brownfield, and Richard Macknight. 2022. ‘Identification of the Genes at *S* and *Z* Reveals the Molecular Basis and Evolution of Grass Self-Incompatibility’. Frontiers in Plant Science 13 (October): 1–15. 10.3389/fpls.2022.1011299.

Honig, Josh A., Ehud Zelzion, Nicole E. Wagner, et al. 2017. ‘Microsatellite Identification in Perennial Ryegrass Using Next-Generation Sequencing’. Crop Science 57 (S1): S–331. 10.2135/cropsci2016.07.0608.

Igic, Boris, Russell Lande, and Joshua R. Kohn. 2008. ‘Loss of Self-Incompatibility and Its Evolutionary Consequences’. International Journal of Plant Sciences 169 (1): 93–104. 10.1086/523362.

Ioerger, Thomas R., Andrew G. Clark, and Teh-Hui Kao. 1990. ‘Polymorphism at the Self-Incompatibility Locus in Solanaceae Predates Speciation.’ Proceedings of the National Academy of Sciences 87 (24): 9732–35. 10.1073/pnas.87.24.9732.

Katoh, Kazutaka, and Daron M. Standley. 2013. ‘MAFFT Multiple Sequence Alignment Software Version 7: Improvements in Performance and Usability’. Molecular Biology and Evolution 30 (4): 772–80. 10.1093/molbev/mst010.

Kiesbauer, Jenny, Christoph Grieder, Meril Sindelar, et al. 2026. ‘The First Nested Association Mapping (NAM) Population for Italian Ryegrass Reveals Genomic Regions Associated with Seed Shattering’. *BMC Plant Biology*, ahead of print, September 12. 10.1186/s12870-026-09936-2.

Kolde, Raivo. 2025. Pheatmap. V. 1.0.13. Released June 5. https://github.com/raivokolde/pheatmap.

Kondou, Youichi, Kosuke Noguchi, Shinsuke Kutsuna, et al. 2013. ‘Overexpression of DWARF AND LESION FORMATION 1 (DLE1) Causes Altered Activation of Plant Defense System in Arabidopsis Thaliana’. Plant Biotechnology 30 (4): 385–92. 10.5511/plantbiotechnology.13.0605a.

Kopecký, David, Jan Bartos, Adam J. Lukaszewski, et al. 2009. ‘Development and Mapping of DArT Markers within the Festuca - Lolium Complex.’ BMC Genomics (England) 10 (October): 473. 10.1186/1471-2164-10-473.

Kubo, Ken-Ichi, Timothy Paape, Masaomi Hatakeyama, et al. 2015. ‘Gene Duplication and Genetic Exchange Drive the Evolution of S-RNase-Based Self-Incompatibility in Petunia.’ Nature Plants (England) 1 (January): 14005. 10.1038/nplants.2014.5.

Kumar, Sudhir, Michael Suleski, Jack M. Craig, et al. 2022. ‘TimeTree 5: An Expanded Resource for Species Divergence Times.’ Molecular Biology and Evolution (United States) 39 (8). 10.1093/molbev/msac174.

Letunic, Ivica, and Peer Bork. 2024. ‘Interactive Tree of Life (iTOL) v6: Recent Updates to the Phylogenetic Tree Display and Annotation Tool’. Nucleic Acids Research 52 (W1): W78–82. 10.1093/nar/gkae268.

Li, Heng. 2023. ‘Protein-to-Genome Alignment with Miniprot’. Bioinformatics 39 (1): btad014. 10.1093/bioinformatics/btad014.

Lian, Xiaoping, Shilai Zhang, Guangfu Huang, Liyu Huang, Jing Zhang, and Fengyi Hu. 2021. ‘Confirmation of a Gametophytic Self-Incompatibility in *Oryza Longistaminata*’. Frontiers in Plant Science 12 (March): 1–9. 10.3389/fpls.2021.576340.

Lin, Zongcheng, Maurice Bosch, and Vernonica E. Franklin-Tong. 2026. ‘Several Plant Self-incompatibility Systems May Be Controlled by Atypical Receptor–Ligand Interactions’. The Plant Journal 126 (1): e70803. 10.1111/tpj.70803.

Lundquist, Arne. 1956. ‘Self-Incompatibility in Rye’. Hereditas 42 (3–4): 293–348. 10.1111/J.1601-5223.1956.TB03021.X.

Lundquist, Arne. 1962. ‘THE NATURE OF THE TWO-LOCI INCOMPATIBILITY SYSTEM IN GRASSES’. Hereditas 48 (1–2): 153–68. 10.1111/j.1601-5223.1962.tb01804.x.

Maenosono, Taiga, Kazuho Isono, Takanori Kuronuma, et al. 2024. ‘Exploring the Allelic Diversity of the Self-Incompatibility Gene Across Natural Populations in *Petunia* (Solanaceae)’. Genome Biology and Evolution 16 (12): evae270. 10.1093/gbe/evae270.

Manzanares, Chloé, Susanne Barth, Daniel Thorogood, et al. 2016. ‘A Gene Encoding a DUF247 Domain Protein Cosegregates with the S Self-Incompatibility Locus in Perennial Ryegrass’. Molecular Biology and Evolution 33 (4): 870–84. 10.1093/molbev/msv335.

Mascher, Martin, Todd A. Richmond, Daniel J. Gerhardt, et al. 2013. ‘Barley Whole Exome Capture: A Tool for Genomic Research in the Genus *Hordeum* and Beyond’. The Plant Journal 76 (3): 494–505. 10.1111/tpj.12294.

Murase, Kohji, Yoshitaka Moriwaki, Tomoyuki Mori, et al. 2020. ‘Mechanism of Self/Nonself-Discrimination in Brassica Self-Incompatibility’. Nature Communications 11 (1): 4916. 10.1038/s41467-020-18698-w.

Nagy, Istvan, Elisabeth Veeckman, Chang Liu, et al. 2022. ‘Chromosome-Scale Assembly and Annotation of the Perennial Ryegrass Genome’. BMC Genomics 23 (1): 505. 10.1186/s12864-022-08697-0.

Nei, Masatoshi, and Wen-Hsiung Li. 1979. ‘Mathematical Model for Studying Genetic Variation in Terms of Restriction Endonucleases.’ Proceedings of the National Academy of Sciences 76 (10): 5269–73. 10.1073/pnas.76.10.5269.

Nettancourt, Dreux de. 1977. *Incompatibility in Angiosperms*. (Berlin, Heidelberg), Monographs on Theoretical and Applied Genetics, vol. 3. 10.1007/978-3-662-12051-4.

Paape, Timothy, Takashi Miyake, Naoki Takebayashi, Diana Wolf, and Joshua R. Kohn. 2011. ‘Evolutionary Genetics of an S-Like Polymorphism in Papaveraceae with Putative Function in Self-Incompatibility’. PLoS ONE 6 (8): e23635. 10.1371/journal.pone.0023635.

Paradis, Emmanuel, and Klaus Schliep. 2019. ‘Ape 5.0: An Environment for Modern Phylogenetics and Evolutionary Analyses in R’. Bioinformatics 35 (3): 526–28. 10.1093/bioinformatics/bty633.

Paril, Jefferson, Gunjan Pandey, Emma M. Barnett, et al. 2022. ‘Rounding up the Annual Ryegrass Genome: High-Quality Reference Genome of Lolium Rigidum’. Frontiers in Genetics 13 (November): 1012694. 10.3389/fgene.2022.1012694.

R Core Team. 2025. R: A Language and Environment for Statistical Computing. V. 4.5.0. Vienna, Austria, Released. https://www.R-project.org/.

Revell, Liam J. 2024. ‘Phytools 2.0: An Updated R Ecosystem for Phylogenetic Comparative Methods (and Other Things).’ PeerJ (United States) 12: e16505. 10.7717/peerj.16505.

Rohner, Marius, Chloé Manzanares, Steven Yates, et al. 2023. ‘Fine-Mapping and Comparative Genomic Analysis Reveal the Gene Composition at the S and Z Self-Incompatibility Loci in Grasses’. Molecular Biology and Evolution 40 (1): msac259. 10.1093/molbev/msac259.

Sato, Keiichi, Takeshi Nishio, Ryo Kimura, et al. 2002. ‘Coevolution of the *S* -Locus Genes *SRK*, *SLG* and *SP11/SCR* in *Brassica Oleracea* and *B. Rapa*’. Genetics 162 (2): 931–40. 10.1093/genetics/162.2.931.

Savage, A. E., and J. S. Miller. 2006. ‘Gametophytic Self-Incompatibility in Lycium Parishii (Solanaceae): Allelic Diversity, Genealogical Structure, and Patterns of Molecular Evolution at the S-RNase Locus’. Heredity 96 (6): 434–44. 10.1038/sj.hdy.6800818.

Schrödinger, LLC. 2015. ‘The PyMOL Molecular Graphics System, Version 1.8’. Unpublished manuscript. November.

Shen, Wei, Botond Sipos, and Liuyang Zhao. 2024. ‘SeqKit2: A Swiss Army Knife for Sequence and Alignment Processing’. iMeta 3 (3): e191. 10.1002/imt2.191.

Sork, Victoria L., Shawn J. Cokus, Sorel T. Fitz-Gibbon, et al. 2022. ‘High-Quality Genome and Methylomes Illustrate Features Underlying Evolutionary Success of Oaks’. Nature Communications 13 (1): 2047. 10.1038/s41467-022-29584-y.

Stamatakis, Alexandros. 2014. ‘RAxML Version 8: A Tool for Phylogenetic Analysis and Post-Analysis of Large Phylogenies’. Bioinformatics 30 (9): 1312–13. 10.1093/bioinformatics/btu033.

Stein, Joshua C., Yeisoo Yu, Dario Copetti, et al. 2018. ‘Genomes of 13 Domesticated and Wild Rice Relatives Highlight Genetic Conservation, Turnover and Innovation across the Genus Oryza’. Nature Genetics 50 (2): 285–96. 10.1038/s41588-018-0040-0.

Steinbachs, Jennifer E., and Kent E. Holsinger. 2002. ‘S-RNase–Mediated Gametophytic Self-Incompatibility Is Ancestral in Eudicots’. Molecular Biology and Evolution 19 (6): 825–29. 10.1093/oxfordjournals.molbev.a004139.

Sun, Sijie, Jianli Wang, Yu Guan, et al. 2026. ‘High-Quality Chromosome-Level Genomes Reveal the Structure and Evolution of the *S* and *Z* Self-Incompatibility Loci in *Leymus Chinensis*’. *Plant Biotechnology Journal*, January 29, pbi.70565. 10.1111/pbi.70565.

Takayama, Seiji, and Akira Isogai. 2005. ‘Self-Incompatibility in Plants’. Annual Review of Plant Biology 56: 467–89. 10.1146/annurev.arplant.56.032604.144249.

Takuno, Shohei, Ryo Fujimoto, Tetsu Sugimura, et al. 2007. ‘Effects of Recombination on Hitchhiking Diversity in the Brassica Self-Incompatibility Locus Complex’. Genetics 177 (2): 949–58. 10.1534/genetics.107.073825.

Thomas Wong, Nhan Ly-Trong, Huaiyan Ren, et al. 2025. IQ-TREE 3: Phylogenomic Inference Software Using Complex Evolutionary Models. V. 3.0.1. Released May 5. 10.32942/X2P62N.

Thorogood, Daniel, Steven Yates, Chloé Manzanares, et al. 2017. ‘A Novel Multivariate Approach to Phenotyping and Association Mapping of Multi-Locus Gametophytic Self-Incompatibility Reveals S, Z, and Other Loci in a Perennial Ryegrass (Poaceae) Population’. Frontiers in Plant Science 8 (August). 10.3389/FPLS.2017.01331/ABSTRACT.

Veeckman, Elisabeth, Sabine Van Glabeke, Annelies Haegeman, et al. 2019. ‘Overcoming Challenges in Variant Calling: Exploring Sequence Diversity in Candidate Genes for Plant Development in Perennial Ryegrass (Lolium Perenne)’. DNA Research 26 (1): 1–12. 10.1093/dnares/dsy033.

Velmurugan, Janaki, Ewan Mollison, Susanne Barth, et al. 2016. ‘An Ultra-High Density Genetic Linkage Map of Perennial Ryegrass (Lolium Perenne) Using Genotyping by Sequencing (GBS) Based on a Reference Shotgun Genome Assembly’. Annals of Botany 118 (1): 71–87. 10.1093/aob/mcw081.

Vogt, Maximilian, Steven Yates, Timothy Sykes, Wilbert Luesink, Michael Koch, and Bruno Studer. 2020. ‘Developing Heterotic Groups for Successful Hybrid Breeding in Perennial Ryegrass’. Agronomy 10 (9): 1410. 10.3390/agronomy10091410.

Wannitikul, Pitchaporn, Pakorn Wattana-Amorn, Sukhita Sathitnaitham, et al. 2023. Disruption of a DUF247 Containing Protein Alters Cell Wall Polysaccharides and Reduces Growth in Arabidopsis. 10.3390/plants12101977.

Wheeler, Michael J., Barend H. J. De Graaf, Natalie Hadjiosif, et al. 2009. Identification of the Pollen Self-Incompatibility Determinant in Papaver Rhoeas. 10.1038/nature08027.

Wright, Sewall. 1939. ‘THE DISTRIBUTION OF SELF-STERILITY ALLELES IN POPULATIONS’. Genetics 24 (4): 538–52. 10.1093/genetics/24.4.538.

Yang, Bicheng, Danny Thorogood, Ian Armstead, and Susanne Barth. 2008. ‘How Far Are We from Unravelling Self-Incompatibility in Grasses?’ New Phytologist 178 (4): 740–53. 10.1111/j.1469-8137.2008.02421.x.

Yang, Ziheng. 2007. ‘PAML 4: Phylogenetic Analysis by Maximum Likelihood’. Molecular Biology and Evolution 24 (8): 1586–91. 10.1093/molbev/msm088.

Yang, Ziheng, Wendy S. W. Wong, and Rasmus Nielsen. 2005. ‘Bayes Empirical Bayes Inference of Amino Acid Sites Under Positive Selection’. Molecular Biology and Evolution 22 (4): 1107–18. 10.1093/molbev/msi097.

Zhang, Diyang, Yuan-Yuan Li, Xuewei Zhao, et al. 2024. ‘Molecular Insights into Self-Incompatibility Systems: From Evolution to Breeding’. Plant Communications 5 (2): 100719. 10.1016/j.xplc.2023.100719.

Zhang, Feifei, Yixi Liu, Fang Liu, et al. 2024. ‘Genome-Wide Characterization and Analysis of Rice DUF247 Gene Family’. BMC Genomics 25 (1): 613. 10.1186/s12864-024-10515-8.

Zhu, Tingting, Le Wang, Hélène Rimbert, et al. 2021. ‘Optical Maps Refine the Bread Wheat Triticum Aestivum Cv. Chinese Spring Genome Assembly.’ *The* Plant Journal : For Cell and Molecular Biology (England) 107 (1): 303–14. 10.1111/tpj.15289.

