## Supplementary information for "Evolutionary signatures of self-incompatibility genes in grasses: high diversity, trans-generic polymorphism and co-evolution"

### Supplementary materials

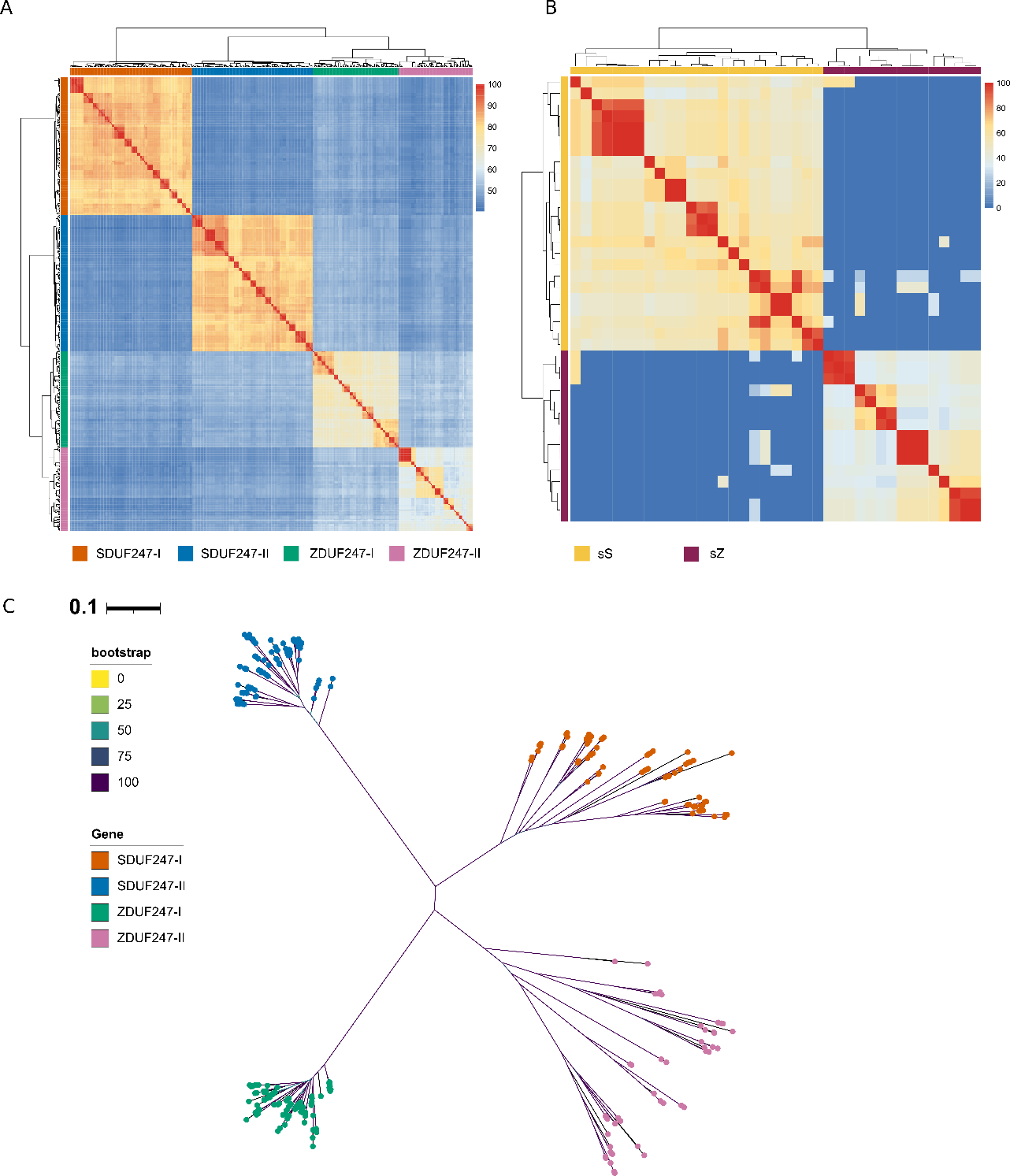

Figure S1. A pair-wise heat map of A) the four SI-DUF247 proteins and B) the sS and sZ. The protein similarity is indicated by a colour gradient from dark blue (0% similarity) to red (100% similarity). C) Maximum likelihood phylogenies of all four SI-DUF247.

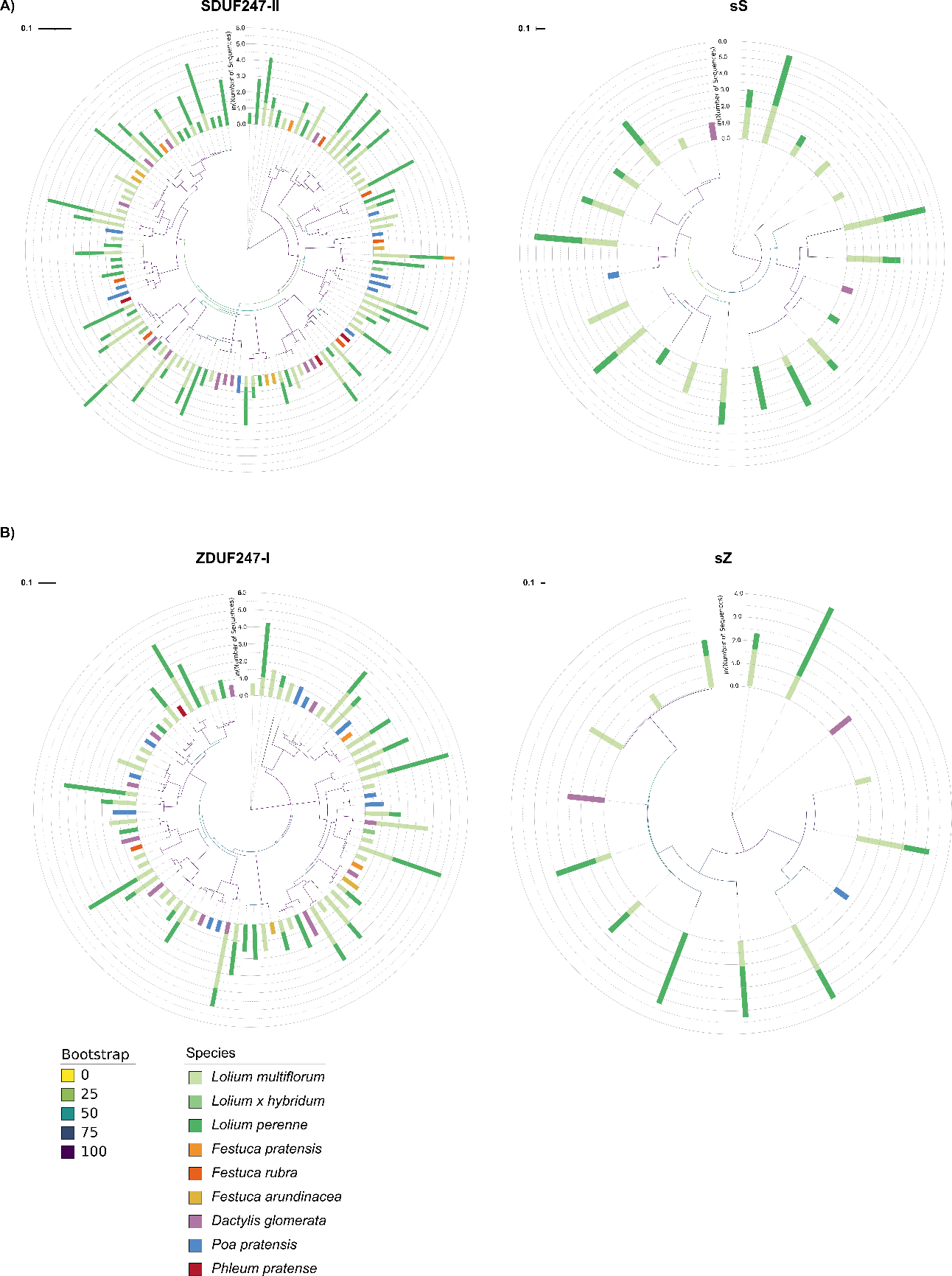

Figure S2. Maximum likelihood phylogenies of SI determinants A) SDUF247-II and sS and B) ZDUF247-I and sZ. Branch lengths represent the number of amino acid substitutions per site, and branch colors indicate bootstrap support. Bar colors refer to the species in which the protein variant was found. The bar height is proportional to the natural logarithm of the number of sequences represented in this data set.

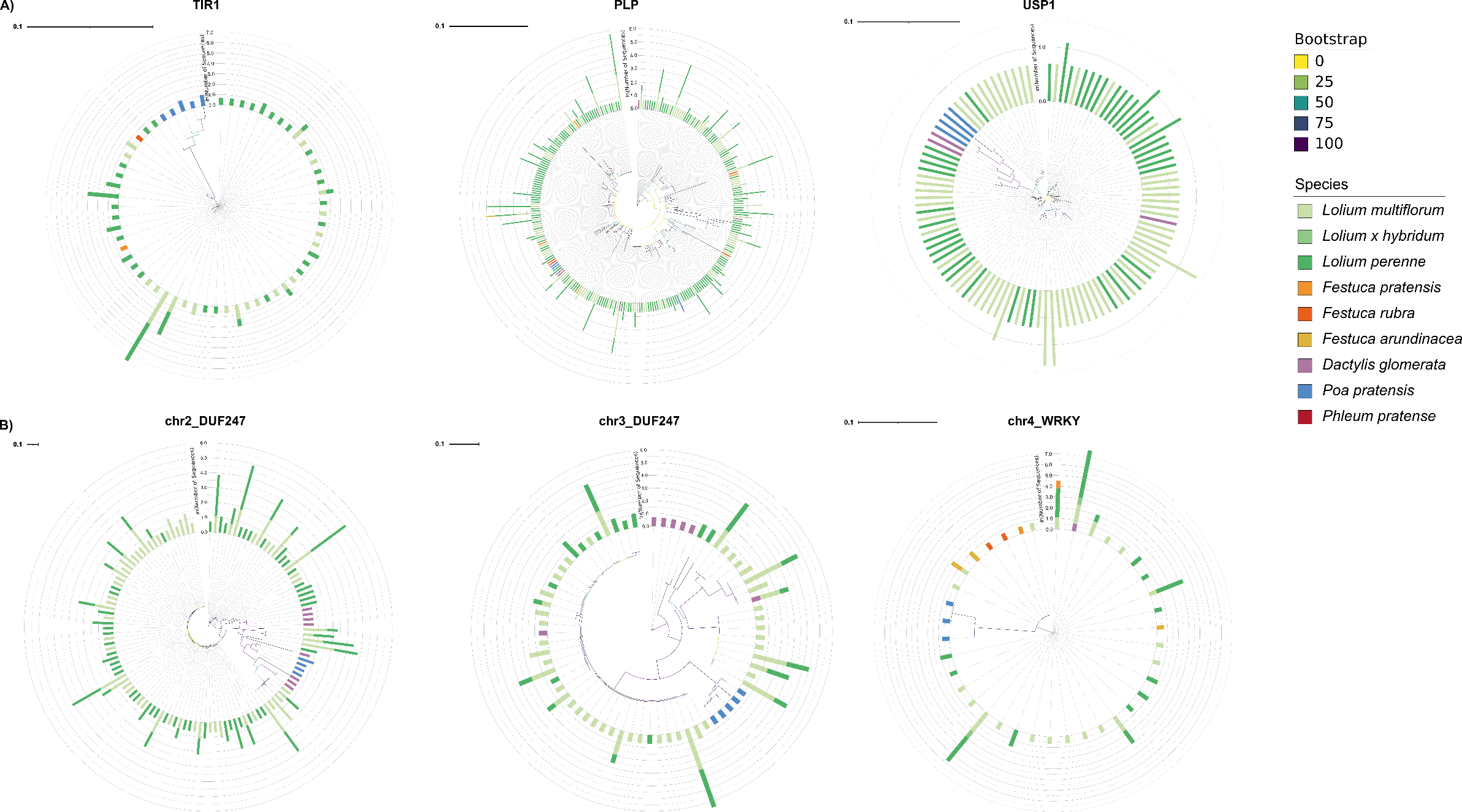

Figure S3. Maximum likelihood phylogenies of A) the flanking genes at the *S*-locus (*TIR1* and *PLP*) and the *Z*-locus (*USP1*); and B) the outgroup genes (two *DUF247* on chromosome 2 and 3 and *WRKY* on chromosome 4). Branch lengths represent the number of amino acid substitutions per site, and branch colors indicate bootstrap support. Bar colors refer to the species in which the protein variant was found. The bar height is proportional to the natural logarithm of the number of sequences represented in this data set.

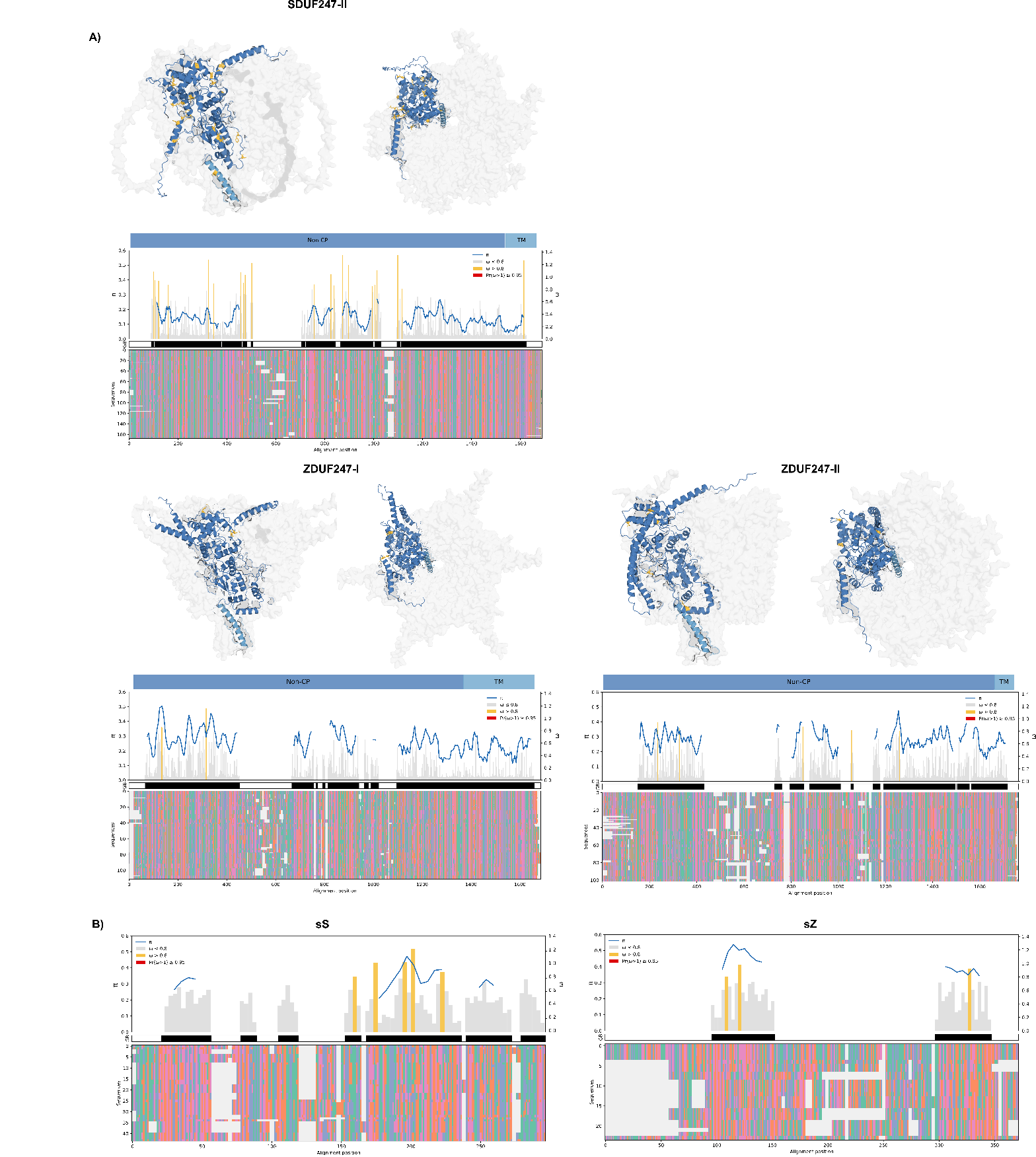

Figure S4. Codon substitution analysis results of A) *SI-DUF247* and B) female SI genes *sS* and *sZ*. From bottom to top: Nucleotide sequence alignment retained GBlocks, nucleotide diversity (π), the ratio of non-synonymous to synonymous substitution site (ω) across codons, and AlphaFold prediction of a homo-hexamer for SI-DUF247. The results of the condon substitution analysis was mapped on the predicted hexamer for the SI-DUF247; amino-acid are colored in yellow for neutral selection (mean ω > 0.8 but Pr(ω > 1) < 0.95) and red for positive selection (Pr(ω > 1) ≥ 0.95). Non-cytoplasmic (Non-CP) and transmembrane (TM) are indicated in dark and light blue.

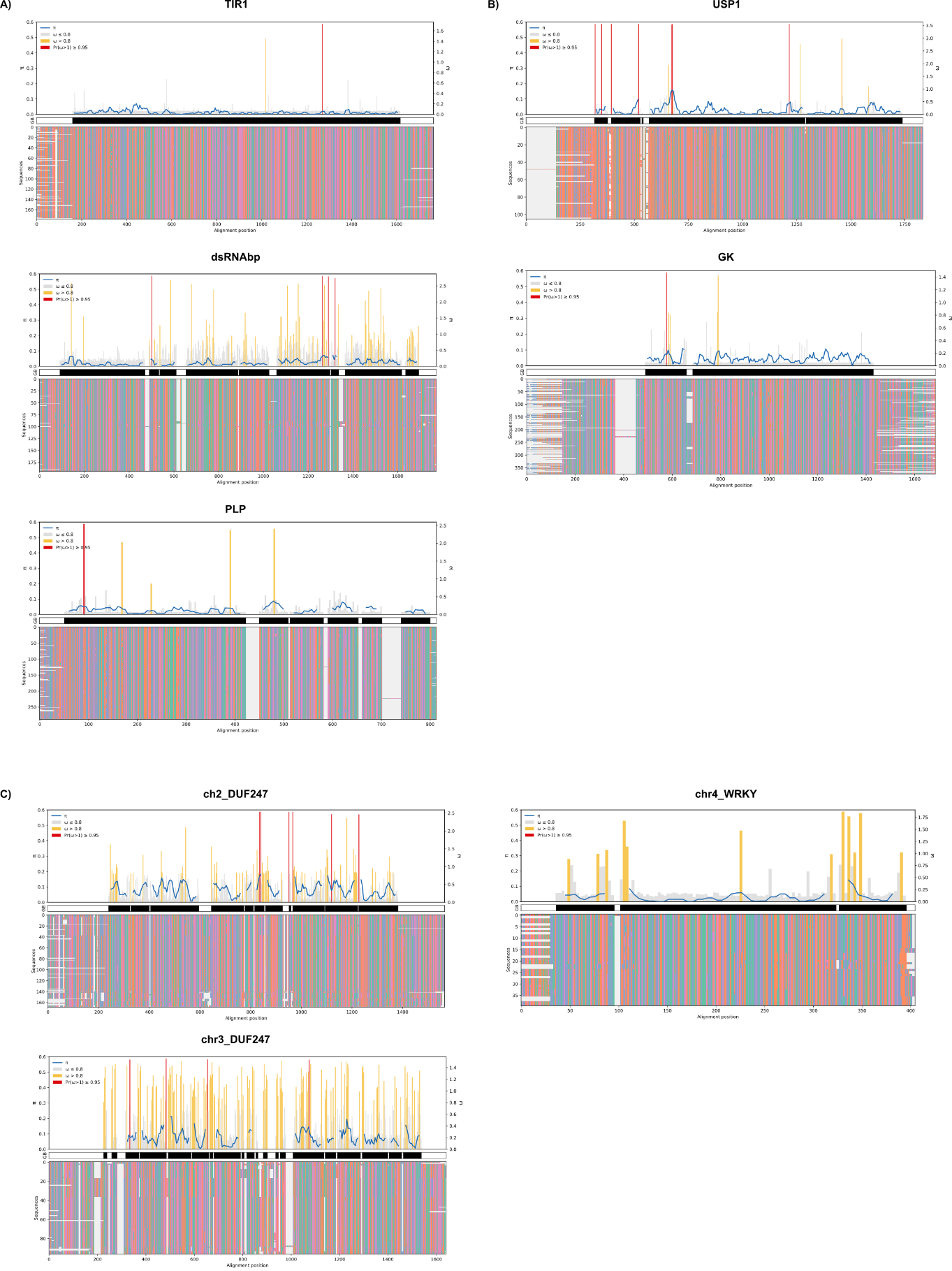

Figure S5. Nucleotide sequence alignment retained GBlocks, nucleotide diversity (π) and the ratio of non-synonymous to synonymous substitution site (ω) across codons for A) the *S*-locus flanking genes, B) the *Z*-locus flanking genes and C) the outgroup genes.

Table S1. Quality assessment of the *de novo* assembled *SI-DUF247*, *sS,* and *sZ* contigs. The region representing the gene sequence (from the start codon to the stop codon) is compared to the available reference sequences. The reference sequences consist of the reference-grade genome assembly of *L. perenne* genotype Kyuss (Frei et al. 2021), a diploid genome assembly of *Lolium perenne* genotype S23 Z (Rohner et al. 2023), the reference-grade genome assembly *L. perenne* genotype P226/135/16 and the diploid genome assembly of *Lolium multiflorum* Lam. Rabiosa (Copetti et al. 2021). If both alleles are accessible within the reference data, they are indicated by A and B after the gene name.

|  | **Genotype** | **Gene** | **Mistakes** | **Gene sequence length** | **Coverage** |
| --- | --- | --- | --- | --- | --- |
| ***Lolium perenne*** | Kyuss | *SDUF247-I* | 0 | 1593 | 175 |
|  |  | *SDUF247-II* | 0 | 1653 | 183 |
|  |  | *sS* | 0 | 321 | 81 |
|  |  | *ZDUF247-I* | 0 | 1590 | 161 |
|  |  | *ZDUF247-II* | 0 | 1662 | 147 |
|  |  | *sZ* | 0 | 333 | 88 |
|  | S23 Z | *SDUF247-I_A* | 0 | 1521 | 128 |
|  |  | *SDUF247-I_B* | 0 | 1596 | 150 |
|  |  | *SDUF247-II_A* | 0 | 1656 | 131 |
|  |  | *SDUF247-II_B* | 0 | 1596 | 128 |
|  |  | *sS_A* | 0 | 355 | 78 |
|  |  | *sS_B* | 0 | 364 | 77 |
|  |  | *ZDUF247-I_A* | 0 | 1638 | 143 |
|  |  | *ZDUF247-I_B* | 0 | 1617 | 155 |
|  |  | *ZDUF247-II_A* | 0 | 1647 | 135 |
|  |  | *ZDUF247-II_B* | 0 | 1644 | 127 |
|  |  | *sZ_A* | 0 | 363 | 38 |
|  |  | *sZ_B* | 0 | 448 | 49 |
|  | P226/135/16 | *SDUF247-I* | 0 | 1596 | 126 |
|  |  | *SDUF247-II* | 0 | 1662 | 150 |
|  |  | *sS* | 0 | 359 | 65 |
|  |  | *ZDUF247-I* | 0 | 1629 | 144 |
|  |  | *ZDUF247-II* | 0 | 1626 | 161 |
|  |  | *sZ* | 0 | 440 | 59 |
| ***Lolium multiflorum*** | Rabiosa | *SDUF247-I_A* | 0 | 1599 | 50 |
|  |  | *SDUF247-I_B* | 0 | 1521 | 1521 |
|  |  | *SDUF247-II_A* | 0 | 1656 | 52 |
|  |  | *SDUF247-II_B* | 0 | 1677 | 63 |
|  |  | *sS_A* | 0 | 364 | 22 |
|  |  | *sS_B* | 0 | 355 | 20 |
|  |  | *ZDUF247-I_A* | 0 | 1623 | 52 |
|  |  | *ZDUF247-I_B* | 0 | 1629 | 50 |
|  |  | *ZDUF247-II_A* | 0 | 1632 | 53 |
|  |  | *ZDUF247-II_B* | 0 | 1638 | 53 |
|  |  | *sZ_A* | 0 | 366 | 14 |
|  |  | *sZ_B* | 0 | 355 | 19 |

Table S2. Summary table of the codon substitution analysis performed with codeML. Estimates of the overall non-synonymous to synonymous substitution rate ratio (ω) under model M0 are shown. Likelihood ratio tests (LRT) were conducted to compare model M1a versus M2a and M7 versus M8. LRT statistics (2ΔlnL), degrees of freedom (df), and χ² test significance are shown.

|  |  | M0 | M1a vs. M2a | | M7 vs. M8 | | |
| --- | --- | --- | --- | --- | --- | --- | --- |
|  |  | ω | LRT | df | LRT | df | % of sites with ω > 1 |
| *S-locus* | *TIR1* | 0.043 | 2.12*** | 2 | 22.19*** | 2 | 0.46 (ω = 1.97) |
|  | *dsRNAbp* | 0.449 | 32.70*** | 2 | 39.88*** | 2 | 3.23 (ω = 3.31) |
|  | *SDUF247-I* | 0.294 | 4.52 | 2 | 10.02** | 2 | 2.07 (ω = 1.47) |
|  | *sS* | 0.446 | 0 | 2 | 0.97 | 2 | 9.62 (ω = 1.14) |
|  | *SDUF247-II* | 0.244 | 0 | 2 | 3.31 | 2 | 5.50 (ω = 1) |
|  | *PLP* | 0.165 | 29.75*** | 2 | 62.384*** | 2 | 1.99 (ω = 2.14) |
| *Z-locus* | *USP1* | 0.127 | 121.14*** | 2 | 127.25*** | 2 | 1.62 (ω = 3.95) |
|  | *ZDUF247-I* | 0.191 | 0 | 2 | 0 | 2 | 0.00 (ω = 2.89) |
|  | *sZ* | 0.433 | 0 | 2 | 0 | 2 | 0.00 (ω = 1) |
|  | *ZDUF247-II* | 0.192 | 0 | 2 | 0 | 2 | 0.00 (ω = 1) |
|  | *GK* | 0.064 | 0 | 2 | 1.297 | 2 | 0.00 (ω = 7.19) |
| Outgroup genes | *chr2-DUF247* | 0.409 | 30.21*** | 2 | 35.55*** | 2 | 2.82 (ω = 2.780) |
|  | *chr3-DUF247* | 0.568 | 8.37* | 2 | 12.55* | 2 | 14.57 (ω = 1.799) |
|  | *chr4-WRKY* | 0.056 | 0 | 2 | 0 | 2 | 3.98 (ω = 1.87) |
| ** : P ≤ 0.05, ** : P ≤ 0.01, *** : P ≤ 0.001* | | | | | | | |

|  |  | π | | | | | | | | | |
| --- | --- | --- | --- | --- | --- | --- | --- | --- | --- | --- | --- |
|  |  | all species | *Dactylis glomerata* | *Festuca pratensis* | *Festuca rubra* | *Festuca arundinacea* | *Lolium multiflorum* | *Lolium hybridium* | *Phleum pratense* | *Lolium perenne* | *Poa pratensis* |
| *S-locus* | *TIR1* | 0.013 | - | - | - | - | 0.009 | - | - | 0.009 | 0.020 |
|  | *dsRNAbp* | 0.023 | 0.127 | - | 0.028 | - | 0.015 | - | - | 0.015 | 0.034 |
|  | *SDUF247-I* | 0.134 | 0.132 | 0.147 | 0.141 | 0.095 | 0.135 | 0.144 | 0.134 | 0.124 | 0.116 |
|  | *sS* | 0.291 | 0.379 | - | - | - | 0.295 | - | - | 0.286 | - |
|  | *SDUF247-II* | 0.149 | 0.142 | 0.176 | 0.155 | 0.122 | 0.167 | 0.148 | 0.149 | 0.146 | 0.136 |
|  | *PLP* | 0.029 | 0.042 | 0.026 | 0.042 | 0.041 | 0.030 | 0.041 | - | 0.027 | 0.017 |
| *Z-locus* | *USP1* | 0.026 | 0.013 | - | - | - | 0.021 | - | - | 0.022 | 0.043 |
|  | *ZDUF247-I* | 0.265 | 0.266 | 0.314 | - | 0.283 | - | 0.268 | 0.269 | - | 0.258 |
|  | *sZ* | 0.403 | 0.417 | - | - | - | 0.417 | - | - | - | 0.421 |
|  | *ZDUF247-II* | 0.281 | 0.273 | - | - | 0.279 | - | 0.284 | 0.263 | - | 0.298 |
|  | *GK* | 0.046 | 0.053 | 0.037 | 0.036 | 0.044 | 0.043 | 0.043 | - | - | 0.034 |
| Outgroup genes | *chr2-DUF247* | 0.088 | 0.172 | - | - | - | 0.082 | - | - | 0.068 | 0.067 |
|  | *chr3-DUF247* | 0.088 | 0.093 | - | - | - | 0.078 | - | - | 0.085 | 0.028 |
|  | *chr4-WRKY* | 0.037 | - | 0.034 | - | 0.059 | 0.024 | 0.040 | - | 0.022 | 0.006 |

Table S3. Interspecific and intraspecific nucleotide diversity (π) per gene.

Table S4. Summary of the sequence acquisition for the probe design. The total number of sequences extracted for each target gene, the gene annotation name from *Lolium perenne* Kyuss (Frei et at. 2021) (positions within a chromosome (chr) are given when no annotation is present), and the source of the acquired sequences are provided.

| **Gene name** | **Number  of sequences** | **Gene annotation name from *Lolium perenne* Kyuss** | **Sources** |
| --- | --- | --- | --- |
| ***SDUF247-I*** | **145** | *chr1 38452229..38452229* | Genome assembly of paternal grandparent of the F2 biomass *L. perenne* population (Velmurugan et al. 2016) Allele reconstruction within a *L. perenne* S-locus mapping population (Manzanares et al. 2016) Genome assembly of *L. perenne* genotype 4540-9 (Honig et al. 2017) Genome assembly of *D. glomerata* genotype 2006-1 (Huang et al. 2020) Genome assembly of *L. multiflorum* (Knorst et al. 2019) Sequence capture assay using a germplasm collection of 736 *L. perenne* genotypes (Veeckman et al. 2019) Genome assembly of *F. pratensis* genotype 1700/16 (Samy et al. 2020) Genome assembly of *F. pratensis* genotype HF7_2 (Samy et al. 2020) Genome assembly of *L. perenne* genotype Kyuss (Frei et al. 2021) Genome assembly of *L. multiflorum* genotype Rabiosa (Copetti et al. 2021) Genome assembly of *L. perenne* genotype P226/135/16 (Nagy et al. 2022) Sequence capture assay using mainly *L. perenne* breeding material from Switzerland (Rohner 2022) Genome assembly of *L. rigidium* genotype Wagga Wagga (Paril et al. 2022) Genome assembly of *L. perenne* genotype S23 Z (Rohner et al. 2023) Genome assembly of *F. pratensis* genotype Kolombus (in-house, unpublished) Genome assembly of *L. multiflorum* Adret2 (in-house, unpublished) Genome assembly of *L. multiflorum* M2289 (in-house, unpublished) Genome assembly of *L. multiflorum* genotype Sikem (Chen et al. 2025) Genome assembly of *Festulolium* (NRGENE, unpublished) |
| ***SDUF247-II*** | **111** | *KYUS_G_chr1.6214* | Genome assembly of paternal grandparent of the F2 biomass *L. perenne* population (Velmurugan et al. 2016) Genome assembly of *L. perenne* genotype 4540-9 (Honig et al. 2017) Genome assembly of *D. glomerata* genotype 2006-1 (Huang et al. 2020) Genome assembly of *L. multiflorum* (Knorst et al. 2019) Sequence capture assay using a germplasm collection of 736 *L. perenne* genotypes (Veeckman et al. 2019) Genome assembly of *F. pratensis* genotype 1700/16 (Samy et al. 2020) Genome assembly of *F. pratensis* genotype HF7_2 (Samy et al. 2020) Genome assembly of *L. perenne* genotype Kyuss (Frei et al. 2021) Genome assembly of *L. multiflorum* genotype Rabiosa (Copetti et al. 2021) Genome assembly of *L. perenne* genotype P226/135/16 (Nagy et al. 2022) Sequence capture assay using mainly *L. perenne* breeding material from Switzerland (Rohner et al. 2023) Genome assembly of *L. rigidium* genotype Wagga Wagga (Paril et al. 2022) Genome assembly of *L. perenne* genotype S23 Z (Rohner et al. 2023) Genome assembly of *F. pratensis* genotype Kolombus (in-house, unpublished) Genome assembly of *L. multiflorum* Adret2 (in-house, unpublished) Genome assembly of *L. multiflorum* M2289 (in-house, unpublished) Genome assembly of *L. multiflorum* genotype Sikem (Chen et al. 2025) Genome assembly of *Festulolium* (NRGENE, unpublished) |
| ***sS*** | **23** | *chr1 38449393..38449713* | Genome assembly of paternal grandparent of the F2 biomass *L. perenne* population (Velmurugan et al. 2016) Genome assembly of *L. perenne* genotype 4540-9 (Honig et al. 2017) Genome assembly of *D. glomerata* genotype 2006-1 (Huang et al. 2020) Genome assembly of *L. multiflorum* (Knorst et al. 2019) Genome assembly of *F. pratensis* genotype 1700/16 (Samy et al. 2020) Genome assembly of *F. pratensis* genotype HF7_2 (Samy et al. 2020) Genome assembly of *L. perenne* genotype Kyuss (Frei et al. 2021) Genome assembly of *L. multiflorum* genotype Rabiosa (Copetti et al. 2021) Genome assembly of *L. perenne* genotype P226/135/16 (Nagy et al. 2022) Genome assembly of *L. rigidium* genotype Wagga Wagga (Paril et al. 2022) Genome assembly of *L. perenne* genotype S23 Z (Rohner et al. 2023) Genome assembly of *F. pratensis* genotype Kolombus (in-house, unpublished) Genome assembly of *L. multiflorum* Adret2 (in-house, unpublished) Genome assembly of *L. multiflorum* M2289 (in-house, unpublished) Genome assembly of L. multiflorum genotype Sikem (Chen et al. 2025)) Genome assembly of *Festulolium* (NRGENE, unpublished) |
| ***ZDUF247-I*** | **71** | *KYUS_G_chr2.53348* | Genome assembly of paternal grandparent of the F2 biomass *L. perenne* population (Velmurugan et al. 2016) Genome assembly of *L. perenne* genotype 4540-9 (Honig et al. 2017) Genome assembly of *D. glomerata* genotype 2006-1 (Huang et al. 2020) Genome assembly of *L. multiflorum* (Knorst et al. 2019) Sequence capture assay using a germplasm collection of 736 *L. perenne* genotypes (Veeckman et al. 2019) Genome assembly of *F. pratensis* genotype 1700/16 (Samy et al. 2020) Genome assembly of *F. pratensis* genotype HF7_2 (Samy et al. 2020) Genome assembly of *L. perenne* genotype Kyuss (Frei et al. 2021) Genome assembly of *L. multiflorum* genotype Rabiosa (Copetti et al. 2021) Genome assembly of *L. perenne* genotype P226/135/16 (Nagy et al. 2022) Sequence capture assay using mainly *L. perenne* breeding material from Switzerland (Rohner 2022) Genome assembly of *L. rigidium* genotype Wagga Wagga (Paril et al. 2022) Genome assembly of *L. perenne* genotype S23 Z (Rohner et al. 2023) *L. perenne* BAC clone P205C9H17P (Rohner et al. 2023) Genome assembly of *F. pratensis* genotype Kolombus (in-house, unpublished) Genome assembly of *L. multiflorum* M2289 (in-house, unpublished) Genome assembly of L. multiflorum genotype Sikem (Chen et al. 2025) Genome assembly of *Festulolium* (NRGENE, unpublished) |
| ***ZDUF247-II*** | **69** | *chr2 33280078..332799119* | Genome assembly of paternal grandparent of the F2 biomass *L. perenne* population (Velmurugan et al. 2016) Genome assembly of *L. perenne* genotype 4540-9 (Honig et al. 2017) Genome assembly of *D. glomerata* genotype 2006-1 (Huang et al. 2020) Genome assembly of *L. multiflorum* (Knorst et al. 2019) Sequence capture assay using a germplasm collection of 736 *L. perenne* genotypes (Veeckman et al. 2019) Genome assembly of *F. pratensis* genotype 1700/16 (Samy et al. 2020) Genome assembly of *F. pratensis* genotype HF7_2 (Samy et al. 2020) Genome assembly of *L. perenne* genotype Kyuss (Frei et al. 2021) Genome assembly of *L. multiflorum* genotype Rabiosa (Copetti et al. 2021) Genome assembly of *L. perenne* genotype P226/135/16 (Nagy et al. 2022) Sequence capture assay using mainly *L. perenne* breeding material from Switzerland (Rohner 2022) Genome assembly of *L. rigidium* genotype Wagga Wagga (Paril et al. 2022) Genome assembly of *L. perenne* genotype S23 Z (Rohner et al. 2023) *L. perenne* BAC clone P205C9H17P (Rohner et al. 2023) Genome assembly of *F. pratensis* genotype Kolombus (in-house, unpublished) Genome assembly of *L. multiflorum* M2289 (in-house, unpublished) Genome assembly of *L. multiflorum* genotype Sikem (Chen et al. 2025) Genome assembly of *Festulolium* (NRGENE, unpublished) |
| ***sZ*** | **14** | *KYUS_G_chr2.53336* | Genome assembly of *L. perenne* genotype 4540-9 (Honig et al. 2017) Genome assembly of *D. glomerata* genotype 2006-1 (Huang et al. 2020) Genome assembly of *L. multiflorum* (Knorst et al. 2019) Genome assembly of *F. pratensis* genotype 1700/16 (Samy et al. 2020) Genome assembly of *L. perenne* genotype Kyuss (Frei et al. 2021) Genome assembly of *L. multiflorum* genotype Rabiosa (Copetti et al. 2021) Genome assembly of *L. perenne* genotype P226/135/16 (Nagy et al. 2022) Genome assembly of *L. rigidium* genotype Wagga Wagga (Paril et al. 2022) Genome assembly of *L. perenne* genotype S23 Z (Rohner et al. 2023) *L. perenne* BAC clone P205C9H17P (Rohner et al. 2023) Genome assembly of *F. pratensis* genotype Kolombus (in-house, unpublished) Genome assembly of *L. multiflorum* M2289 (in-house, unpublished) Genome assembly of *Festulolium* (NRGENE, unpublished) |
| ***TIR1*** | **13** | *KYUS_G_chr1.6316* | Genome assembly of *L. perenne* genotype Kyuss (Frei et al. 2021) Genome assembly of *L. multiflorum* genotype Rabiosa (Copetti et al. 2021) Genome assembly of *L. perenne* genotype P226/135/16 (Nagy et al. 2022) Genome assembly of *L. rigidium* genotype Wagga Wagga (Paril et al. 2022) Genome assembly of *L. perenne* genotype S23 Z (Rohner et al. 2023) Genome assembly of *F. pratensis* genotype Kolombus (in-house, unpublished) Genome assembly of *Festulolium* (NRGENE, unpublished) |
| ***dsRNAbp*** | **13** | *KYUS_G_chr1.6309* | Genome assembly of *L. perenne* genotype Kyuss (Frei et al. 2021) Genome assembly of *L. multiflorum* genotype Rabiosa (Copetti et al. 2021) Genome assembly of *L. perenne* genotype P226/135/16 (Nagy et al. 2022) Genome assembly of *L. rigidium* genotype Wagga Wagga (Paril et al. 2022) Genome assembly of *L. perenne* genotype S23 Z (Rohner et al. 2023) Genome assembly of *F. pratensis* genotype Kolombus (in-house, unpublished) Genome assembly of *Festulolium* (NRGENE, unpublished) |
| ***PLP*** | **19** | *KYUS_G_chr1.6210* | Genome assembly of *L. perenne* genotype Kyuss (Frei et al. 2021) Genome assembly of *L. multiflorum* genotype Rabiosa (Copetti et al. 2021) Genome assembly of *L. perenne* genotype P226/135/16 (Nagy et al. 2022) Genome assembly of *L. rigidium* genotype Wagga Wagga (Paril et al. 2022) Genome assembly of *L. perenne* genotype S23 Z (Rohner et al. 2023) Genome assembly of *F. pratensis* genotype Kolombus (in-house, unpublished) Genome assembly of *Festulolium* (NRGENE, unpublished) |
| ***USP1*** | **13** | *KYUS_G_chr2.53349* | Genome assembly of *L. perenne* genotype Kyuss (Frei et al. 2021) Genome assembly of *L. multiflorum* genotype Rabiosa (Copetti et al. 2021) Genome assembly of *L. perenne* genotype P226/135/16 (Nagy et al. 2022) Genome assembly of *L. rigidium* genotype Wagga Wagga (Paril et al. 2022) Genome assembly of *L. perenne* genotype S23 Z (Rohner et al. 2023) *L. perenne* BAC clone P205C9H17P (Rohner et al. 2023) Genome assembly of *F. pratensis* genotype Kolombus (in-house, unpublished) Genome assembly of *L. multiflorum* genotype Sikem (Chen et al. 2025) Genome assembly of *Festulolium* (NRGENE, unpublished)) |
| ***GK*** | **15** | *KYUS_G_chr2.53334* | Genome assembly of *L. perenne* genotype Kyuss (Frei et al. 2021) Genome assembly of *L. multiflorum* genotype Rabiosa (Copetti et al. 2021) Genome assembly of *L. perenne* genotype P226/135/16 (Nagy et al. 2022) Genome assembly of *L. rigidium* genotype Wagga Wagga (Paril et al. 2022) Genome assembly of *L. perenne* genotype S23 Z (Rohner et al. 2023) *L. perenne* BAC clone P205C9H17P (Rohner et al. 2023) Genome assembly of *F. pratensis* genotype Kolombus (in-house, unpublished) Genome assembly of *L. multiflorum* genotype Sikem (Chen et al. 2025) Genome assembly of Festulolium (NRGENE, unpublished) |
| ***chr2.48100*** | **12** | *chr2.48100* | Genome assembly of *L. perenne* genotype Kyuss (Frei et al. 2021) Genome assembly of *L. multiflorum* genotype Rabiosa (Copetti et al. 2021) Genome assembly of *L. perenne* genotype P226/135/16 (Nagy et al. 2022) Genome assembly of *L. perenne* genotype S23 Z (Rohner et al. 2023)  Genome assembly of *F. pratensis* genotype Kolombus (in-house, unpublished) Genome assembly of *L. multiflorum* genotype Sikem (Chen et al. 2025) Genome assembly of Festulolium (NRGENE, unpublished) |
| ***chr3.27160*** | **13** | *chr3.27160* | Genome assembly of *L. perenne* genotype Kyuss (Frei et al. 2021) Genome assembly of *L. multiflorum* genotype Rabiosa (Copetti et al. 2021) Genome assembly of *L. perenne* genotype P226/135/16 (Nagy et al. 2022) Genome assembly of *L. perenne* genotype S23 Z (Rohner et al. 2023)  Genome assembly of *F. pratensis* genotype Kolombus (in-house, unpublished) Genome assembly of *L. multiflorum* genotype Sikem (Chen et al. 2025) Genome assembly of Festulolium (NRGENE, unpublished) |
| ***chr4.WRKY*** | **5** | *chr4. 100592301..100593873* | Genome assembly of *L. perenne* genotype Kyuss (Frei et al. 2021) Genome assembly of *L. multiflorum* M2289 (in-house, unpublished)  Genome assembly of Festulolium (NRGENE, unpublished) |

Table S5. Detailed overview of the genotypes sequenced in the two rounds of targeted sequence capture (TSC).

| **Genotype name** | | **Genotype full name** | **Species** | | **Description and source** | | **Sequence capture** |
| --- | --- | --- | --- | --- | --- | --- | --- |
| 5-B674G | | 5-B674G | *Lolium perenne* | | F13 population (Thorogood et al. 2017) | | TSC1 |
| 12-B674G | | 12-B674G | *Lolium perenne* | | F13 population (Thorogood et al. 2017) | | TSC1 |
| 23-B674G | | 23-B674G | *Lolium perenne* | | F13 population (Thorogood et al. 2017) | | TSC1 |
| 27-B674G | | 27-B674G | *Lolium perenne* | | F13 population (Thorogood et al. 2017) | | TSC1 |
| 33-B674G | | 33-B674G | *Lolium perenne* | | F13 population (Thorogood et al. 2017) | | TSC1 |
| 39-B674G | | 39-B674G | *Lolium perenne* | | F13 population (Thorogood et al. 2017) | | TSC1 |
| 42-B674G | | 42-B674G | *Lolium perenne* | | F13 population (Thorogood et al. 2017) | | TSC1 |
| 43-B674G | | 43-B674G | *Lolium perenne* | | F13 population (Thorogood et al. 2017) | | TSC1 |
| 45-B674G | | 45-B674G | *Lolium perenne* | | F13 population (Thorogood et al. 2017) | | TSC1 |
| 46-B674G | | 46-B674G | *Lolium perenne* | | F13 population (Thorogood et al. 2017) | | TSC1 |
| 47-B674G | | 47-B674G | *Lolium perenne* | | F13 population (Thorogood et al. 2017) | | TSC1 |
| 115-B674G | | 115-B674G | *Lolium perenne* | | F13 population (Thorogood et al. 2017) | | TSC1 |
| 123-B674G | | 123-B674G | *Lolium perenne* | | F13 population (Thorogood et al. 2017) | | TSC1 |
| 131-B674G | | 131-B674G | *Lolium perenne* | | F13 population (Thorogood et al. 2017) | | TSC1 |
| 132-B674G | | 132-B674G | *Lolium perenne* | | F13 population (Thorogood et al. 2017) | | TSC1 |
| 133-B674G | | 133-B674G | *Lolium perenne* | | F13 population (Thorogood et al. 2017) | | TSC1 |
| 134-B674G | | 134-B674G | *Lolium perenne* | | F13 population (Thorogood et al. 2017) | | TSC1 |
| 136-B674G | | 136-B674G | *Lolium perenne* | | F13 population (Thorogood et al. 2017) | | TSC1 |
| 138-B674G | | 138-B674G | *Lolium perenne* | | F13 population (Thorogood et al. 2017) | | TSC1 |
| 143-B674G | | 143-B674G | *Lolium perenne* | | F13 population (Thorogood et al. 2017) | | TSC1 |
| 145-B674G | | 145-B674G | *Lolium perenne* | | F13 population (Thorogood et al. 2017) | | TSC1 |
| 146-B674G | | 146-B674G | *Lolium perenne* | | F13 population (Thorogood et al. 2017) | | TSC1 |
| 147-B674G | | 147-B674G | *Lolium perenne* | | F13 population (Thorogood et al. 2017) | | TSC1 |
| 148-B674G | | 148-B674G | *Lolium perenne* | | F13 population (Thorogood et al. 2017) | | TSC1 |
| 204-B674G | | 204-B674G | *Lolium perenne* | | F13 population (Thorogood et al. 2017) | | TSC1 |
| 209-B674G | | 209-B674G | *Lolium perenne* | | F13 population (Thorogood et al. 2017) | | TSC1 |
| 213-B674G | | 213-B674G | *Lolium perenne* | | F13 population (Thorogood et al. 2017) | | TSC1 |
| 216-B674G | | 216-B674G | *Lolium perenne* | | F13 population (Thorogood et al. 2017) | | TSC1 |
| 218-B674G | | 218-B674G | *Lolium perenne* | | F13 population (Thorogood et al. 2017) | | TSC1 |
| 222-B674G | | 222-B674G | *Lolium perenne* | | F13 population (Thorogood et al. 2017) | | TSC1 |
| 223-B674G | | 223-B674G | *Lolium perenne* | | F13 population (Thorogood et al. 2017) | | TSC1 |
| 228-B674G | | 228-B674G | *Lolium perenne* | | F13 population (Thorogood et al. 2017) | | TSC1 |
| 231-B674G | | 231-B674G | *Lolium perenne* | | F13 population (Thorogood et al. 2017) | | TSC1 |
| 233-B674G | | 233-B674G | *Lolium perenne* | | F13 population (Thorogood et al. 2017) | | TSC1 |
| 235-B674G | | 235-B674G | *Lolium perenne* | | F13 population (Thorogood et al. 2017) | | TSC1 |
| 238-B674G | | 238-B674G | *Lolium perenne* | | F13 population (Thorogood et al. 2017) | | TSC1 |
| 294-B674G | | 294-B674G | *Lolium perenne* | | F13 population (Thorogood et al. 2017) | | TSC1 |
| 295-B674G | | 295-B674G | *Lolium perenne* | | F13 population (Thorogood et al. 2017) | | TSC1 |
| 298-B674G | | 298-B674G | *Lolium perenne* | | F13 population (Thorogood et al. 2017) | | TSC1 |
| 301-B674G | | 301-B674G | *Lolium perenne* | | F13 population (Thorogood et al. 2017) | | TSC1 |
| 307-B674G | | 307-B674G | *Lolium perenne* | | F13 population (Thorogood et al. 2017) | | TSC1 |
| 314-B674G | | 314-B674G | *Lolium perenne* | | F13 population (Thorogood et al. 2017) | | TSC1 |
| 318-B674G | | 318-B674G | *Lolium perenne* | | F13 population (Thorogood et al. 2017) | | TSC1 |
| 320-B674G | | 320-B674G | *Lolium perenne* | | F13 population (Thorogood et al. 2017) | | TSC1 |
| 323-B674G | | 323-B674G | *Lolium perenne* | | F13 population (Thorogood et al. 2017) | | TSC1 |
| 329-B674G | | 329-B674G | *Lolium perenne* | | F13 population (Thorogood et al. 2017) | | TSC1 |
| 331-B674G | | 331-B674G | *Lolium perenne* | | F13 population (Thorogood et al. 2017) | | TSC1 |
| 332-B674G | | 332-B674G | *Lolium perenne* | | F13 population (Thorogood et al. 2017) | | TSC1 |
| 334-B674G | | 334-B674G | *Lolium perenne* | | F13 population (Thorogood et al. 2017) | | TSC1 |
| 337-B674G | | 337-B674G | *Lolium perenne* | | F13 population (Thorogood et al. 2017) | | TSC1 |
| 342-B674G | | 342-B674G | *Lolium perenne* | | F13 population (Thorogood et al. 2017) | | TSC1 |
| 345-B674G | | 345-B674G | *Lolium perenne* | | F13 population (Thorogood et al. 2017) | | TSC1 |
| 347-B674G | | 347-B674G | *Lolium perenne* | | F13 population (Thorogood et al. 2017) | | TSC1 |
| 348-B674G | | 348-B674G | *Lolium perenne* | | F13 population (Thorogood et al. 2017) | | TSC1 |
| 2001a | | 2001a_NAM | *Lolium multiflorum* | | Parent of the NAM population (Kiesbauer et al. 2026) | | TSC1 |
| 2001b | | 2001b_NAM | *Lolium multiflorum* | | Parent of the NAM population (Kiesbauer et al. 2026) | | TSC1 |
| 2003 | | 2003_NAM | *Lolium multiflorum* | | Parent of the NAM population (Kiesbauer et al. 2026) | | TSC1 |
| 2004 | | 2004_NAM | *Lolium multiflorum* | | Parent of the NAM population (Kiesbauer et al. 2026) | | TSC1 |
| 2009 | | 2009_NAM | *Lolium multiflorum* | | Parent of the NAM population (Kiesbauer et al. 2026) | | TSC1 |
| 2010 | | 2010_NAM | *Lolium multiflorum* | | Parent of the NAM population (Kiesbauer et al. 2026) | | TSC1 |
| 2012 | | 2012_NAM | *Lolium multiflorum* | | Parent of the NAM population (Kiesbauer et al. 2026) | | TSC1 |
| 2014 | | 2014_NAM | *Lolium multiflorum* | | Parent of the NAM population (Kiesbauer et al. 2026) | | TSC1 |
| 2017 | | 2017_NAM | *Lolium multiflorum* | | Parent of the NAM population (Kiesbauer et al. 2026) | | TSC1 |
| 2020 | | 2020_NAM | *Lolium multiflorum* | | Parent of the NAM population (Kiesbauer et al. 2026) | | TSC1 |
| 2021 | | 2021_NAM | *Lolium multiflorum* | | Parent of the NAM population (Kiesbauer et al. 2026) | | TSC1 |
| 2026 | | 2026_NAM | *Lolium multiflorum* | | Parent of the NAM population (Kiesbauer et al. 2026) | | TSC1 |
| 2028 | | 2028_NAM | *Lolium multiflorum* | | Parent of the NAM population (Kiesbauer et al. 2026) | | TSC1 |
| 2030 | | 2030_NAM | *Lolium multiflorum* | | Parent of the NAM population (Kiesbauer et al. 2026) | | TSC1 |
| 2031 | | 2031_NAM | *Lolium multiflorum* | | Parent of the NAM population (Kiesbauer et al. 2026) | | TSC1 |
| 2032 | | 2032_NAM | *Lolium multiflorum* | | Parent of the NAM population (Kiesbauer et al. 2026) | | TSC1 |
| 2033 | | 2033_NAM | *Lolium multiflorum* | | Parent of the NAM population (Kiesbauer et al. 2026) | | TSC1 |
| 7588 | | 7588 | *Dactylis glomerata  var. barexcel* | | In-house grass reference material (ETH Zurich, Switzerland) | | TSC1 |
| 7602 | | 7602 | *Festuca pratensis  var. pradel* | | In-house grass reference material (ETH Zurich, Switzerland) | | TSC1 |
| 7603 | | 7603 | *Festuca pratensis  var. pradel* | | In-house grass reference material (ETH Zurich, Switzerland) | | TSC1 |
| 7617 | | 7617 | *Festuca arundinacea  var. dulcia* | | In-house grass reference material (ETH Zurich, Switzerland) | | TSC1 |
| 7618 | | 7618 | *Festuca arundinacea  var. dulcia* | | In-house grass reference material (ETH Zurich, Switzerland) | | TSC1 |
| 7632 | | 7632 | *Festuca rubra  var. echo* | | In-house grass reference material (ETH Zurich, Switzerland) | | TSC1 |
| 7633 | | 7633 | *Festuca rubra  var. echo* | | In-house grass reference material (ETH Zurich, Switzerland) | | TSC1 |
| 7648 | | 7648 | *Lolium multiflorum  var. westerwoldicum* | | In-house grass reference material (ETH Zurich, Switzerland) | | TSC1 |
| 7677 | | 7677 | *Lolium x hybridum  var. ocadia* | | In-house grass reference material (ETH Zurich, Switzerland) | | TSC1 |
| 7698 | | 7698 | *Lolium multiflorum  var. italicum* | | In-house grass reference material (ETH Zurich, Switzerland) | | TSC1 |
| 7820 | | 7820 | *Poa pratensis  var. tommy* | | In-house grass reference material (ETH Zurich, Switzerland) | | TSC1 |
| 7821 | | 7821 | *Poa pratensis  var. tommy* | | In-house grass reference material (ETH Zurich, Switzerland) | | TSC1 |
| 7822 | | 7822 | *Phleum pratense  var. tiller* | | In-house grass reference material (ETH Zurich, Switzerland) | | TSC1 |
| ARARA-A | | ARARA-A | *Lolium perenne* | | Variety (Agroscope Reckenholz, Switzerland) | | TSC1 |
| ARARA-B | | ARARA-B | *Lolium perenne* | | Variety (Agroscope Reckenholz, Switzerland) | | TSC1 |
| Arolus | | Arolus | *Lolium perenne* | | Variety (Agroscope Reckenholz, Switzerland) | | TSC1 |
| DH2015-9906-47 | | DH2015-9906-47 | *Lolium perenne* | | Variety (Agroscope Reckenholz, Switzerland) | | TSC1 |
| F1-39 | | F1-39 | *Lolium perenne* | | Genotype used in SI research (ETH Zurich, Switzerland) | | TSC1 |
| F2-2-200 | | F2-2-200 | *Lolium perenne* | | F2 biomass populaton | | TSC1 |
| F2-2-282 | | F2-2-282 | *Lolium perenne* | | F2 biomass populaton | | TSC1 |
| F2-2-363 | | F2-2-363 | *Lolium perenne* | | F2 biomass populaton | | TSC1 |
| F2-210 | | F2-210 | *Lolium perenne* | | F2 biomass populaton | | TSC1 |
| F2-257 | | F2-257 | *Lolium perenne* | | F2 biomass populaton | | TSC1 |
| F2-277 | | F2-277 | *Lolium perenne* | | F2 biomass populaton | | TSC1 |
| IOWA506-12 | | IOWA506-12 | *Lolium perenne* | | IOWA population | | TSC1 |
| IOWA-506-13 | | IOWA-506-13 | *Lolium perenne* | | IOWA population | | TSC1 |
| IOWA-P3C2 | | IOWA-P3C2 | *Lolium perenne* | | IOWA population | | TSC1 |
| IOWA-P5E10 | | IOWA-P5E10 | *Lolium perenne* | | IOWA population | | TSC1 |
| IOWA-P11C9 | | IOWA-P11C9 | *Lolium perenne* | | IOWA population | | TSC1 |
| IOWA-P13G5 | | IOWA-P13G5 | *Lolium perenne* | | IOWA population | | TSC1 |
| IOWA-P15A10 | | IOWA-P15A10 | *Lolium perenne* | | IOWA population | | TSC1 |
| IOWA-P16H1 | | IOWA-P16H1 | *Lolium perenne* | | IOWA population | | TSC1 |
| IOWA-P20H12 | | IOWA-P20H12 | *Lolium perenne* | | IOWA population | | TSC1 |
| M34-MIC | | M34-MIC | *Lolium perenne* | | Breeding program at DSV (Zaden, Netherlands) | | TSC1 |
| M35-MIC | | M35-MIC | *Lolium perenne* | | Breeding program at DSV (Zaden, Netherlands) | | TSC1 |
| M611-02-27 | | M611-02-27 | *Lolium perenne* | | M611 population | | TSC1 |
| M-06001 | | M-06001 | *Lolium perenne* | | Diploid nursery population (Agroscope Reckenholz, Switzerland) | | TSC1 |
| M-06002 | | M-06002 | *Lolium perenne* | | Diploid nursery population (Agroscope Reckenholz, Switzerland) | | TSC1 |
| M-06003 | | M-06003 | *Lolium perenne* | | Diploid nursery population (Agroscope Reckenholz, Switzerland) | | TSC1 |
| M-06004 | | M-06004 | *Lolium perenne* | | Diploid nursery population (Agroscope Reckenholz, Switzerland) | | TSC1 |
| M-06005 | | M-06005 | *Lolium perenne* | | Diploid nursery population (Agroscope Reckenholz, Switzerland) | | TSC1 |
| M-06006 | | M-06006 | *Lolium perenne* | | Diploid nursery population (Agroscope Reckenholz, Switzerland) | | TSC1 |
| M-06007 | | M-06007 | *Lolium perenne* | | Diploid nursery population (Agroscope Reckenholz, Switzerland) | | TSC1 |
| M-06008 | | M-06008 | *Lolium perenne* | | Diploid nursery population (Agroscope Reckenholz, Switzerland) | | TSC1 |
| M-06009 | | M-06009 | *Lolium perenne* | | Diploid nursery population (Agroscope Reckenholz, Switzerland) | | TSC1 |
| M-06010 | | M-06010 | *Lolium perenne* | | Diploid nursery population (Agroscope Reckenholz, Switzerland) | | TSC1 |
| M-06011 | | M-06011 | *Lolium perenne* | | Diploid nursery population (Agroscope Reckenholz, Switzerland) | | TSC1 |
| M-06012 | | M-06012 | *Lolium perenne* | | Diploid nursery population (Agroscope Reckenholz, Switzerland) | | TSC1 |
| M-06013 | | M-06013 | *Lolium perenne* | | Diploid nursery population (Agroscope Reckenholz, Switzerland) | | TSC1 |
| M-06014 | | M-06014 | *Lolium perenne* | | Diploid nursery population (Agroscope Reckenholz, Switzerland) | | TSC1 |
| M-06015 | | M-06015 | *Lolium perenne* | | Diploid nursery population (Agroscope Reckenholz, Switzerland) | | TSC1 |
| M-06017 | | M-06017 | *Lolium perenne* | | Diploid nursery population (Agroscope Reckenholz, Switzerland) | | TSC1 |
| M-06018 | | M-06018 | *Lolium perenne* | | Diploid nursery population (Agroscope Reckenholz, Switzerland) | | TSC1 |
| M-06019 | | M-06019 | *Lolium perenne* | | Diploid nursery population (Agroscope Reckenholz, Switzerland) | | TSC1 |
| M-06020 | | M-06020 | *Lolium perenne* | | Diploid nursery population (Agroscope Reckenholz, Switzerland) | | TSC1 |
| M-06021 | | M-06021 | *Lolium perenne* | | Diploid nursery population (Agroscope Reckenholz, Switzerland) | | TSC1 |
| M-06022 | | M-06022 | *Lolium perenne* | | Diploid nursery population (Agroscope Reckenholz, Switzerland) | | TSC1 |
| M-06023 | | M-06023 | *Lolium perenne* | | Diploid nursery population (Agroscope Reckenholz, Switzerland) | | TSC1 |
| M-06025 | | M-06025 | *Lolium perenne* | | Diploid nursery population (Agroscope Reckenholz, Switzerland) | | TSC1 |
| M-06026 | | M-06026 | *Lolium perenne* | | Diploid nursery population (Agroscope Reckenholz, Switzerland) | | TSC1 |
| M-06027 | | M-06027 | *Lolium perenne* | | Diploid nursery population (Agroscope Reckenholz, Switzerland) | | TSC1 |
| M-06028 | | M-06028 | *Lolium perenne* | | Diploid nursery population (Agroscope Reckenholz, Switzerland) | | TSC1 |
| M-06029 | | M-06029 | *Lolium perenne* | | Diploid nursery population (Agroscope Reckenholz, Switzerland) | | TSC1 |
| M-06030 | | M-06030 | *Lolium perenne* | | Diploid nursery population (Agroscope Reckenholz, Switzerland) | | TSC1 |
| M-06031 | | M-06031 | *Lolium perenne* | | Diploid nursery population (Agroscope Reckenholz, Switzerland) | | TSC1 |
| M-06032 | | M-06032 | *Lolium perenne* | | Diploid nursery population (Agroscope Reckenholz, Switzerland) | | TSC1 |
| M-06033 | | M-06033 | *Lolium perenne* | | Diploid nursery population (Agroscope Reckenholz, Switzerland) | | TSC1 |
| M-06034 | | M-06034 | *Lolium perenne* | | Diploid nursery population (Agroscope Reckenholz, Switzerland) | | TSC1 |
| M-06035 | | M-06035 | *Lolium perenne* | | Diploid nursery population (Agroscope Reckenholz, Switzerland) | | TSC1 |
| M-06036 | | M-06036 | *Lolium perenne* | | Diploid nursery population (Agroscope Reckenholz, Switzerland) | | TSC1 |
| M-06037 | | M-06037 | *Lolium perenne* | | Diploid nursery population (Agroscope Reckenholz, Switzerland) | | TSC1 |
| M-06038 | | M-06038 | *Lolium perenne* | | Diploid nursery population (Agroscope Reckenholz, Switzerland) | | TSC1 |
| M-06039 | | M-06039 | *Lolium perenne* | | Diploid nursery population (Agroscope Reckenholz, Switzerland) | | TSC1 |
| M-06040 | | M-06040 | *Lolium perenne* | | Diploid nursery population (Agroscope Reckenholz, Switzerland) | | TSC1 |
| M-06041 | | M-06041 | *Lolium perenne* | | Diploid nursery population (Agroscope Reckenholz, Switzerland) | | TSC1 |
| M-06042 | | M-06042 | *Lolium perenne* | | Diploid nursery population (Agroscope Reckenholz, Switzerland) | | TSC1 |
| M-06043 | | M-06043 | *Lolium perenne* | | Diploid nursery population (Agroscope Reckenholz, Switzerland) | | TSC1 |
| M-06044 | | M-06044 | *Lolium perenne* | | Diploid nursery population (Agroscope Reckenholz, Switzerland) | | TSC1 |
| M-06045 | | M-06045 | *Lolium perenne* | | Diploid nursery population (Agroscope Reckenholz, Switzerland) | | TSC1 |
| M6111-02-36 | | M6111-02-36 | *Lolium perenne* | | Diploid nursery population (Agroscope Reckenholz, Switzerland) | | TSC1 |
| M6115-02-17 | | M6115-02-17 | *Lolium perenne* | | Diploid nursery population (Agroscope Reckenholz, Switzerland) | | TSC1 |
| M6115-02-18 | | M6115-02-18 | *Lolium perenne* | | Diploid nursery population (Agroscope Reckenholz, Switzerland) | | TSC1 |
| M-07133 | | M-07133 | *Lolium perenne* | | Tetraploid nursery population (Agroscope Reckenholz, Switzerland) | | TSC1 |
| M-07134 | | M-07134 | *Lolium perenne* | | Tetraploid nursery population (Agroscope Reckenholz, Switzerland) | | TSC1 |
| M-07139 | | M-07139 | *Lolium perenne* | | Tetraploid nursery population (Agroscope Reckenholz, Switzerland) | | TSC1 |
| M-07144 | | M-07144 | *Lolium perenne* | | Tetraploid nursery population (Agroscope Reckenholz, Switzerland) | | TSC1 |
| M-07147 | | M-07147 | *Lolium perenne* | | Tetraploid nursery population (Agroscope Reckenholz, Switzerland) | | TSC1 |
| M-07148 | | M-07148 | *Lolium perenne* | | Tetraploid nursery population (Agroscope Reckenholz, Switzerland) | | TSC1 |
| M-07149 | | M-07149 | *Lolium perenne* | | Tetraploid nursery population (Agroscope Reckenholz, Switzerland) | | TSC1 |
| M-07150 | | M-07150 | *Lolium perenne* | | Tetraploid nursery population (Agroscope Reckenholz, Switzerland) | | TSC1 |
| M-07151 | | M-07151 | *Lolium perenne* | | Tetraploid nursery population (Agroscope Reckenholz, Switzerland) | | TSC1 |
| M-07153 | | M-07153 | *Lolium perenne* | | Tetraploid nursery population (Agroscope Reckenholz, Switzerland) | | TSC1 |
| M-07155 | | M-07155 | *Lolium perenne* | | Tetraploid nursery population (Agroscope Reckenholz, Switzerland) | | TSC1 |
| M-07157 | | M-07157 | *Lolium perenne* | | Tetraploid nursery population (Agroscope Reckenholz, Switzerland) | | TSC1 |
| M-07160 | | M-07160 | *Lolium perenne* | | Tetraploid nursery population (Agroscope Reckenholz, Switzerland) | | TSC1 |
| NgB | | NgB | *Lolium perenne* | | Genotype used in SI research (ETH Zurich, Switzerland) | | TSC1 |
| NV20-F1-30 | | NV20-F1-30 | *Lolium perenne* | | Genotype used in SI research (ETH Zurich, Switzerland) | | TSC1 |
| S23-Z-WT | | S23 Z | *Lolium perenne* | | Sequenced genotype (Rohner et al. 2023) | | TSC1 |
| VRNA-1038 | | VRNA-1038 | *Lolium perenne* | | Genotype used in SI research (ETH Zurich, Switzerland) | | TSC1 |
| XANTHIA-24 | | XANTHIA-24 | *Lolium perenne* | | Genotype used in SI research (ETH Zurich, Switzerland) | | TSC1 |
| Pp1 | | Pp1 | *Poa pratensis* | | Common meadow-grass breeding material (Agroscope Reckenholz, Switzerland) | | TSC2 |
| Pp2 | | Pp2 | *Poa pratensis* | | Common meadow-grass breeding material (Agroscope Reckenholz, Switzerland) | | TSC2 |
| Pp3 | | Pp3 | *Poa pratensis* | | Common meadow-grass breeding material (Agroscope Reckenholz, Switzerland) | | TSC2 |
| Pp4 | | Pp4 | *Poa pratensis* | | Common meadow-grass breeding material (Agroscope Reckenholz, Switzerland) | | TSC2 |
| Pp5 | | Pp5 | *Poa pratensis* | | Common meadow-grass breeding material (Agroscope Reckenholz, Switzerland) | | TSC2 |
| M2002 | | M2002_NAM | *Lolium multiflorum* | | Considered as a parent of the NAM population (Kiesbauer et al. 2026) | | TSC2 |
| M2007 | | M2007_NAM | *Lolium multiflorum* | | Considered as a parent of the NAM population (Kiesbauer et al. 2026) | | TSC2 |
| M2013 | | M2013_NAM | *Lolium multiflorum* | | Considered as a parent of the NAM population (Kiesbauer et al. 2026) | | TSC2 |
| M2016 | | M2016_NAM | *Lolium multiflorum* | | Considered as a parent of the NAM population (Kiesbauer et al. 2026) | | TSC2 |
| M2018 | | M2018_NAM | *Lolium multiflorum* | | Considered as a parent of the NAM population (Kiesbauer et al. 2026) | | TSC2 |
| M2019 | | M2019_NAM | *Lolium multiflorum* | | Considered as a parent of the NAM population (Kiesbauer et al. 2026) | | TSC2 |
| M2022 | | M2022_NAM | *Lolium multiflorum* | | Considered as a parent of the NAM population (Kiesbauer et al. 2026) | | TSC2 |
| M2024 | | M2024_NAM | *Lolium multiflorum* | | Considered as a parent of the NAM population (Kiesbauer et al. 2026) | | TSC2 |
| M2025 | | M2025_NAM | *Lolium multiflorum* | | Considered as a parent of the NAM population (Kiesbauer et al. 2026) | | TSC2 |
| M2027 | | M2027_NAM | *Lolium multiflorum* | | Considered as a parent of the NAM population (Kiesbauer et al. 2026) | | TSC2 |
| IOWA1 | | IOWA Plate 3 C2 | *Lolium perenne* | | F2 SC population (Cropano et al. 2021) | | TSC2 |
| IOWA2 | | IOWA Plate 20 M12 | *Lolium perenne* | | F2 SC population (Cropano et al. 2021) | | TSC2 |
| IOWA3 | | IOWA Plate 11 C9 | *Lolium perenne* | | F2 SC population (Cropano et al. 2021) | | TSC2 |
| IOWA4 | | IOWA Plate 5 E10 | *Lolium perenne* | | F2 SC population (Cropano et al. 2021) | | TSC2 |
| IOWA5 | | IOWA Plate 5 G11 | *Lolium perenne* | | F2 SC population (Cropano et al. 2021) | | TSC2 |
| IOWA6 | | IOWA Plate 17 H9 | *Lolium perenne* | | F2 SC population (Cropano et al. 2021) | | TSC2 |
| Restorer (CMS) | | Restorer (CMS) | *Lolium perenne* | | Genotype involved in cytoplasmic male sterility (Vogt et al. 2020) | | TSC2 |
| Maintainer (CMS) | | Maintainer (CMS) | *Lolium perenne* | | Genotype involved in cytoplasmic male sterility (Vogt et al. 2020) | | TSC2 |
| Steril (CMS) | | Steril (CMS) | *Lolium perenne* | | Genotype involved in cytoplasmic male sterility (Vogt et al. 2020) | | TSC2 |
| Arara A | | Arara A | *Lolium perenne* | | Genotype of a commercial perennial ryegrasss variety (Agroscope Reckenholz, Switzerland) | | TSC2 |
| Arara B | | Arara B | *Lolium perenne* | | Genotype of a commercial perennial ryegrasss variety (Agroscope Reckenholz, Switzerland) | | TSC2 |
| NgB | | NgB | *Lolium perenne* | | Genotype used in SI research (Kopecký et al. 2009) | | TSC2 |
| VrnA | | VrnA | *Lolium perenne* | | Genotype used in SI research (Manzanares et al. 2016 & Rohner et al. 2023) | | TSC2 |
| F1-30 | | F1-30 | *Lolium perenne* | | Genotype used in SI research (Manzanares et al. 2016 & Rohner et al. 2023) | | TSC2 |
| F1-39 | | F1-39 | *Lolium perenne* | | Genotype used in SI research (Manzanares et al. 2016 & Rohner et al. 2023) | | TSC2 |
| Veyo | Veyo | | *Lolium perenne* | Genotype used in SI research (Manzanares et al. 2016 & Rohner et al. 2023) | | TSC2 | |
| P235/59/150 | P235/59/150 | | *Lolium perenne* | Genotype used in SI research (Manzanares et al. 2016) | | TSC2 | |
| Lm_13 | Lm_13_F1_BS_POP_2018 | | *Lolium multiflorum* | Genotype with bacterial wilt resistance (Goettelmann et al. 2023) | | TSC2 | |
| Lm_16 | Lm_16_F1_BS_POP_2018 | | *Lolium multiflorum* | Genotype with bacterial wilt resistance (Goettelmann 2023) | | TSC2 | |
| Lm_27 | Lm_27_F1_BS_POP_2018 | | *Lolium multiflorum* | Genotype with bacterial wilt resistance (Goettelmann 2023) | | TSC2 | |
| Lm_38 | Lm_38_F1_BS_POP_2018 | | *Lolium multiflorum* | Genotype with bacterial wilt resistance (Goettelmann 2023) | | TSC2 | |
| Lm_41 | Lm_41_F1_BS_POP_2018 | | *Lolium multiflorum* | Genotype with bacterial wilt resistance (Goettelmann 2023) | | TSC2 | |
| Lm_73 | Lm_73_F1_BS_POP_2018 | | *Lolium multiflorum* | Genotype with bacterial wilt resistance (Goettelmann 2023) | | TSC2 | |
| Lm_81 | Lm_81_F1_BS_POP_2018 | | *Lolium multiflorum* | Genotype with bacterial wilt resistance (Goettelmann 2023) | | TSC2 | |
| Lm_84 | Lm_84_F1_BS_POP_2018 | | *Lolium multiflorum* | Genotype with bacterial wilt resistance (Goettelmann 2023) | | TSC2 | |
| Lm_99 | Lm_99_F1_BS_POP_2018 | | *Lolium multiflorum* | Genotype with bacterial wilt resistance (Goettelmann 2023) | | TSC2 | |
| Lm_112 | Lm_112_F1_BS_POP_2018 | | *Lolium multiflorum* | Genotype with bacterial wilt resistance (Goettelmann 2023) | | TSC2 | |
| Lm_117 | Lm_117_F1_BS_POP_2018 | | *Lolium multiflorum* | Genotype with bacterial wilt resistance (Goettelmann 2023) | | TSC2 | |
| Lm_121 | Lm_121_F1_BS_POP_2018 | | *Lolium multiflorum* | Genotype with bacterial wilt resistance (Goettelmann 2023) | | TSC2 | |
| Lm_125 | Lm_125_F1_BS_POP_2018 | | *Lolium multiflorum* | Genotype with bacterial wilt resistance (Goettelmann 2023) | | TSC2 | |
| LmD1 | G2242_4729_01 | | *Lolium multiflorum* | Italian ryegrass breeding material (Agroscope Reckenholz, Switzerland) | | TSC2 | |
| LmD2 | G2242_4729_02 | | *Lolium multiflorum* | Italian ryegrass breeding material (Agroscope Reckenholz, Switzerland) | | TSC2 | |
| LmD3 | G2242_4730_01 | | *Lolium multiflorum* | Italian ryegrass breeding material (Agroscope Reckenholz, Switzerland) | | TSC2 | |
| LmD4 | G2242_4730_02 | | *Lolium multiflorum* | Italian ryegrass breeding material (Agroscope Reckenholz, Switzerland) | | TSC2 | |
| LmD5 | G2242_4731_01 | | *Lolium multiflorum* | Italian ryegrass breeding material (Agroscope Reckenholz, Switzerland) | | TSC2 | |
| LmD24 | G2242_4809_01 | | *Lolium multiflorum* | Italian ryegrass breeding material (Agroscope Reckenholz, Switzerland) | | TSC2 | |
| LmD7 | G2242_4732_01 | | *Lolium multiflorum* | Italian ryegrass breeding material (Agroscope Reckenholz, Switzerland) | | TSC2 | |
| LmD8 | G2242_4733_01 | | *Lolium multiflorum* | Italian ryegrass breeding material (Agroscope Reckenholz, Switzerland) | | TSC2 | |
| LmD27 | G2242_4811_01 | | *Lolium multiflorum* | Italian ryegrass breeding material (Agroscope Reckenholz, Switzerland) | | TSC2 | |
| LmD32 | G2242_4814_02 | | *Lolium multiflorum* | Italian ryegrass breeding material (Agroscope Reckenholz, Switzerland) | | TSC2 | |
| LmD11 | G2242_4735_02 | | *Lolium multiflorum* | Italian ryegrass breeding material (Agroscope Reckenholz, Switzerland) | | TSC2 | |
| LmD12 | G2242_4801_01 | | *Lolium multiflorum* | Italian ryegrass breeding material (Agroscope Reckenholz, Switzerland) | | TSC2 | |
| LmD13 | G2242_4801_02 | | *Lolium multiflorum* | Italian ryegrass breeding material (Agroscope Reckenholz, Switzerland) | | TSC2 | |
| LmD14 | G2242_4802_01 | | *Lolium multiflorum* | Italian ryegrass breeding material (Agroscope Reckenholz, Switzerland) | | TSC2 | |
| LmD15 | G2242_4803_02 | | *Lolium multiflorum* | Italian ryegrass breeding material (Agroscope Reckenholz, Switzerland) | | TSC2 | |
| LmD16 | G2242_4803_03 | | *Lolium multiflorum* | Italian ryegrass breeding material (Agroscope Reckenholz, Switzerland) | | TSC2 | |
| LmD17 | G2242_4804_01 | | *Lolium multiflorum* | Italian ryegrass breeding material (Agroscope Reckenholz, Switzerland) | | TSC2 | |
| LmD18 | G2242_4805_01 | | *Lolium multiflorum* | Italian ryegrass breeding material (Agroscope Reckenholz, Switzerland) | | TSC2 | |
| LmD33 | G2242_4815_01 | | *Lolium multiflorum* | Italian ryegrass breeding material (Agroscope Reckenholz, Switzerland) | | TSC2 | |
| LmD54 | G2242_4830_01 | | *Lolium multiflorum* | Italian ryegrass breeding material (Agroscope Reckenholz, Switzerland) | | TSC2 | |
| LmD56 | G2242_4831_01 | | *Lolium multiflorum* | Italian ryegrass breeding material (Agroscope Reckenholz, Switzerland) | | TSC2 | |
| LmD57 | G2242_4832_01 | | *Lolium multiflorum* | Italian ryegrass breeding material (Agroscope Reckenholz, Switzerland) | | TSC2 | |
| LmD59 | G2242_4834_01 | | *Lolium multiflorum* | Italian ryegrass breeding material (Agroscope Reckenholz, Switzerland) | | TSC2 | |
| LmD60 | G2242_4835_02 | | *Lolium multiflorum* | Italian ryegrass breeding material (Agroscope Reckenholz, Switzerland) | | TSC2 | |
| LmD20 | G2242_4806_02 | | *Lolium multiflorum* | Italian ryegrass breeding material (Agroscope Reckenholz, Switzerland) | | TSC2 | |
| LmD21 | G2242_4807_01 | | *Lolium multiflorum* | Italian ryegrass breeding material (Agroscope Reckenholz, Switzerland) | | TSC2 | |
| LmD22 | G2242_4808_01 | | *Lolium multiflorum* | Italian ryegrass breeding material (Agroscope Reckenholz, Switzerland) | | TSC2 | |
| LmD23 | G2242_4808_02 | | *Lolium multiflorum* | Italian ryegrass breeding material (Agroscope Reckenholz, Switzerland) | | TSC2 | |
| LmD25 | G2242_4810_02 | | *Lolium multiflorum* | Italian ryegrass breeding material (Agroscope Reckenholz, Switzerland) | | TSC2 | |
| LmD30 | G2242_4813_01 | | *Lolium multiflorum* | Italian ryegrass breeding material (Agroscope Reckenholz, Switzerland) | | TSC2 | |
| LmD34 | G2242_4816_01 | | *Lolium multiflorum* | Italian ryegrass breeding material (Agroscope Reckenholz, Switzerland) | | TSC2 | |
| LmD35 | G2242_4816_02 | | *Lolium multiflorum* | Italian ryegrass breeding material (Agroscope Reckenholz, Switzerland) | | TSC2 | |
| LmD36 | G2242_4817_01 | | *Lolium multiflorum* | Italian ryegrass breeding material (Agroscope Reckenholz, Switzerland) | | TSC2 | |
| LmD37 | G2242_4818_01 | | *Lolium multiflorum* | Italian ryegrass breeding material (Agroscope Reckenholz, Switzerland) | | TSC2 | |
| LmD42 | G2242_4821_01 | | *Lolium multiflorum* | Italian ryegrass breeding material (Agroscope Reckenholz, Switzerland) | | TSC2 | |
| LmD45 | G2242_4823_03 | | *Lolium multiflorum* | Italian ryegrass breeding material (Agroscope Reckenholz, Switzerland) | | TSC2 | |
| LmD46 | G2242_4824_01 | | *Lolium multiflorum* | Italian ryegrass breeding material (Agroscope Reckenholz, Switzerland) | | TSC2 | |
| LmD48 | G2242_4825_02 | | *Lolium multiflorum* | Italian ryegrass breeding material (Agroscope Reckenholz, Switzerland) | | TSC2 | |
| LmD49 | G2242_4826_01 | | *Lolium multiflorum* | Italian ryegrass breeding material (Agroscope Reckenholz, Switzerland) | | TSC2 | |
| LmD50 | G2242_4827_01 | | *Lolium multiflorum* | Italian ryegrass breeding material (Agroscope Reckenholz, Switzerland) | | TSC2 | |
| LmD51 | G2242_4827_02 | | *Lolium multiflorum* | Italian ryegrass breeding material (Agroscope Reckenholz, Switzerland) | | TSC2 | |
| M34_17 | M34_17 | | *Lolium perenne* | M34 and M35 population (Cropano 2021) | | TSC2 | |
| M34_18 | M34_18 | | *Lolium perenne* | M34 and M35 population (Cropano 2021) | | TSC2 | |
| M35_5 | M35_5 | | *Lolium perenne* | M34 and M35 population (Cropano 2021) | | TSC2 | |
| M34_10_CG | M34_10_CG | | *Lolium perenne* | M34 and M35 population (Cropano 2021) | | TSC2 | |
| M35_13 | M35_13 | | *Lolium perenne* | M34 and M35 population (Cropano 2021) | | TSC2 | |
| M35_2 | M35_2 | | *Lolium perenne* | M34 and M35 population (Cropano 2021) | | TSC2 | |
| Dg1 | Dg1 | | *Dactylis glomerata* | Orchard grass breeding material (Agroscope Reckenholz, Switzerland) | | TSC2 | |
| Dg2 | Dg2 | | *Dactylis glomerata* | Orchard grass breeding material (Agroscope Reckenholz, Switzerland) | | TSC2 | |
| Dg3 | Dg3 | | *Dactylis glomerata* | Orchard grass breeding material (Agroscope Reckenholz, Switzerland) | | TSC2 | |
| Dg4 | Dg4 | | *Dactylis glomerata* | Orchard grass breeding material (Agroscope Reckenholz, Switzerland) | | TSC2 | |
| Dg5 | Dg5 | | *Dactylis glomerata* | Orchard grass breeding material (Agroscope Reckenholz, Switzerland) | | TSC2 | |
| M2001 | M2001_NAM | | *Lolium multiflorum* | Parent of the NAM population (Kiesbauer et al. 2026) | | TSC2 | |
| M2003 | M2003_NAM | | *Lolium multiflorum* | Parent of the NAM population (Kiesbauer et al. 2026) | | TSC2 | |
| M2004 | M2004_NAM | | *Lolium multiflorum* | Parent of the NAM population (Kiesbauer et al. 2026) | | TSC2 | |
| M2006 | M2006_NAM | | *Lolium multiflorum* | Parent of the NAM population (Kiesbauer et al. 2026) | | TSC2 | |
| M2008 | M2008_NAM | | *Lolium multiflorum* | Parent of the NAM population (Kiesbauer et al. 2026) | | TSC2 | |
| M2009 | M2009_NAM | | *Lolium multiflorum* | Parent of the NAM population (Kiesbauer et al. 2026) | | TSC2 | |
| M2010 | M2010_NAM | | *Lolium multiflorum* | Parent of the NAM population (Kiesbauer et al. 2026) | | TSC2 | |
| M2011 | M2011_NAM | | *Lolium multiflorum* | Parent of the NAM population (Kiesbauer et al. 2026) | | TSC2 | |
| M2012 | M2012_NAM | | *Lolium multiflorum* | Parent of the NAM population (Kiesbauer et al. 2026) | | TSC2 | |
| M2014 | M2014_NAM | | *Lolium multiflorum* | Parent of the NAM population (Kiesbauer et al. 2026) | | TSC2 | |
| M2015 | M2015_NAM | | *Lolium multiflorum* | Parent of the NAM population (Kiesbauer et al. 2026) | | TSC2 | |
| M2017 | M2017_NAM | | *Lolium multiflorum* | Parent of the NAM population (Kiesbauer et al. 2026) | | TSC2 | |
| M2020 | M2020_NAM | | *Lolium multiflorum* | Parent of the NAM population (Kiesbauer et al. 2026) | | TSC2 | |
| M2021 | M2021_NAM | | *Lolium multiflorum* | Parent of the NAM population (Kiesbauer et al. 2026) | | TSC2 | |
| M2023 | M2023_NAM | | *Lolium multiflorum* | Parent of the NAM population (Kiesbauer et al. 2026) | | TSC2 | |
| M2026 | M2026_NAM | | *Lolium multiflorum* | Parent of the NAM population (Kiesbauer et al. 2026) | | TSC2 | |
| M2028 | M2028_NAM | | *Lolium multiflorum* | Parent of the NAM population (Kiesbauer et al. 2026) | | TSC2 | |
| M2029 | M2029_NAM | | *Lolium multiflorum* | Parent of the NAM population (Kiesbauer et al. 2026) | | TSC2 | |
| M2030 | M2030_NAM | | *Lolium multiflorum* | Parent of the NAM population (Kiesbauer et al. 2026) | | TSC2 | |
| M2031 | M2031_NAM | | *Lolium multiflorum* | Parent of the NAM population (Kiesbauer et al. 2026) | | TSC2 | |
| M2032 | M2032_NAM | | *Lolium multiflorum* | Parent of the NAM population (Kiesbauer et al. 2026) | | TSC2 | |
| M2033 | M2033_NAM | | *Lolium multiflorum* | Parent of the NAM population (Kiesbauer et al. 2026) | | TSC2 | |
| PSC1 | M-06008/20 | | *Lolium perenne* | Perennial ryegrasss breeding material (Agroscope Reckenholz, Switzerland) | | TSC2 | |
| PSC2 | M-06022/20 | | *Lolium perenne* | Perennial ryegrasss breeding material (Agroscope Reckenholz, Switzerland) | | TSC2 | |
| PSC3 | M-06042/20 | | *Lolium perenne* | Perennial ryegrasss breeding material (Agroscope Reckenholz, Switzerland) | | TSC2 | |
| PC1655-1-1 | PC1655 4xLP Vf1-1 | | *Lolium perenne* | Perennial ryegrasss five-parent polycross (Agroscope Reckenholz, Switzerland) | | TSC2 | |
| PC1655-1-3 | PC1655 4xLP Vf1-3 | | *Lolium perenne* | Perennial ryegrasss five-parent polycross (Agroscope Reckenholz, Switzerland) | | TSC2 | |
| PC1655-1-4 | PC1655 4xLP Vf1-4 | | *Lolium perenne* | Perennial ryegrasss five-parent polycross (Agroscope Reckenholz, Switzerland) | | TSC2 | |
| PC1655-1-5 | PC1655 4xLP Vf1-5 | | *Lolium perenne* | Perennial ryegrasss five-parent polycross (Agroscope Reckenholz, Switzerland) | | TSC2 | |
| PC1655-1-7 | PC1655 4xLP Vf1-7 | | *Lolium perenne* | Perennial ryegrasss five-parent polycross (Agroscope Reckenholz, Switzerland) | | TSC2 | |
| PC1655-1-12 | PC1655 4xLP Vf1-12 | | *Lolium perenne* | Perennial ryegrasss five-parent polycross (Agroscope Reckenholz, Switzerland) | | TSC2 | |
| PC1655-1-13 | PC1655 4xLP Vf1-13 | | *Lolium perenne* | Perennial ryegrasss five-parent polycross (Agroscope Reckenholz, Switzerland) | | TSC2 | |
| PC1655-1-10 | PC1655 4xLP Vf1-10 | | *Lolium perenne* | Perennial ryegrasss five-parent polycross (Agroscope Reckenholz, Switzerland) | | TSC2 | |
| PC1655-1-14 | PC1655 4xLP Vf1-14 | | *Lolium perenne* | Perennial ryegrasss five-parent polycross (Agroscope Reckenholz, Switzerland) | | TSC2 | |
| PC1655-1-15 | PC1655 4xLP Vf1-15 | | *Lolium perenne* | Perennial ryegrasss five-parent polycross (Agroscope Reckenholz, Switzerland) | | TSC2 | |
| PC1655-1-16 | PC1655 4xLP Vf1-16 | | *Lolium perenne* | Perennial ryegrasss five-parent polycross (Agroscope Reckenholz, Switzerland) | | TSC2 | |
| PC1655-1-11 | PC1655 4xLP Vf1-11 | | *Lolium perenne* | Perennial ryegrasss five-parent polycross (Agroscope Reckenholz, Switzerland) | | TSC2 | |
| PC1655-1-18 | PC1655 4xLP Vf1-18 | | *Lolium perenne* | Perennial ryegrasss five-parent polycross (Agroscope Reckenholz, Switzerland) | | TSC2 | |
| PC1541-5-4 | PC1541 2xLp Vf5-4 | | *Lolium perenne* | Perennial ryegrasss four-parent polycross (Agroscope Reckenholz, Switzerland) | | TSC2 | |
| PC1541-5-5 | PC1541 2xLp Vf5-5 | | *Lolium perenne* | Perennial ryegrasss four-parent polycross (Agroscope Reckenholz, Switzerland) | | TSC2 | |
| PC1541-5-6 | PC1541 2xLp Vf5-6 | | *Lolium perenne* | Perennial ryegrasss four-parent polycross (Agroscope Reckenholz, Switzerland) | | TSC2 | |
| PC1541-5-7 | PC1541 2xLp Vf5-7 | | *Lolium perenne* | Perennial ryegrasss four-parent polycross (Agroscope Reckenholz, Switzerland) | | TSC2 | |
| PC1541-5-3 | PC1541 2xLp Vf5-3 | | *Lolium perenne* | Perennial ryegrasss four-parent polycross (Agroscope Reckenholz, Switzerland) | | TSC2 | |
| PC1541-1-1 | PC1541 2xLp Vf1-1 | | *Lolium perenne* | Perennial ryegrasss four-parent polycross (Agroscope Reckenholz, Switzerland) | | TSC2 | |
| PC1541-1-2 | PC1541 2xLp Vf1-2 | | *Lolium perenne* | Perennial ryegrasss four-parent polycross (Agroscope Reckenholz, Switzerland) | | TSC2 | |
| PC1541-1-3 | PC1541 2xLp Vf1-3 | | *Lolium perenne* | Perennial ryegrasss four-parent polycross (Agroscope Reckenholz, Switzerland) | | TSC2 | |
| PC1541-1-4 | PC1541 2xLp Vf1-4 | | *Lolium perenne* | Perennial ryegrasss four-parent polycross (Agroscope Reckenholz, Switzerland) | | TSC2 | |
| PC1541-1-5 | PC1541 2xLp Vf1-5 | | *Lolium perenne* | Perennial ryegrasss four-parent polycross (Agroscope Reckenholz, Switzerland) | | TSC2 | |
| PC1541-1-6 | PC1541 2xLp Vf1-6 | | *Lolium perenne* | Perennial ryegrasss four-parent polycross (Agroscope Reckenholz, Switzerland) | | TSC2 | |
| PC1541-3-13 | PC1541 2xLp Vf3-13 | | *Lolium perenne* | Perennial ryegrasss four-parent polycross (Agroscope Reckenholz, Switzerland) | | TSC2 | |
| PC1541-3-14 | PC1541 2xLp Vf3-14 | | *Lolium perenne* | Perennial ryegrasss four-parent polycross (Agroscope Reckenholz, Switzerland) | | TSC2 | |
| PC1541-1-9 | PC1541 2xLp Vf1-9 | | *Lolium perenne* | Perennial ryegrasss four-parent polycross (Agroscope Reckenholz, Switzerland) | | TSC2 | |
| PC1541-3-15 | PC1541 2xLp Vf3-15 | | *Lolium perenne* | Perennial ryegrasss four-parent polycross (Agroscope Reckenholz, Switzerland) | | TSC2 | |
| PC1541-1-11 | PC1541 2xLp Vf1-11 | | *Lolium perenne* | Perennial ryegrasss four-parent polycross (Agroscope Reckenholz, Switzerland) | | TSC2 | |
| PC1541-3-20 | PC1541 2xLp Vf3-20 | | *Lolium perenne* | Perennial ryegrasss four-parent polycross (Agroscope Reckenholz, Switzerland) | | TSC2 | |
| PC1541-1-13 | PC1541 2xLp Vf1-13 | | *Lolium perenne* | Perennial ryegrasss four-parent polycross (Agroscope Reckenholz, Switzerland) | | TSC2 | |
| PC1541-1-14 | PC1541 2xLp Vf1-14 | | *Lolium perenne* | Perennial ryegrasss four-parent polycross (Agroscope Reckenholz, Switzerland) | | TSC2 | |
| PC1541-1-15 | PC1541 2xLp Vf1-15 | | *Lolium perenne* | Perennial ryegrasss four-parent polycross (Agroscope Reckenholz, Switzerland) | | TSC2 | |
| PC1541-2-1 | PC1541 2xLp Vf2-1 | | *Lolium perenne* | Perennial ryegrasss four-parent polycross (Agroscope Reckenholz, Switzerland) | | TSC2 | |
| PC1541-2-2 | PC1541 2xLp Vf2-2 | | *Lolium perenne* | Perennial ryegrasss four-parent polycross (Agroscope Reckenholz, Switzerland) | | TSC2 | |
| PC1541-3-17 | PC1541 2xLp Vf3-17 | | *Lolium perenne* | Perennial ryegrasss four-parent polycross (Agroscope Reckenholz, Switzerland) | | TSC2 | |
| PC1541-2-4 | PC1541 2xLp Vf2-4 | | *Lolium perenne* | Perennial ryegrasss four-parent polycross (Agroscope Reckenholz, Switzerland) | | TSC2 | |
| PC1541-2-5 | PC1541 2xLp Vf2-5 | | *Lolium perenne* | Perennial ryegrasss four-parent polycross (Agroscope Reckenholz, Switzerland) | | TSC2 | |
| PC1541-2-6 | PC1541 2xLp Vf2-6 | | *Lolium perenne* | Perennial ryegrasss four-parent polycross (Agroscope Reckenholz, Switzerland) | | TSC2 | |
| PC1541-2-7 | PC1541 2xLp Vf2-7 | | *Lolium perenne* | Perennial ryegrasss four-parent polycross (Agroscope Reckenholz, Switzerland) | | TSC2 | |
| PC1541-2-8 | PC1541 2xLp Vf2-8 | | *Lolium perenne* | Perennial ryegrasss four-parent polycross (Agroscope Reckenholz, Switzerland) | | TSC2 | |
| PC1541-2-9 | PC1541 2xLp Vf2-9 | | *Lolium perenne* | Perennial ryegrasss four-parent polycross (Agroscope Reckenholz, Switzerland) | | TSC2 | |
| PC1541-2-10 | PC1541 2xLp Vf2-10 | | *Lolium perenne* | Perennial ryegrasss four-parent polycross (Agroscope Reckenholz, Switzerland) | | TSC2 | |
| PC1541-2-11 | PC1541 2xLp Vf2-11 | | *Lolium perenne* | Perennial ryegrasss four-parent polycross (Agroscope Reckenholz, Switzerland) | | TSC2 | |
| PC1541-2-12 | PC1541 2xLp Vf2-12 | | *Lolium perenne* | Perennial ryegrasss four-parent polycross (Agroscope Reckenholz, Switzerland) | | TSC2 | |
| PC1541-3-18 | PC1541 2xLp Vf3-18 | | *Lolium perenne* | Perennial ryegrasss four-parent polycross (Agroscope Reckenholz, Switzerland) | | TSC2 | |
| PC1541-2-14 | PC1541 2xLp Vf2-14 | | *Lolium perenne* | Perennial ryegrasss four-parent polycross (Agroscope Reckenholz, Switzerland) | | TSC2 | |
| PC1541-3-19 | PC1541 2xLp Vf2-19 | | *Lolium perenne* | Perennial ryegrasss four-parent polycross (Agroscope Reckenholz, Switzerland) | | TSC2 | |
| PC1541-2-16 | PC1541 2xLp Vf2-16 | | *Lolium perenne* | Perennial ryegrasss four-parent polycross (Agroscope Reckenholz, Switzerland) | | TSC2 | |
| PC1541-2-17 | PC1541 2xLp Vf2-17 | | *Lolium perenne* | Perennial ryegrasss four-parent polycross (Agroscope Reckenholz, Switzerland) | | TSC2 | |
| PC1541-5-2 | PC1541 2xLp Vf5-2 | | *Lolium perenne* | Perennial ryegrasss four-parent polycross (Agroscope Reckenholz, Switzerland) | | TSC2 | |
| PC1541-2-19 | PC1541 2xLp Vf2-19 | | *Lolium perenne* | Perennial ryegrasss four-parent polycross (Agroscope Reckenholz, Switzerland) | | TSC2 | |
| PC1541-3-1 | PC1541 2xLp Vf3-1 | | *Lolium perenne* | Perennial ryegrasss four-parent polycross (Agroscope Reckenholz, Switzerland) | | TSC2 | |
| PC1541-3-2 | PC1541 2xLp Vf3-2 | | *Lolium perenne* | Perennial ryegrasss four-parent polycross (Agroscope Reckenholz, Switzerland) | | TSC2 | |
| PC1541-3-3 | PC1541 2xLp Vf3-3 | | *Lolium perenne* | Perennial ryegrasss four-parent polycross (Agroscope Reckenholz, Switzerland) | | TSC2 | |
| PC1541-3-4 | PC1541 2xLp Vf3-4 | | *Lolium perenne* | Perennial ryegrasss four-parent polycross (Agroscope Reckenholz, Switzerland) | | TSC2 | |
| PC1541-3-5 | PC1541 2xLp Vf3-5 | | *Lolium perenne* | Perennial ryegrasss four-parent polycross (Agroscope Reckenholz, Switzerland) | | TSC2 | |
| PC1541-3-6 | PC1541 2xLp Vf3-6 | | *Lolium perenne* | Perennial ryegrasss four-parent polycross (Agroscope Reckenholz, Switzerland) | | TSC2 | |
| PC1541-3-7 | PC1541 2xLp Vf3-7 | | *Lolium perenne* | Perennial ryegrasss four-parent polycross (Agroscope Reckenholz, Switzerland) | | TSC2 | |
| PC1541-3-8 | PC1541 2xLp Vf3-8 | | *Lolium perenne* | Perennial ryegrasss four-parent polycross (Agroscope Reckenholz, Switzerland) | | TSC2 | |
| PC1541-3-9 | PC1541 2xLp Vf3-9 | | *Lolium perenne* | Perennial ryegrasss four-parent polycross (Agroscope Reckenholz, Switzerland) | | TSC2 | |
| PC1541-3-10 | PC1541 2xLp Vf3-10 | | *Lolium perenne* | Perennial ryegrasss four-parent polycross (Agroscope Reckenholz, Switzerland) | | TSC2 | |
| PC1541-3-11 | PC1541 2xLp Vf3-11 | | *Lolium perenne* | Perennial ryegrasss four-parent polycross (Agroscope Reckenholz, Switzerland) | | TSC2 | |
| PC1541-3-12 | PC1541 2xLp Vf3-12 | | *Lolium perenne* | Perennial ryegrasss four-parent polycross (Agroscope Reckenholz, Switzerland) | | TSC2 | |
| Sikem | Sikem | | *Lolium multiflorum* | Sequenced genotype (DLF, Denmark) | | TSC2 | |
| S23 Z | S23 Z | | *Lolium perenne* | Sequenced genotype (Rohner et al. 2023) | | TSC2 | |
| Rabiosa | Rabiosa | | *Lolium multiflorum* | Sequenced genotype (Copetti et a. 2021) | | TSC2 | |
| Kyuss | Kyuss | | *Lolium perenne* | Sequenced genotype (Frei et al. 2022) | | TSC2 | |
| T17 | T17 | | *Lolium perenne* | Sequenced genotype (Nagy et al. 2022) | | TSC2 | |

|  | **Coverage (x)** | **Minimum gene length (bp)** | **Minimum coding sequence length (bp)** | **Minimum protein length (aa)** |
| --- | --- | --- | --- | --- |
| *TIR1* | 10 | 3200 | 1550 | 500 |
| *dsRNAbp* | 10 | 2000 | 1400 | 534 |
| *SDUF247-I* | 10 | 1000 | 1500 | 500 |
| *sS* | 10 | 200 | 200 | 66 |
| *SDUF247-II* | 10 | 1000 | 1500 | 500 |
| *PLP* | 10 | 1300 | 700 | 200 |
| *USP1* | 10 | 1600 | 1500 | 400 |
| *ZDUF247-I* | 10 | 1000 | 1500 | 500 |
| *sZ* | 10 | 200 | 200 | 66 |
| *ZDUF247-II* | 10 | 1000 | 1500 | 580 |
| *GK* | 10 | 1300 | 1200 | 400 |
| *chr2-DUF247* | 10 | 1300 | 1300 | 433 |
| *chr3-DUF247* | 10 | 1200 | 1200 | 400 |
| *chr4-WRKY* | 10 | 660 | 350 | 90 |

Table S6. Overview of filtering parameter applied to each gene.

Table S7. Overview of genotypes represented by the groups in the tanglegram. Each group represents a unique protein sequence per gene; for each gene, the name of the sequences, and hence also the genotype, represented by the group, are listed.

| **Group** | **ZDUF247I** | **sZ** | **ZDUF247II** | **SDUF247I** | **sS** | **SDUF247II** |
| --- | --- | --- | --- | --- | --- | --- |
| Group_1 | Lp_Arara_A_1 | Lp_Arara_A_1 | Lp_Arara_A_1 | Lp_PC1541-1-14_1 | Lp_PC1541-1-13_1 | Lp_PC1541-1-13_2 |
|  | Lp_IOWA3_2 | Lp_IOWA3_2 | Lp_IOWA3_1 | Lp_PC1655-1-13_1 | Lp_PC1541-1-15_1 | Lp_PC1541-1-15_1 |
|  | Lp_IOWA4_1 | Lp_IOWA4_1 | Lp_IOWA4_1 | Lp_PC1655-1-14_3 | Lp_PC1541-2-12_1 | Lp_PC1541-2-12_2 |
|  | Lp_Kyuss_1 | Lp_Kyuss_1 | Lp_Kyuss_1 | Lp_PC1655-1-18_1 | Lp_PC1541-2-1_1 | Lp_PC1541-2-1_1 |
|  | Lm_D45_2 | Lm_D45_1 | Lm_D45_1 |  |  |  |
|  | Lm_M2010_1 | Lm_M2010_2 | Lm_M2010_1 |  |  |  |
|  | Lp_PC1541-1-11_1 | Lp_PC1541-1-11_1 | Lp_PC1541-1-11_1 |  |  |  |
|  | Lp_PC1541-1-13_1 | Lp_PC1541-1-13_1 | Lp_PC1541-1-13_1 |  |  |  |
|  | Lp_PC1541-1-14_2 | Lp_PC1541-1-14_1 | Lp_PC1541-1-14_1 |  |  |  |
|  | Lp_PC1541-2-10_1 | Lp_PC1541-2-10_1 | Lp_PC1541-2-10_1 |  |  |  |
|  | Lp_PC1541-2-17_1 | Lp_PC1541-2-17_1 | Lp_PC1541-2-17_1 |  |  |  |
|  | Lp_PC1541-2-19_1 | Lp_PC1541-2-19_1 | Lp_PC1541-2-19_1 |  |  |  |
|  | Lp_PC1541-2-2_2 | Lp_PC1541-2-2_1 | Lp_PC1541-2-2_1 |  |  |  |
|  | Lp_PC1541-2-7_1 | Lp_PC1541-2-7_1 | Lp_PC1541-2-7_1 |  |  |  |
|  | Lp_PC1541-2-9_1 | Lp_PC1541-2-9_1 | Lp_PC1541-2-9_1 |  |  |  |
|  | Lp_PC1541-3-11_2 | Lp_PC1541-3-11_1 | Lp_PC1541-3-11_1 |  |  |  |
|  | Lp_PC1541-3-13_2 | Lp_PC1541-3-13_1 | Lp_PC1541-3-13_1 |  |  |  |
|  | Lp_PC1541-3-15_1 | Lp_PC1541-3-15_1 | Lp_PC1541-3-15_1 |  |  |  |
|  | Lp_PC1541-3-19_1 | Lp_PC1541-3-19_1 | Lp_PC1541-3-19_1 |  |  |  |
|  | Lp_PC1541-3-20_2 | Lp_PC1541-3-20_1 | Lp_PC1541-3-20_1 |  |  |  |
|  | Lp_PC1541-3-2_1 | Lp_PC1541-3-2_1 | Lp_PC1541-3-2_1 |  |  |  |
|  | Lp_PC1541-3-9_1 | Lp_PC1541-3-9_1 | Lp_PC1541-3-9_1 |  |  |  |
|  | Lp_PC1541-5-2_1 | Lp_PC1541-5-2_1 | Lp_PC1541-5-2_1 |  |  |  |
|  | Lp_PC1541-5-4_1 | Lp_PC1541-5-4_1 | Lp_PC1541-5-4_1 |  |  |  |
|  | Lp_PC1541-5-5_1 | Lp_PC1541-5-5_1 | Lp_PC1541-5-5_1 |  |  |  |
|  | Lp_PC1541-5-6_2 | Lp_PC1541-5-6_1 | Lp_PC1541-5-6_1 |  |  |  |
|  | Lp_PC1655-1-11_3 | Lp_PC1655-1-11_1 | Lp_PC1655-1-11_1 |  |  |  |
| Group_2 | Lp_F1-30_1 | Lp_F1-30_1 | Lp_F1-30_1 | Lp_F1-30_2 | Lp_F1-30_1 | Lp_F1-30_2 |
|  | Lp_F1-39_2 | Lp_F1-39_2 | Lp_F1-39_1 | Lp_F1-39_2 | Lp_F1-39_1 | Lp_F1-39_2 |
|  | Lp_IOWA1_1 | Lp_IOWA1_1 | Lp_IOWA1_1 | Lp_NgB_1 | Lp_NgB_1 | Lp_NgB_1 |
|  | Lp_NgB_1 | Lp_NgB_2 | Lp_NgB_1 | Lp_PC1541-3-13_2 | Lp_PC1541-3-13_2 | Lp_PC1541-3-13_2 |
|  | Lp_PC1541-1-15_1 | Lp_PC1541-1-15_1 | Lp_PC1541-1-15_1 | Lp_PC1541-3-2_2 | Lp_PC1541-3-2_1 | Lp_PC1541-3-2_2 |
|  | Lp_PC1541-2-12_2 | Lp_PC1541-2-12_1 | Lp_PC1541-2-12_1 | Lp_PC1541-3-8_1 | Lp_PC1541-3-8_1 | Lp_PC1541-3-8_2 |
|  | Lp_PC1541-2-14_1 | Lp_PC1541-2-14_1 | Lp_PC1541-2-14_1 | Lp_PC1541-3-9_2 | Lp_PC1541-3-9_1 | Lp_PC1541-3-9_2 |
|  | Lp_PC1541-2-16_1 | Lp_PC1541-2-16_1 | Lp_PC1541-2-16_1 | Lp_PC1541-5-4_1 | Lp_PC1541-5-4_1 | Lp_PC1541-5-4_2 |
|  | Lp_PC1541-3-5_1 | Lp_PC1541-3-5_1 | Lp_PC1541-3-5_1 | Lp_PSC1_2 | Lp_PSC1_1 | Lp_PSC1_2 |
|  | Lp_PC1541-3-6_1 | Lp_PC1541-3-6_1 | Lp_PC1541-3-6_1 | Lp_PSC2_1 | Lp_PSC2_1 | Lp_PSC2_2 |
|  | Lp_PC1541-3-8_1 | Lp_PC1541-3-8_1 | Lp_PC1541-3-8_1 | Lp_PSC3_2 | Lp_PSC3_1 | Lp_PSC3_2 |
|  | Lp_PC1541-5-3_2 | Lp_PC1541-5-3_1 | Lp_PC1541-5-3_1 | Lp_VrnA_1 | Lp_VrnA_1 | Lp_VrnA_1 |
|  | Lp_PC1541-5-7_1 | Lp_PC1541-5-7_1 | Lp_PC1541-5-7_1 |  |  |  |
|  | Lp_PC1655-1-13_1 | Lp_PC1655-1-13_2 | Lp_PC1655-1-13_3 |  |  |  |
|  | Lp_PC1655-1-14_1 | Lp_PC1655-1-14_1 | Lp_PC1655-1-14_1 |  |  |  |
|  | Lp_PC1655-1-18_2 | Lp_PC1655-1-18_1 | Lp_PC1655-1-18_1 |  |  |  |
|  | Lp_PC1655-1-4_1 | Lp_PC1655-1-4_1 | Lp_PC1655-1-4_1 |  |  |  |
|  | Lp_PC1655-1-7_2 | Lp_PC1655-1-7_1 | Lp_PC1655-1-7_1 |  |  |  |
|  | Lp_PSC1_2 | Lp_PSC1_1 | Lp_PSC1_1 |  |  |  |
|  | Lp_PSC2_1 | Lp_PSC2_1 | Lp_PSC2_1 |  |  |  |
|  | Lp_PSC3_1 | Lp_PSC3_1 | Lp_PSC3_1 |  |  |  |
|  | Lp_S23_Z_1 | Lp_S23_Z_2 | Lp_S23_Z_2 |  |  |  |
|  | Lp_S23_Z_repetition_1 | Lp_S23_Z_repetition_2 | Lp_S23_Z_repetition_1 |  |  |  |
|  | Lp_Veyo_1 | Lp_Veyo_2 | Lp_Veyo_1 |  |  |  |
|  | Lp_VrnA_1 | Lp_VrnA_1 | Lp_VrnA_1 |  |  |  |
| Group_3 | Lm_D20_1 | Lm_D20_1 | Lm_D20_1 | Lm_D21_2 | Lm_D21_1 | Lm_D21_2 |
|  | Lm_D49_1 | Lm_D49_1 | Lm_D49_1 | Lm_D22_2 | Lm_D22_1 | Lm_D22_2 |
|  | Lm_M2033_2 | Lm_M2033_2 | Lm_M2033_2 | Lm_D23_2 | Lm_D23_1 | Lm_D23_2 |
|  | Lm_M34_10_CG_1 | Lm_M34_10_CG_2 | Lm_M34_10_CG_2 | Lm_D57_1 | Lm_D57_1 | Lm_D57_1 |
|  | Lp_235_59_150_2 | Lp_235_59_150_1 | Lp_235_59_150_1 | Lp_PC1541-1-1_1 | Lp_PC1541-1-1_1 | Lp_PC1541-1-1_1 |
|  | Lp_S23_Z_2 | Lp_S23_Z_1 | Lp_S23_Z_1 | Lp_PC1541-2-7_2 | Lp_PC1541-2-7_2 | Lp_PC1541-2-7_2 |
|  | Lp_S23_Z_repetition_2 | Lp_S23_Z_repetition_1 | Lp_S23_Z_repetition_2 | Lp_PC1541-3-13_1 | Lp_PC1541-3-13_1 | Lp_PC1541-3-13_1 |
|  |  |  |  | Lp_PC1541-3-15_2 | Lp_PC1541-3-15_1 | Lp_PC1541-3-15_1 |
|  |  |  |  | Lp_PC1541-3-20_2 | Lp_PC1541-3-20_2 | Lp_PC1541-3-20_2 |
|  |  |  |  | Lp_PC1541-3-3_2 | Lp_PC1541-3-3_1 | Lp_PC1541-3-3_1 |
|  |  |  |  | Lp_PC1541-5-5_2 | Lp_PC1541-5-5_1 | Lp_PC1541-5-5_2 |
|  |  |  |  | Lp_PC1655-1-10_3 | Lp_PC1655-1-10_2 | Lp_PC1655-1-10_2 |
|  |  |  |  | Lp_PC1655-1-4_1 | Lp_PC1655-1-4_1 | Lp_PC1655-1-4_1 |
|  |  |  |  | Lp_PC1655-1-5_3 | Lp_PC1655-1-5_3 | Lp_PC1655-1-5_2 |
|  |  |  |  | Lp_PC1655-1-7_4 | Lp_PC1655-1-7_1 | Lp_PC1655-1-7_4 |
|  |  |  |  | Lm_Rabiosa_2 | Lm_Rabiosa_2 | Lm_Rabiosa_1 |
| Group_4 | Lp_Arara_B_1 | Lp_Arara_B_1 | Lp_Arara_B_1 | Lp_IOWA2_2 |  |  |
|  | Lp_IOWA1_2 | Lp_IOWA1_2 | Lp_IOWA1_2 | Lp_Kyuss_1 | Lp_Kyuss_1 | Lp_Kyuss_1 |
|  | Lp_IOWA3_1 | Lp_IOWA3_1 | Lp_IOWA3_2 | Lm_M2020_2 | Lm_M2020_2 | Lm_M2020_1 |
|  | Lm_D54_1 | Lm_D54_1 | Lm_D54_1 | Lp_PC1541-2-11_1 | Lp_PC1541-2-11_2 | Lp_PC1541-2-11_1 |
|  | Lm_D56_2 | Lm_D56_1 | Lm_D56_1 | Lp_PC1541-2-14_2 | Lp_PC1541-2-14_1 | Lp_PC1541-2-14_1 |
|  | Lp_PC1541-1-2_1 | Lp_PC1541-1-2_2 | Lp_PC1541-1-2_1 | Lp_PC1541-2-9_2 | Lp_PC1541-2-9_1 | Lp_PC1541-2-9_1 |
|  | Lp_PC1541-1-4_2 | Lp_PC1541-1-4_1 | Lp_PC1541-1-4_1 | Lp_PC1541-3-11_2 | Lp_PC1541-3-11_1 | Lp_PC1541-3-11_1 |
|  | Lp_PC1655-1-13_3 | Lp_PC1655-1-13_1 | Lp_PC1655-1-13_2 | Lp_PC1541-3-19_1 | Lp_PC1541-3-19_1 | Lp_PC1541-3-18_1 |
|  | Lp_PC1655-1-14_2 | Lp_PC1655-1-14_2 | Lp_PC1655-1-14_2 | Lp_PC1655-1-11_1 | Lp_PC1655-1-11_2 | Lp_PC1655-1-11_1 |
|  | Lp_PC1655-1-1_1 | Lp_PC1655-1-1_2 | Lp_PC1655-1-1_1 | Lp_PC1655-1-13_3 | Lp_PC1655-1-13_2 | Lp_PC1655-1-13_1 |
|  |  |  |  | Lp_PC1655-1-14_1 | Lp_PC1655-1-14_2 | Lp_PC1655-1-14_2 |
|  |  |  |  | Lp_PC1655-1-16_1 | Lp_PC1655-1-16_2 | Lp_PC1655-1-16_2 |
|  |  |  |  | Lp_PC1655-1-18_2 | Lp_PC1655-1-18_2 | Lp_PC1655-1-18_3 |
|  |  |  |  | Lp_PC1655-1-7_3 | Lp_PC1655-1-7_2 | Lp_PC1655-1-7_2 |
|  |  |  |  | Lp_restorer_CMS_2 | Lp_restorer_CMS_1 | Lp_restorer_CMS_1 |
| Group_5 | Lm_D27_1 | Lm_D27_1 | Lm_D27_2 | Lm_121_2 | Lm_121_1 | Lm_121_2 |
|  | Lm_D30_2 | Lm_D30_2 | Lm_D30_2 | Lm_27_2 | Lm_27_1 | Lm_27_1 |
|  | Lm_D42_1 | Lm_D42_1 | Lm_D42_1 | Lm_81_2 | Lm_81_2 | Lm_81_1 |
|  | Lm_D51_2 | Lm_D51_1 | Lm_D51_1 | Lm_84_2 | Lm_84_1 | Lm_84_2 |
|  | Lm_M2027_1 | Lm_M2027_1 | Lm_M2027_1 | Lm_99_2 | Lm_99_1 | Lm_99_2 |
|  | Lm_M2030_1 | Lm_M2030_1 | Lm_M2030_1 | Lm_D54_2 | Lm_D54_1 | Lm_D54_1 |
|  | Lp_PC1541-1-9_1 | Lp_PC1541-1-9_1 | Lp_PC1541-1-9_2 | Lm_M2006_3 | Lm_M2006_2 | Lm_M2006_4 |
|  | Lm_Rabiosa_1 | Lm_Rabiosa_1 | Lm_Rabiosa_2 |  |  |  |
| Group_6 | Lp_F1-39_2 | Lp_F1-39_1 | Lp_F1-39_2 | Lm_112_2 | Lm_112_1 | Lm_112_1 |
|  | Lp_PC1541-2-11_2 | Lp_PC1541-2-11_1 | Lp_PC1541-2-11_1 | Lm_125_1 | Lm_125_1 | Lm_125_2 |
|  | Lp_PC1655-1-10_2 | Lp_PC1655-1-10_1 | Lp_PC1655-1-10_1 | Lm_13_1 | Lm_13_1 | Lm_13_2 |
|  | Lp_T17_1 | Lp_T17_1 | Lp_T17_1 | Lm_16_1 | Lm_16_2 | Lm_16_2 |
|  | Lp_Veyo_2 | Lp_Veyo_1 | Lp_Veyo_2 | Lm_38_1 | Lm_38_1 | Lm_38_1 |
|  |  |  |  | Lm_41_1 | Lm_41_1 | Lm_41_2 |
|  |  |  |  | Lm_73_1 | Lm_73_1 | Lm_73_2 |
|  |  |  |  | Lm_D11_2 | Lm_D11_1 | Lm_D11_1 |
|  |  |  |  | Lm_D1_1 | Lm_D1_1 | Lm_D1_2 |
|  |  |  |  | Lm_D21_1 | Lm_D21_2 | Lm_D21_1 |
|  |  |  |  | Lm_D2_2 | Lm_D2_1 | Lm_D2_1 |
|  |  |  |  | Lm_D36_2 | Lm_D36_1 | Lm_D36_2 |
|  |  |  |  | Lm_D42_1 | Lm_D42_1 | Lm_D42_2 |
|  |  |  |  | Lm_M2026_2 | Lm_M2026_1 | Lm_M2026_1 |
| Group_7 | Lm_D37_2 | Lm_D37_1 | Lm_D37_1 | Lp_IOWA1_2 | Lp_IOWA1_1 | Lp_IOWA1_1 |
|  | Lm_D57_1 | Lm_D57_1 | Lm_D57_1 | Lp_IOWA4_2 | Lp_IOWA4_1 | Lp_IOWA4_1 |
|  | Lm_M2018_1 | Lm_M2018_1 | Lm_M2018_1 | Lp_PC1541-1-11_2 | Lp_PC1541-1-11_2 | Lp_PC1541-1-11_1 |
|  | Lm_M2026_2 | Lm_M2026_1 | Lm_M2026_1 | Lp_PC1541-1-13_1 | Lp_PC1541-1-13_2 | Lp_PC1541-1-13_1 |
|  | Lp_PC1541-1-1_2 | Lp_PC1541-1-1_1 | Lp_PC1541-1-1_1 | Lp_PC1541-2-16_2 | Lp_PC1541-2-16_2 | Lp_PC1541-2-6_1 |
|  |  |  |  | Lp_PC1541-2-6_1 | Lp_PC1541-2-6_1 | Lp_PC1541-2-7_1 |
|  |  |  |  | Lp_PC1541-2-7_1 | Lp_PC1541-2-7_1 | Lp_PC1541-2-8_1 |
|  |  |  |  | Lp_PC1541-2-8_2 | Lp_PC1541-2-8_2 | Lp_PC1541-3-12_2 |
|  |  |  |  | Lp_PC1541-3-12_2 | Lp_PC1541-3-12_2 | Lp_PC1541-3-17_1 |
|  |  |  |  | Lp_PC1541-3-17_2 | Lp_PC1541-3-17_1 | Lp_PC1541-3-1_2 |
|  |  |  |  | Lp_PC1541-3-18_1 | Lp_PC1541-3-1_2 | Lp_PC1541-3-6_1 |
|  |  |  |  | Lp_PC1541-3-1_2 | Lp_PC1541-3-6_2 | Lp_PC1541-3-8_1 |
|  |  |  |  | Lp_PC1541-3-6_2 | Lp_PC1541-3-8_2 | Lp_PC1541-3-9_1 |
|  |  |  |  | Lp_PC1541-3-8_2 | Lp_PC1541-3-9_2 | Lp_PC1541-5-6_1 |
| Group_8 | Lm_D16_1 | Lm_D16_1 | Lm_D16_1 | Lm_D27_1 | Lm_D27_1 | Lm_D27_1 |
|  | Lm_D24_2 | Lm_D24_1 | Lm_D24_2 | Lm_D34_1 | Lm_D34_2 | Lm_D34_2 |
|  | Lm_D33_1 | Lm_D33_1 | Lm_D33_1 | Lm_D51_1 | Lm_D51_1 | Lm_D51_1 |
|  | Lm_D34_1 | Lm_D34_1 | Lm_D34_2 | Lp_PC1541-1-9_1 | Lp_PC1541-1-9_1 | Lp_PC1541-1-9_1 |
| Group_9 | Lp_Arara_B_2 | Lp_Arara_B_2 | Lp_Arara_B_2 | Lm_125_2 | Lm_125_2 | Lm_125_1 |
|  | Lm_D35_2 | Lm_D35_2 | Lm_D35_1 | Lm_16_2 | Lm_16_1 | Lm_16_1 |
|  | Lm_Rabiosa_2 | Lm_Rabiosa_2 | Lm_Rabiosa_1 | Lm_38_2 | Lm_38_2 | Lm_38_2 |
|  |  |  |  | Lm_73_2 | Lm_73_2 | Lm_73_1 |
|  |  |  |  | Lm_81_1 | Lm_81_1 | Lm_81_2 |
|  |  |  |  | Lm_D42_2 | Lm_D42_2 | Lm_D42_1 |
|  |  |  |  | Lm_D48_1 | Lm_D48_1 | Lm_D48_1 |
|  |  |  |  | Lm_M2004_2 | Lm_M2004_1 | Lm_M2004_1 |
|  |  |  |  | Lp_PC1541-1-5_2 | Lp_PC1541-1-5_1 | Lp_PC1541-1-5_1 |
|  |  |  |  | Lp_PC1541-1-6_2 | Lp_PC1541-1-6_1 | Lp_PC1541-1-6_1 |
| Group_10 | Dg_1_2 | Dg_1_1 | Dg_1_2 | Lm_D34_2 | Lm_D34_1 | Lm_D34_1 |
|  | Dg_2_1 | Dg_2_1 | Dg_2_1 | Lm_D8_1 | Lm_D8_1 | Lm_D8_1 |
|  | Dg_4_2 | Dg_4_1 | Dg_4_1 | Lm_M2002_2 | Lm_M2002_1 | Lm_M2002_1 |
|  | Dg_5_2 | Dg_5_1 | Dg_5_1 | Lm_M2009_2 | Lm_M2009_2 | Lm_M2009_2 |
|  |  |  |  | Lm_M2017_2 | Lm_M2017_1 | Lm_M2017_2 |
|  |  |  |  | Lm_M2021_1 | Lm_M2021_1 | Lm_M2021_1 |
|  |  |  |  | Lm_M2028_1 | Lm_M2028_2 | Lm_M2028_1 |
|  |  |  |  | Lp_PC1541-1-6_1 | Lp_PC1541-1-6_2 | Lp_PC1541-1-6_2 |
|  |  |  |  | Lm_D56_2 | Lm_D56_1 | Lm_D56_2 |
| Group_11 | Lm_D30_1 | Lm_D30_1 | Lm_D30_1 | Lm_M2002_1 | Lm_M2002_2 | Lm_M2002_2 |
|  | Lp_PC1655-1-11_4 | Lp_PC1655-1-11_2 | Lp_PC1655-1-11_2 | Lm_M2003_2 | Lm_M2003_2 | Lm_M2003_1 |
|  | Lp_PC1655-1-7_1 | Lp_PC1655-1-7_2 | Lp_PC1655-1-7_2 | Lm_M2019_2 | Lm_M2019_1 | Lm_M2019_1 |
|  |  |  |  | Lm_M2028_2 | Lm_M2028_1 | Lm_M2028_2 |
|  |  |  |  | Lp_PC1541-1-2_2 | Lp_PC1541-1-2_1 | Lp_PC1541-1-2_2 |
|  |  |  |  | Lp_PC1655-1-3_2 | Lp_PC1655-1-3_1 | Lp_PC1655-1-3_2 |
| Group_12 |  |  |  | Lp_IOWA6_2 | Lp_IOWA6_1 | Lp_IOWA6_1 |
|  |  |  |  | Lm_D18_2 | Lm_D18_1 | Lm_D18_1 |
|  |  |  |  | Lm_D24_1 | Lm_D24_2 | Lm_D24_1 |
|  |  |  |  | Lm_M2006_1 | Lm_M2006_1 | Lm_M2006_1 |
|  |  |  |  | Lm_M2015_2 | Lm_M2015_1 | Lm_M2015_1 |
|  |  |  |  | Lm_M2033_1 | Lm_M2033_2 | Lm_M2033_1 |
| Group_13 |  |  |  | Lm_D12_2 | Lm_D12_1 | Lm_D12_1 |
|  |  |  |  | Lm_D13_2 | Lm_D13_1 | Lm_D13_2 |
|  |  |  |  | Lm_D49_1 | Lm_D49_1 | Lm_D49_2 |
|  |  |  |  | Lm_M2033_2 | Lm_M2033_1 | Lm_M2033_2 |

### References

Chen, Yutang, Dario Copetti, Roland Kölliker, et al. 2025. ‘Chromosome-Level Haplotype-Resolved Assembly of Highly Heterozygous Grass Genomes with PhaseGrass’. *Nature Communications* 17 (1): 12. https://doi.org/10.1038/s41467-025-66377-5.

Copetti, Dario, Steven A. Yates, Maximilian M. Vogt, et al. 2021. *Evidence for High Intergenic Sequence Variation in Heterozygous Italian Ryegrasss (* Lolium Multiflorum *Lam.) Genome Revealed by a High-Quality Draft Diploid Genome Assembly*. Preprint. Genomics. https://doi.org/10.1101/2021.05.05.442707.

Cropano, Claudio, Iain Place, Chloé Manzanares, et al. 2021. ‘Characterization and Practical Use of Self-Compatibility in Outcrossing Grass Species’. *Annals of Botany* 127: 841–52. https://doi.org/10.1093/aob/mcab043.

Frei, Daniel, Elisabeth Veekman, Daniel Grogg, et al. 2021. ‘Ultralong Oxford Nanopore Reads Enable the Development of a Reference-Grade Perennial Ryegrasss Genome Assembly’. *Genome Biology and Evolution* 13 (8): 1–6. https://doi.org/10.1093/gbe/evab159.

Goettelmann, Florian, Ralf Koebnik, Veronica Roman-Reyna, Bruno Studer, and Roland Kölliker. 2023. ‘High Genomic Plasticity and Unique Features of Xanthomonas Translucens Pv. Graminis Revealed through Comparative Analysis of Complete Genome Sequences.’ *BMC Genomics* (England) 24 (1): 741. https://doi.org/10.1186/s12864-023-09855-8.

Honig, Josh A., Ehud Zelzion, Nicole E. Wagner, et al. 2017. ‘Microsatellite Identification in Perennial Ryegrasss Using Next-Generation Sequencing’. *Crop Science* 57 (S1): S-331. https://doi.org/10.2135/cropsci2016.07.0608.

Huang, Linkai, Guangyan Feng, Haidong Yan, et al. 2020. ‘Genome Assembly Provides Insights into the Genome Evolution and Flowering Regulation of Orchardgrass’. *Plant Biotechnology Journal* 18 (2): 373–88. https://doi.org/10.1111/pbi.13205.

Kiesbauer, Jenny, Christoph Grieder, Meril Sindelar, et al. 2026. ‘The First Nested Association Mapping (NAM) Population for Italian Ryegrasss Reveals Genomic Regions Associated with Seed Shattering’. *BMC Plant Biology*, ahead of print, September 12. https://doi.org/10.1186/s12870-026-09936-2.

Knorst, Verena, Steven Yates, Stephen Byrne, et al. 2019. ‘First Assembly of the Gene‐space of *Lolium Multiflorum* and Comparison to Other Poaceae Genomes’. *Grassland Science* 65 (2): 125–34. https://doi.org/10.1111/grs.12225.

Kopecký, David, Jan Bartos, Adam J. Lukaszewski, et al. 2009. ‘Development and Mapping of DArT Markers within the Festuca - Lolium Complex.’ *BMC Genomics* (England) 10 (October): 473. https://doi.org/10.1186/1471-2164-10-473.

Manzanares, Chloé, Susanne Barth, Daniel Thorogood, et al. 2016. ‘A Gene Encoding a DUF247 Domain Protein Cosegregates with the *S* Self-Incompatibility Locus in *Perennial Ryegrasss*’. *Molecular Biology and Evolution* 33 (4): 870–84. https://doi.org/10.1093/molbev/msv335.

Nagy, Istvan, Elisabeth Veeckman, Chang Liu, et al. 2022. ‘Chromosome-Scale Assembly and Annotation of the Perennial Ryegrasss Genome’. *BMC Genomics* 23 (1): 505. https://doi.org/10.1186/s12864-022-08697-0.

Paril, Jefferson, Gunjan Pandey, Emma M. Barnett, et al. 2022. ‘Rounding up the Annual Ryegrasss Genome: High-Quality Reference Genome of Lolium Rigidum’. *Frontiers in Genetics* 13 (Kloot 1983). https://doi.org/10.3389/fgene.2022.1012694.

Rohner, Marius, Chloé Manzanares, Steven Yates, et al. 2023. ‘Fine-Mapping and Comparative Genomic Analysis Reveal the Gene Composition at the S and Z Self-Incompatibility Loci in Grasses’. *Molecular Biology and Evolution* 40 (1): msac259. https://doi.org/10.1093/molbev/msac259.

Samy, Jeevan Karloss Antony, Odd Arne Rognli, and Mallikarjuna Rao Kovi. 2020. ‘ForageGrassBase: Molecular Resource for the Forage Grass Meadow Fescue (Festuca Pratensis Huds.)’. *Database* 2020 (January): baaa046. https://doi.org/10.1093/database/baaa046.

Thorogood, Daniel, Steven Yates, Chloé Manzanares, et al. 2017. ‘A Novel Multivariate Approach to Phenotyping and Association Mapping of Multi-Locus Gametophytic Self-Incompatibility Reveals S, Z, and Other Loci in a Perennial Ryegrasss (Poaceae) Population’. *Frontiers in Plant Science* 8 (August). https://doi.org/10.3389/FPLS.2017.01331/ABSTRACT.

Veeckman, Elisabeth, Sabine Van Glabeke, Annelies Haegeman, et al. 2019. ‘Overcoming Challenges in Variant Calling: Exploring Sequence Diversity in Candidate Genes for Plant Development in Perennial Ryegrasss (Lolium Perenne)’. *DNA Research* 26 (1): 1–12. https://doi.org/10.1093/dnares/dsy033.

Velmurugan, Janaki, Ewan Mollison, Susanne Barth, et al. 2016. ‘An Ultra-High Density Genetic Linkage Map of Perennial Ryegrasss (Lolium Perenne) Using Genotyping by Sequencing (GBS) Based on a Reference Shotgun Genome Assembly’. *Annals of Botany* 118 (1): 71–87. https://doi.org/10.1093/aob/mcw081.

Vogt, Maximilian, Steven Yates, Timothy Sykes, Wilbert Luesink, Michael Koch, and Bruno Studer. 2020. ‘Developing Heterotic Groups for Successful Hybrid Breeding in Perennial Ryegrasss’. *Agronomy* 10 (9): 1410. https://doi.org/10.3390/agronomy10091410.
